# Treatment of wound infections in a mouse model using Zn^2+^-releasing phage bound to gold nanorods

**DOI:** 10.1101/2022.01.05.475129

**Authors:** Huan Peng, Daniele Rossetto, Sheref S. Mansy, Maria C. Jordan, Kenneth P. Roos, Irene A. Chen

## Abstract

Infections caused by drug-resistant bacteria, particularly gram-negative organisms, are increasingly difficult to treat using antibiotics. A potential alternative is ‘phage therapy’, in which phages infect and lyse the bacterial host. However, phage therapy poses serious drawbacks and safety concerns, such as the risk of genetic transduction of antibiotic resistance genes, inconsistent pharmacokinetics, and unknown evolutionary potential. In contrast, metallic nanoparticles possess precise, tunable properties, including efficient conversion of electronic excitation into heat. In this work, we demonstrate that engineered phage-nanomaterial conjugates that target the gram-negative pathogen *P. aeruginosa*, are highly effective as a treatment of infected wounds in mice. Photothermal heating, performed as a single treatment (15 min) or as two treatments on consecutive days, rapidly reduced the bacterial load and released Zn^2+^ to promote wound healing. The phage-nanomaterial treatment was significantly more effective than systemic fluoroquinolone antibiotics in reducing both bacterial load and wound size, and was notably effective against a *P. aeruginosa* strain resistant to polymyxins, a last-line antibiotic therapy. Unlike these antibiotics, the phage-nanomaterial showed no detectable toxicity or systemic effects in mice, consistent with the short duration and localized nature of phage- nanomaterial treatment. Our results demonstrate that phage therapy controlled by inorganic nanomaterials can be a safe and effective antimicrobial strategy *in vivo*.

## Introduction

Infections caused by multidrug-resistant bacteria pose an increasing threat to modern medicine ^1–3^. Multidrug-resistant organisms presently cause >700,000 deaths annually, and this number is projected to reach 10 million, exceeding the number of deaths from cancer, by 2050 ^4^. The economic burden associated with drug-resistant pathogens is also high, estimated at $133 billion annually in the U.S. alone ^5^. In addition to improved antibiotic stewardship, the development of alternative antimicrobial classes, particularly for multidrug-resistant gram-negative organisms, is a crucial element of the response to this problem ^4^. One potential alternative antimicrobial agent is phages ^6^, but phage therapy, as traditionally envisioned, also poses serious disadvantages. Naturally occurring phages can carry toxin genes (e.g., cholera toxin ^7^), transfer antibiotic resistance among pathogens (e.g., among *Staphylococcus aureus* ^8^), and physically support a pathological biofilm ^9–10^, causing biosafety concerns. Furthermore, phage self-replication causes unusual or unpredictable pharmacokinetics and pharmacodynamics, and rapid cell lysis can result in unwanted release of bacterial endotoxins ^11–12^.

At the same time, engineered phages are increasingly exploited for a variety of applications ^13^. For therapeutic aims, one possible approach to mitigate the problems of phage therapy is to use a non-lytic phage for bacterial attachment, while relying on a conjugated nanomaterial, specifically gold nanorods (AuNRs), to destroy the bacteria as well as the phage in a controlled manner. Gold nanorods are excited by light, leading to coherent electronic oscillation (surface plasmon resonance), whose energy is released as heat. This photothermal property has been used therapeutically to kill malignant cells^14–15^, particularly when the nanomaterial is tuned to absorb in near-infrared (NIR) wavelengths, which can penetrate a few centimeters into the skin ^16–18^. M13 is a well-studied, non-lytic, filamentous *E. coli* phage that is widely used as a platform for phage display ^19–20^. The coat of M13 is primarily composed of ∼2700 copies of g8p protein, in addition to several copies of the receptor-binding protein (g3p). Each g8p possesses multiple solvent-accessible carboxylates in addition to an exposed N-terminus, allowing chemical conjugation ^21^. M13 can be rationally designed to attach to different gram-negative pathogens by swapping the receptor-binding domain of g3p ^22–23^. Phage-AuNR bioconjugates (‘phanorods’) have been shown to be effective against bacterial cells *in vitro,* through the photothermal effect ^24^. However, whether the *in vitro* properties translate to *in vivo* efficacy as an antimicrobial treatment is unknown. Potential issues with phanorod treatment *in vivo* include: inactivation or clearance of phanorods by the immune system; inhibition of the antimicrobial effect by a biofilm; inhibition of receptor binding by the physiological medium; possible toxicity of phanorods to the animal; or insufficient selectivity causing thermal damage to animal tissue.

Wounds represent an important medical burden, estimated at 2-4% of total health care costs ^25^. Multidrug-resistant *Pseudomonas aeruginosa*, a gram-negative microorganism, is a particular concern in chronic and acute wound infections, including burns, especially in nosocomial settings ^26–29^. Indeed, *P. aeruginosa* has been designated as a critical Priority 1 bacterial pathogen by the World Health Organization as well as a ‘serious threat’ microbe by the Centers for Disease Control and Prevention ^5, 30^. Multiple murine models of *P. aeruginosa* wound infections have been described, including for open wounds ^31^. Although bioavailability can be a concern for phages, wounds are anatomically accessible for direct topical application of phage-based reagents ^32^, and wounds can also be readily exposed to light for photothermal activation of AuNRs. A chimeric phage, M13-g3p(Pf1), was previously engineered to bind to the type IV pili of *P. aeruginosa*, a virulence factor involved in both twitching motility and attachment sensing, which has been recently suggested as a therapeutic target ^33–36^.

In this work, we prepared a new material, phanorod-Zn, based on phanorods that were synthesized by conjugating M13-g3p(Pf1) to gold nanorods absorbing in the NIR wavelengths. The design of phanorod-Zn comprises three modular modifications to the phage. First, tight binding to the targeted host cell (*P. aeruginosa*) is accomplished through modification of the receptor-binding domain, namely the N-terminal domain of the minor coat protein g3p, of the phage. For the phanorod-Zn described in this work, the receptor-binding domain of M13, which attaches to the F pilus of *E. coli*, has been swapped for the receptor-binding domain of phage Pf1, which attaches to the type IV pili of *P. aeruginosa*. This modification directs the chimeric phage to bind the receptor pili of *P. aeruginosa* ^22, 37^. Second, although the chimeric phages are not expected to replicate on the new host, cell-killing activity is conferred through conjugation of a photothermal agent, AuNRs, to the chimeric phage. Since the phages are approximately 1 micron in length, each virion is able to carry multiple AuNRs, which are approximately 50 nanometers in length^37^. We have previously shown that such phanorod (phage-AuNR) conjugates have excellent cell-killing properties *in vitro* ^37^, although the efficacy *in vivo* was not previously studied. Third, for the *in vivo* application studied here, we further loaded the chimeric phages with a zinc-binding peptide (Pol-K, described below), in order to deliver Zn^2+^ to the bacterial infection. The motivation behind this third modification is two-fold. Zinc (Zn^2+^) has been observed to both promote wound healing ^38–39^ and to inhibit bacterial growth, likely through multiple molecular mechanisms. Therefore, Zn^2+^ release would be a desirable feature for a material treating wound infections. We decorated the phanorods with a zinc-binding peptide (Pol-K) designed to release Zn^2+^ upon photothermal heating (phanorod-Zn). The dodecapeptide Pol-K represents a minimal Zn^2+^-binding motif, based on a bioinformatic analysis of Zn^2+^- binding protein sequences.

In this design, Pol-K acts as a carrier for Zn^2+^, and the phage acts as a targeting agent to bring both the AuNRs as well as bound Zn^2+^ to the bacterial cells. Phages have been shown to possess advantages over antibodies as targeting agents, having both greater stability against environmental factors as well as greater cell-killing efficiency ^37^. Photothermal activation of the AuNRs should then cause localized heating that kills the bacterial cells, and photothermal release of Zn^2+^ should potentiate the antibacterial activity and promote wound healing. Using a mouse model of *P. aeruginosa*-infected wounds, we assessed the efficacy and toxicity of phanorod and phanorod-Zn treatment (Scheme 1). Our experimental results show that phanorod and phanorod- Zn treatments were highly effective, yielding faster wound healing compared to standard antibiotic treatments (e.g., ciprofloxacin) or antibody-conjugated AuNRs. Phanorod-Zn treatment was also effective when administered late into infection, and prevented death in cases of otherwise terminal wounds. Wounds infected by a strain of *P. aeruginosa* that was resistant to last-line antibiotic therapy (polymyxins) were also effectively treated by phanorod-Zn. Serum biomarkers and histological analysis did not show systemic toxicity of phanorods or phanorod- Zn to the mice, in contrast to the fluoroquinolone and polymyxin antibiotics. The results indicate that phage-based nanomaterials, such as phanorods and phanorod-Zn, may be a promising alternative antimicrobial strategy for treatment of multidrug-resistant bacterial infections of wounds.

**Scheme 1.**
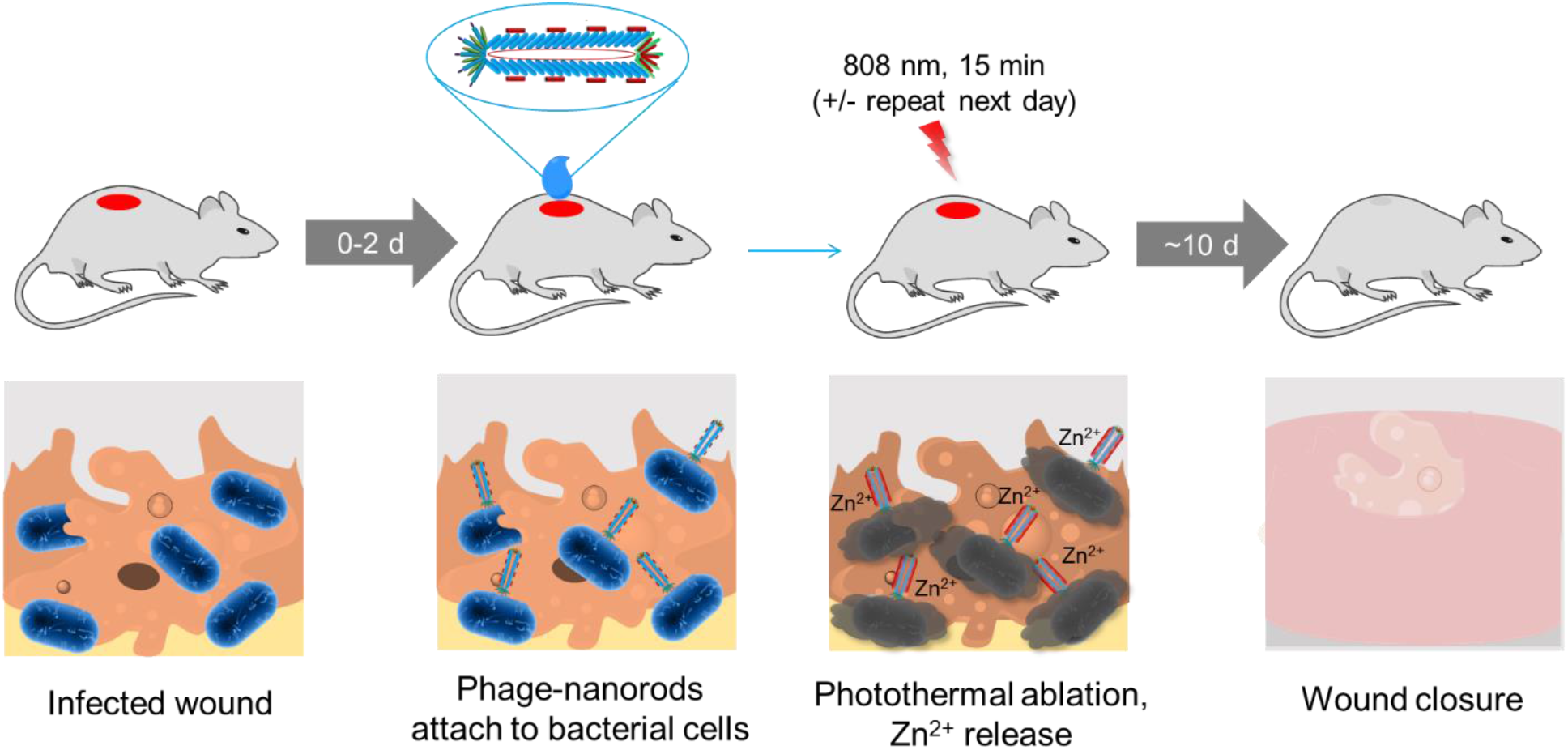
Treatment of a *P. aeruginosa*-infected wound by phage-conjugated gold nanorods decorated with Zn^2+^. Treatment is initiated 0-2 days after wound inoculation. The phage-based reagent is applied to the wound and allowed to bind for 30 min. The wound is irradiated with near-infrared light for 15 min to trigger photothermal ablation of *P. aeruginosa* and release of Zn^2+^; irradiation may be repeated the next day. Wound healing is observed over the next ∼10 days.

## Results

### Synthesis of phanorod-Zn and loading of Zn^2+^

Phage-AuNRs (‘phanorods’) were synthesized as reported previously ^24^. The phage used in this work, M13-g3p(Pf1), was engineered as a chimera of M13 and the *P. aeruginosa* phage Pf1. In M13-g3p(Pf1), the receptor-binding domain of M13 (N-terminal domain of g3p) was replaced by the corresponding domain from Pf1, thereby enabling the chimera M13-g3p(Pf1) to attach to *P. aeruginosa* via type IV pili. The gold nanorods (average length, 53.2 nm; average width, 13.7 nm) were thiolated, conjugated to phages, and passivated by HOOC-PEG-SH.

Phanorod-Zn was prepared by conjugation of the synthetic Zn^2+^-binding peptide Pol-K (amino acid sequence: GCFCEDACDKCG; *K_d,Zn_* for Zn^2+^ = 41 μM; Figure S1a-d) with phanorods through EDC (1-ethyl-3-(3-dimethylaminopropyl)carbodiimide) chemistry. Available carboxylic acid groups for conjugation include three solvent-accessible residues (Glu2, Asp4, and Asp5) per copy of the major coat protein, g8p (of which there are ∼2700 copies per particle) ^40^, as well as –COOH groups on the surface of gold nanorods. The Fourier transform infrared (FTIR) spectra of phanorods, Pol-K, and phanorod-Zn showed that phanorod-Zn exhibited increased signals at 1510 and 3036 cm^-1^ (matching the C-C and C-H stretching, respectively, of the phenyl group of Pol-K), and at 2550 cm^-1^ (from the S-H stretching of the thiol groups), indicating successful peptide conjugation (Figure S1e). Conjugated Pol-K on phanorod-Zn was quantified with Ellman’s assay (measuring added thiol groups compared to phanorods) and the phage particles were quantified by real-time PCR, allowing estimation that each phanorod-Zn carries 1769 ± 357 Pol-K peptides (mean +/- standard deviation, n = 3 samples). Finally, Zn^2+^ was loaded by overnight incubation with ZnCl_2_ followed by washing.

The loading capacity of phanorod-Zn for Zn^2+^ was measured by quantifying Zn^2+^ by inductively coupled plasma mass spectrometry (ICP-MS). While phanorods (without Pol-K) did not bind detectable Zn^2+^ (i.e., < 0.002 ng/mL), phanorod-Zn were found to be loaded by 1410 ± 672 Zn^2+^ ions per phanorod-Zn particle, close to the expectation based on a 1:1 stoichiometry with the amount of Pol-K as measured by Ellman’s assay. Thus, this bioconjugate contained approximately one Pol-K copy for every two copies of g8p, and each Pol-K copy was bound to ∼0.8 Zn^2+^ ion on average.

Additionally, the binding affinity of phanorods and phanorod-Zn to host cells was measured to assess the strength of attachment to host cells. The phage-based reagents were mixed with host cells, centrifuged, and washed, and the amount bound was determined by quantitative PCR (Supporting Methods). The dissociation constants (*K_d_*) of M13-g3p(Pf1), phanorods, and phanorod-Zn for the host cell *P. aeruginosa* (ATCC25102 (Schroeter) Migula) were found to be similar to each other (5-6 pM; Figure S2a), with affinities similar to that seen for the interaction between M13 and *E. coli* (2 pM^41^). This result indicates that bioconjugation and modification with Pol-K did not substantially perturb the binding affinity to host cells.

### Release of Zn^2+^ from phanorod-Zn

The Pol-K dodecapeptide was designed based on the compact Zn^2+^-binding motif of RNA polymerase II and contains four cysteine ligands that tetrahedrally coordinate Zn^2+^. We hypothesized that release of Zn^2+^ would occur at high temperatures, due to an increase in the off-rate of the bound Zn^2+^, structural perturbations to Pol- K, or both ^42–43^. Laser irradiation of AuNRs leads to localized plasmonic heating and temperature increase (characterized further below). We therefore irradiated phanorod-Zn in PBS and monitored release of Zn^2+^ by inductively coupled plasma mass spectrometry (ICP-MS). After irradiation for 15 min, the bulk temperature reached 55°C, and the cumulative Zn^2+^ ion release reached ∼1.06 ppm (17 μM) after 1 day (Figure 1a). Additional laser irradiation on day 2 (15 min per day, again giving a bulk temperature increase to 55°C) resulted in additional release of Zn^2+^, yielding a concentration of 1.69 ppm (26 μM). Laser irradiation was again applied on day 3, but no significant additional release of Zn^2+^ was detected on day 3 (Figure 1b), indicating that maximal release was achieved after 2 days. When no laser was applied, <0.2 ppm of Zn^2+^ was released after three days of incubation in PBS (Figure 1). Phanorods (in the absence of conjugation to Pol-K) did not capture detectable Zn^2+^ ions as assessed by ICP-MS, indicating that the capture and release of Zn^2+^ ions were attributable to the conjugated Pol-K peptide.

**Figure 1.**
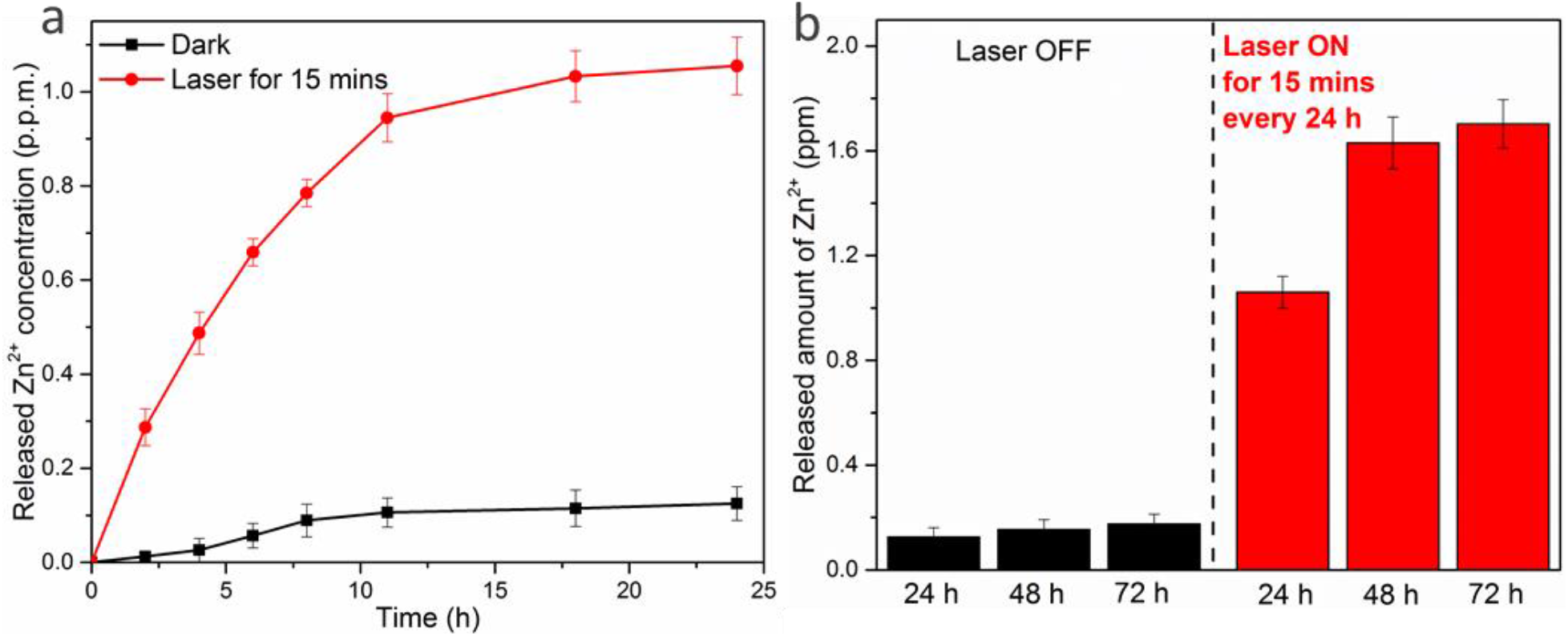
Release of Zn^2+^ ions by phanorod-Zn, measured by ICP-MS. Cumulative Zn^2+^ release curves of phanorod-Zn (10^14^ phage per mL) in PBS buffer after a single 15 min irradiation (a), or with daily irradiation (15 min/day for 3 days) (b). The amount of Zn^2+^ loaded in phanorod-Zn is 15.29 ± 0.73 ppm before irradiation. Error bars indicate mean ± standard deviation (n = 3 independent samples).

To determine whether Zn^2+^ release could be due to higher temperatures alone, we measured the effect of bulk temperature, without laser irradiation, on the rate of ion release. Although Zn^2+^ is spectroscopically silent, Co^2+^ is a well-established substitute of Zn^2+^ for spectroscopic study of Zn^2+^-binding proteins ^44^. Using this system, ligand release from Pol-K was observed to be negligible at room temperature, but increased with increasing bulk temperature, in the absence of irradiation (Figure S1d). Thus, heating by itself accelerates the rate of metal ligand release. We further tested the dependence of Zn^2+^ release on photothermal heating at different levels in the context of phanorod-Zn. We exposed phanorod-Zn to different intensities of laser irradiation, resulting in a range of different bulk temperatures, and quantified Zn^2+^ release by ICP-MS. As expected, while Zn^2+^ release was negligible at lower intensity, greater release was seen at higher intensity (Figure S2c). Therefore, the increased rate of ligand ion release at higher temperatures is a probable mechanism for Zn^2+^ release from phanorod-Zn upon photothermal heating.

It is also possible that laser irradiation might denature Pol-K of the phanorod-Zn, reducing Zn^2+^-loading and consequently causing release of Zn^2+^. We measured the loading capacity of phanorod-Zn, with or without irradiation, by incubating with a solution of ZnCl_2_, separating phanorod-Zn by centrifugation and washing, and quantifying bound Zn^2+^ by ICP-MS. The results showed that laser irradiation (15 min) does not measurably affect loading capacity (Figure S2b). Since the coordination strength could be affected without loss of loading capacity, we also tested whether laser irradiation caused destruction of the thiol groups required to coordinate Zn^2+^ (e.g., by denaturation through oxidation)^45^. We quantified the thiol groups by Ellman’s assay after irradiation, and found that irradiation caused a 6.1% decrease in thiols (from 1.23 +/- 0.035 mM to 1.15 +/- 0.026 mM thiols; *p* = 0.0034). Although this decrease in thiols did not appear to reduce overall loading capacity, it may reduce the coordination strength and thus increase the rate of Zn^2+^ release.

### Photothermal stability and efficiency of phanorods and phanorod-Zn

The photothermal stability and efficiency of AuNR, phanorods and phanorod-Zn solutions ([Au] = 3.3 μM) was measured under gradually increased 808 nm near-infrared (NIR) laser irradiation (0.3 W cm^−2^), close to the absorbance peak of the AuNRs (Figure S1g-h). The solution temperatures of AuNRs, phanorods and phanorod-Zn reached 59.8 ± 0.78 °C, 56.6 ± 0.62 °C and 55.2 ± 0.17 °C after 5 min, respectively (n = 5 in all cases). Under the same treatment, the temperature of PBS buffer increased by < 2°C, confirming that the AuNRs, phanorods and phanorod-Zn efficiently convert the energy of NIR light into heat. We also assessed photothermal reversibility and stability over multiple irradiation cycles (Figure S3). No significant deterioration was observed over five on/off cycles. To determine whether conjugation of phages affected the photothermal conversion efficiency of the AuNRs, the conversion efficiency (η) of light into heat was quantitatively estimated by the time constant (τ_s_) and the maximum steady-state temperature. The photothermal conversion efficiency was slightly perturbed by phage modification, with η = 42.6%, 40.6%, and 38.9% for AuNRs, phanorods, and phanorod-Zn (Figure S3), respectively, which is consistent with an enhancement in scattered light by the phage and peptide^46–47^. Nevertheless, photothermal properties of the AuNRs were largely preserved in the bioconjugates.

### Stability and infectivity of bioconjugates

We studied the stability of phanorod-Zn after storing at 4°C in a sealed flask (under air) for 15 days. The photothermal heating curves, binding curves for *P. aeruginosa*, and Zn^2+^-loading capacity are similar when measured on day 1 and day 15 (Figure S2d-f), indicating that these properties of the bioconjugates were stable during this storage.

To determine whether the biological activity of the bioconjugates was affected by laser irradiation, we measured replication of phages after irradiation. Since the chimeric phage M13- g3p(Pf1) is not expected to propagate in *P. aeruginosa* or to attach to *E. coli* due to the mismatch of host species, we studied bioconjugates using M13KE phage, i.e., M13KE–AuNR and M13KE-AuNR-Pol-K. These bioconjugates were prepared and irradiated with laser by the same method as described above. Afterwards, their ability to propagate on *E. coli* ER2738 was tested by incubation with the host bacteria. Putative phage DNA was extracted and quantified with qPCR^22^. No DNA was detected from propagation of the treated sample (Figure S4), indicating that the infectivity of the phages was inactivated by the laser treatment, as expected due to thermal denaturation of the phages. These results are consistent with previous observations^37^.

### Antibacterial activity of phanorod-Zn with planktonic *P. aeruginosa in vitro*

Prior work ^24^ demonstrated the bacteriocidal effect of phanorods *in vitro*. Conjugation of Pol-K and Zn^2+^- loading of phanorods (i.e., phanorod-Zn) was expected to increase the antibacterial effect due to the known antimicrobial effects of Zn^2+^.^48–49^ We first tested the bacteriocidal activity of phanorod-Zn in a suspension of *P. aeruginosa* cells. Irradiation of phanorods and phanorod-Zn (10^14^ particles/mL) by 808 nm near-infrared (NIR) light (0.3 W cm^−2^) for 15 min resulted in an equilibrium bulk temperature of ∼55 °C (Figure S1). Without irradiation, *P. aeruginosa* suspensions incubated with phanorods or phanorod-Zn showed no antibacterial activity (Figure 2a). However, after 15 min of irradiation, approximately 88.5 ± 3.1% and 97.5 ± 1.5% of *P. aeruginosa* colony-forming units were destroyed in the phanorod and phanorod-Zn groups, respectively, showing significantly greater cell-killing by phanorod-Zn than phanorods (*p* < 0.001; Figure 2b). Transmission electron microscopy (TEM) of *P. aeruginosa* cells showed that phanorod-Zn carried multiple AuNRs to the cells, and cell morphology was grossly altered after photothermal treatment with phanorod-Zn (Figure S5a-c, f-g).

**Figure 2.**
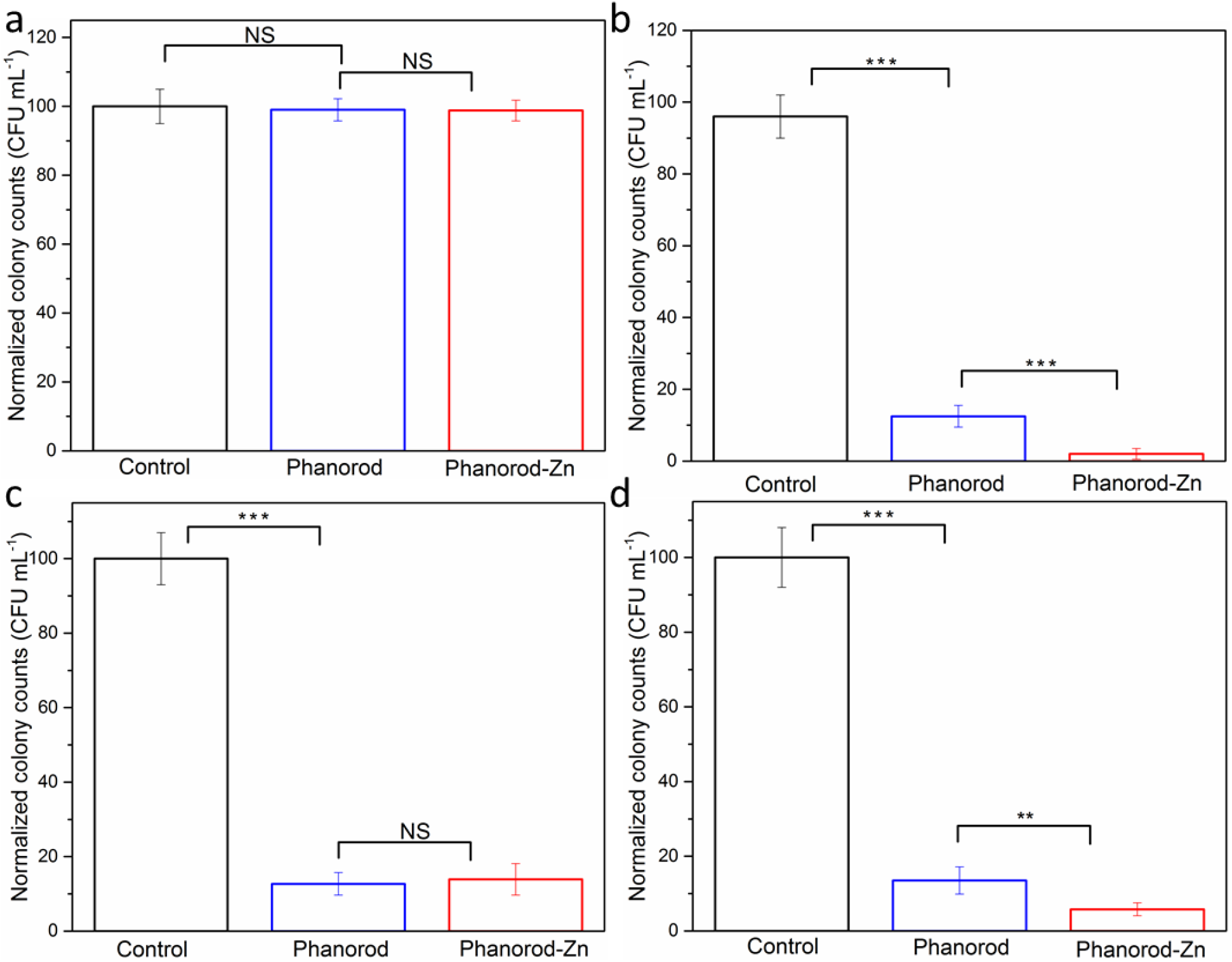
Antibacterial effect of phanorods and phanorod-Zn *in vitro*, measured by colony- forming units. *P. aeruginosa* cells in suspension were exposed to phanorods and phanorod-Zn without irradiation. The presence of these reagents did not affect bacterial survival (a) (control = no phanorods or phanorod-Zn added). With NIR laser irradiation, cell survival was reduced in the presence of phanorods, and phanorod-Zn showed a significantly greater reduction in cell survival (b). If repeated washing was applied to remove released Zn^2+^ (c), the colony counts (with NIR laser irradiation) were similar between phanorods and phanorod-Zn exposure. Colony counts of a treated *P. aeruginosa* biofilm (d) show trends similar to *P. aeruginosa* in suspension (b). Error bars are standard deviation (n = 5): \*\**p* < 0.01, \*\*\**p* < 0.001, NS, not significant (*p* > 0.05, two-sided *t* test).

Live/dead staining confirmed that phanorod-Zn resulted in significantly greater bacterial cell-killing compared to phanorods (phanorods: 87.2 ± 3.9% cell death; phanorod-Zn: 98.8 ± 1.1% cell death; *P* < 0.001) (Figure S6-7). Because the heating profiles of phanorods and phanorod-Zn are similar (Figure S1g and S3), the higher cell-killing efficiency of phanorod-Zn was suspected to be due the released Zn^2+^ ions. To test whether released Zn^2+^ ions might account for the difference, the bacteria-phanorod-Zn complexes were centrifuged and washed in PBS buffer three times after irradiation, to remove Zn^2+^ ions. Washing the bacteria-phanorod-Zn complexes resulted in lower antibacterial efficiency (86.1 ± 4.2% cell death) compared with intact phanorod-Zn (97.5 ± 1.5%; *p* < 0.001), and was comparable to the efficiency of phanorods (Figure 2c). This result supported the interpretation that released Zn^2+^ ions improved antibacterial efficiency by ∼10%, for phanorod-Zn compared to phanorods ^50^.

Because nonspecific effects may occur due to bulk heating of the suspension, we tested the specificity of bacteriocidal activity of phanorods and phanorod-Zn by incubation with *E. coli* and *Vibrio cholerae* (Figure S8). Some reduction (∼20%) in colony-forming units was observed, but survival of these species was substantially greater than that observed for *P. aeruginosa*. The difference in bacteriocidal efficiency is consistent with the engineered specificity of the phage M13-g3p(Pf1) ^24^. A similar degree of non-specific cell-killing was also observed with AuNRs alone under the same conditions (Figure S15b).

### Antibacterial activity of phanorod-Zn with *P. aeruginosa* biofilms *in vitro*

*P. aeruginosa* is known to produce robust biofilms, leading to persistent infections and increased antimicrobial resistance ^51–52^. *P. aeruginosa* biofilms were grown *in vitro* and exposed to phanorods and phanorod-Zn. Consistent with the results from planktonic culture, the antibacterial efficiency for *P. aeruginosa* biofilm eradication by phanorod-Zn was significantly higher (94.2 ± 1.7%) than for phanorods (86.5 ± 3.6%, *P* < 0.01) (Figure 2d). Consistent with this, live/dead staining of the biofilm showed significantly greater dead cells (97.6 ± 1.3%) with phanorod-Zn compared to phanorods (84.7 ± 1.5% dead cells; *p* < 0.0001) (Figure S9 and S10). These studies verified that phanorod-Zn was ∼10-13% more effective than phanorods in the biofilm, quantitatively consistent with the results obtained for planktonic culture. To determine whether the cell-killing was effective deeper into the biofilm, confocal image slices of the live/dead staining were used to reconstruct a three-dimensional image, which showed efficient cell-killing throughout the biofilm under these conditions (Figure S11a, b).

### Antibacterial activity of antibody-AuNRs with *P. aeruginosa in vitro*

For comparison to phanorods and phanorod-Zn, gold nanorods (AuNRs) were also conjugated to a monoclonal antibody (1010/287) that recognizes *P. aeruginosa* serotype 6 (ATCC25102 (Schroeter) Migula) ^24, 53–54^. Photothermal properties and absorbance spectra of antibody-AuNRs were confirmed to be similar to phanorods and phanorod-Zn (Figure S1g-h). Binding of the antibody-AuNRs to *P. aeruginosa* cells was confirmed by transmission electron microscopy (TEM), which showed antibody-AuNRs bound to the surfaces of *P. aeruginosa* cells, with ∼50 antibody-AuNRs per cell (Figure S5d-e). As expected, free AuNRs did not associate with *P. aeruginosa* cells. The antibacterial efficiency of antibody-AuNRs was measured using a particle suspension containing the same concentration of Au as the phanorod and phanorod-Zn suspensions. In suspension, cell-killing was less efficient with antibody-AuNRs compared to phanorods and phanorod-Zn (70- 80% cell death; Figure S12a, S13b), and even lower in a biofilm (55-65% cell death; Figure S11c, S12b, S14b).

### Comparison of antibacterial effect of phanorod-Zn, phanorods with free Zn^2+^, and AuNR- Pol-K *in vitro*

The results above showed that the antibacterial efficiency of phanorod-Zn was ∼10% higher than that of phanorods *in vitro*. In addition, if released Zn^2+^ ions were removed by centrifugation and washing, the antibacterial efficiency of phanorod-Zn dropped to that of phanorods, suggesting that the released Zn^2+^ improved the antimicrobial efficiency. To understand whether these two components, phanorods and Zn^2+^, interacted additively or synergistically with each other to produce the antibacterial effect of phanorod-Zn, we compared phanorod-Zn to an equivalent preparation of phanorods mixed with free Zn^2+^. The concentration of Zn^2+^ released after one irradiation *in vitro* was measured to be 1.06 ppm at 24 h (Figure 1a). Alone, Zn^2+^ of this concentration showed insignificant bacterial killing (Figure S15b). Phanorods were compared to a mixture of phanorods and Zn^2+^ (1.06 ppm). No significant difference was seen in cell-killing rates of the phanorods (4.5± 2.2%) compared to phanorods mixed with Zn^2+^ (6.3 ± 2.8%; *p* > 0.05), given irradiation achieving low temperature (45°C; Figure S15a).

However, at higher laser flux achieving higher temperature, the cell-killing rates of both groups increased, with the phanorods mixed with Zn^2+^ being significantly more effective than phanorods alone (at 55°C: 87.5±1.8% cell-killing by phanorods and 94.5±1.1% cell-killing by phanorods mixed with Zn^2+^; *p* < 0.001) (Figure S15a). Nevertheless, phanorod-Zn (97.5 ± 1.5%) was still significantly more effective compared to phanorods mixed with Zn^2+^ (*p* < 0.001), indicating that phanorods and Zn^2+^ interact synergistically to augment bacteriocidal activity in phanorod-Zn.

To assess the importance of targeting the bioconjugates to the host cells, we conjugated AuNRs to Pol-K and loaded them with Zn^2+^. These bioconjugates would release Zn^2+^ upon photothermal heating, but lack binding activity for the bacterial cells. The Zn^2+^-loaded AuNR- Pol-K shows significantly lower antimicrobial efficiency than phanorod-Zn (*p* < 0.001; Figure S15b; Figure 2b), illustrating the importance of targeting using phage.

### *In vitro* cytocompatibility of phanorods, phanorod-Zn, and Zn^2+^

The *in vitro* cytocompatibility of the phanorods and phanorod-Zn with mammalian cells was studied by the PrestoBlue cell viability assay using human embryonic kidney (HEK) 293T cells. The results (Figure S15c) indicate that the bioconjugates are nontoxic to the HEK293T cells, with cell viabilities above 98% for all concentrations tested (up to 0.4 μM Au). For free Zn^2+^ (Figure S15d), low concentrations (2 ppm and below) were found to be nontoxic to the HEK293T cells, with cell viability of >98%. However, cell viability was substantially decreased at [Zn^2+^] higher than 6 ppm. The maximum cumulative Zn^2+^ concentration released by phanorod-Zn is 1.69 ppm (Figure 1), given laser irradiation for 15 mins every 24 h, for 72 h. Zn^2+^ at this concentration (1.69 ppm) displays insignificant cytotoxicity toward HEK293T cells, indicating that the amount of Zn^2+^ released during phanorod-Zn treatment would not decrease the viability of these cells.

### Treatment of *P. aeruginosa*-infected wounds by phanorods and phanorod-Zn *in vivo*

We evaluated the *in vivo* efficacy of phanorods and phanorod-Zn for treatment of *P. aeruginosa-* infected wounds using a mouse model. Wounds were introduced dorsally by scissors on anesthetized mice and inoculated with *P. aeruginosa* (1 x 10^7^ cfu). For initial experiments, after incubation for 1 h, phanorods or phanorod-Zn were applied topically to the wound, incubated for 30 min, and activated by exposure to NIR irradiation for 15 min (808 nm, 0.3-4 W cm^−2^). Mice were monitored by thermal imaging to measure the temperature of the wound during treatment. Higher laser intensity (0.35-4 W cm^−2^) caused heating beyond 55 °C, resulting in a burn characterized by macroscopic discoloration, inflammation, and neutrophil infiltration (Figure S16). While the higher laser intensities resulted in high bacterial cell-killing rate (91.3±2.1% cell death at 60°C; 94.4±1.1% cell death at 65°C), a moderate laser intensity (0.3 W cm^−2^), causing heating to 55 °C, gave a similar cell-killing rate (90.7±1.2%) (Figure S17) and relatively little tissue damage. This intensity (0.3 W cm^−2^) is also within the scope of permissible skin exposure formulated by the American National Standards Institute (ANSI Z136.1-2000)^55^. Therefore, heating to 55 °C at this laser intensity for 15 min was chosen as the appropriate condition for subsequent experiments.

Wounds were created and infected on day 0. In the control group (wound exposed to PBS buffer; untreated), the wound temperature did not change observably under laser irradiation (Figure 3a). On day 2, macroscopic suppuration and inflammation were observed in the wounds of the control group, and the wound area had approximately doubled from the initial size (Figure 3b,c). Subsequently, the untreated wounds decreased slowly in size, reaching the original wound size on day 8, but were not closed by day 10. When treated with an equivalent amount of free Zn^2+^ (1.06 ppm) on day 1 and 2, the wound area increased in size similar to the control group (Figure 3, S19b), as expected since this concentration of Zn^2+^ is known to be too low to prevent the formation of biofilms by most pathogenic bacteria^56^.

**Figure 3.**
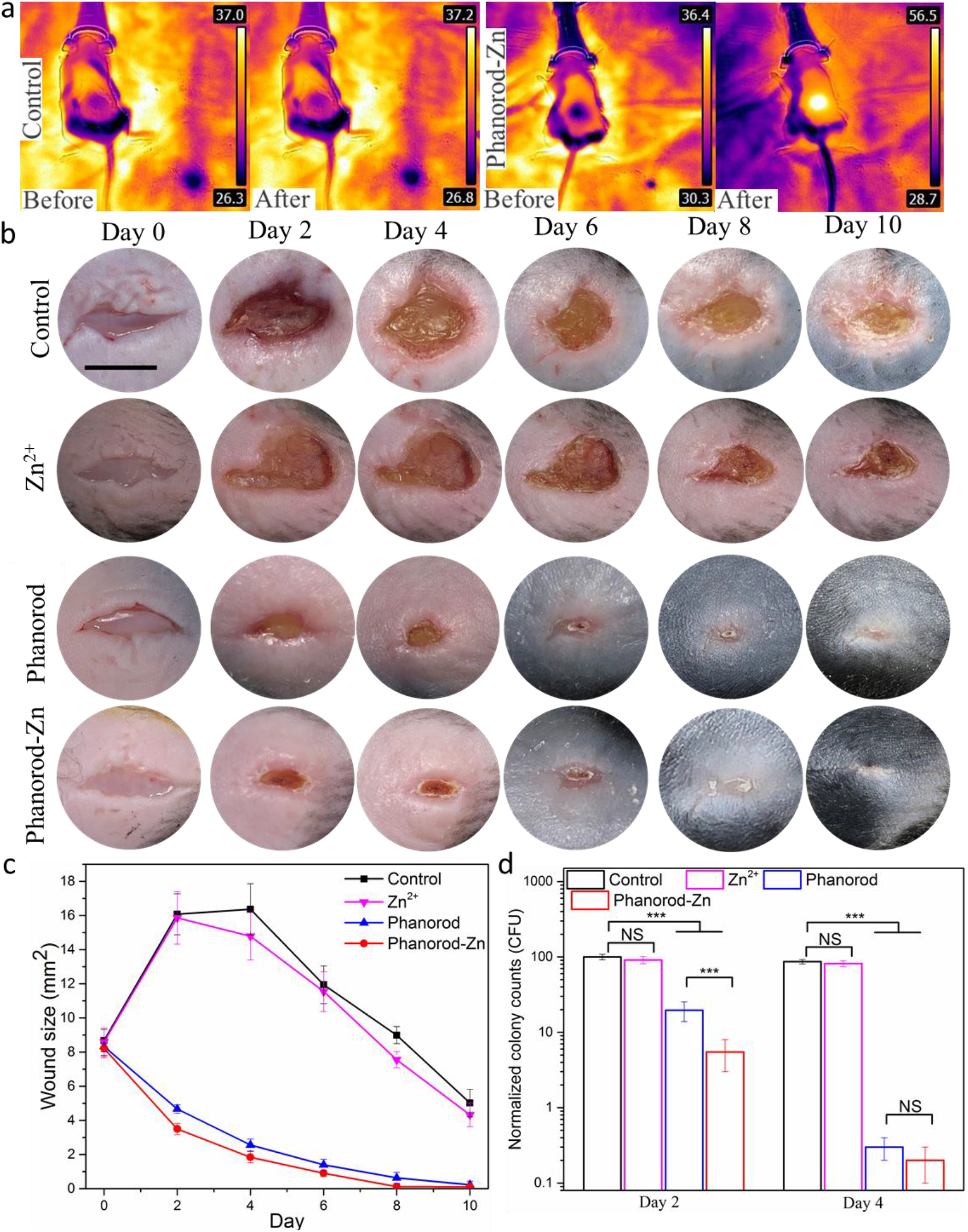
Treatment of *P. aeruginosa*-infected wounds by phanorod-Zn *in vivo*. (a) Thermal images of control and phanorod-Zn treatment before and after the NIR laser irradiation. (b) Representative photographs of the infected wound over time for control, Zn^2+^ ion, phanorod, and phanorod-Zn treatment (scale bar = 5 mm). (c) Wound area over time for control, phanorod, and phanorod-Zn treatment. Also see Figure S19. (d) Bacterial load assessed by colony-forming units from wound tissue on days 2 and 4. Error bars show standard deviation (n = 5 mice): *** indicates *p* < 0.001; NS = not significant (*p* > 0.05); two-sided *t* test.

With phanorod and phanorod-Zn treatment (day 0: reagent applied and incubated for 30 min, after wound inoculation, and irradiated for 15 min; day 1: irradiated again for 15 min), the wound temperature rapidly increased to ∼55 °C in 2-3 mins during laser irradiation (Figure 3a, S18). With phanorod and phanorod-Zn treatment, wound size did not increase, and instead decreased markedly over the next several days; by day 4, the wound area with treatment was <20% of the area of an untreated wound. Macroscopic signs of inflammation were minimal, and the wound reached complete closure by day 8. Assessment of bacterial load by colony-forming units in the wound tissue confirmed that the decrease in wound size corresponded to a decrease in cfu by >10-fold by day 2, and >100-fold by day 4, relative to the control (Figure 3d). With phanorod treatment, a similar trend was observed, although healing and cfu decrease appeared slightly less rapid compared to phanorod-Zn (Figure 3c-d). A close comparison of the wound areas of the two groups on different days revealed that phanorod-Zn showed a significantly better therapeutic effect than the phanorods (Figure S19a).

Treatment with phage M13-g3p(Pf1) or unconjugated AuNRs gave results similar to the negative control by wound size and bacterial load (Figure S20), indicating that only the bioconjugated reagents were active. Interestingly, phanorod or phanorod-Zn treatment resulted in a wound closure rate that was slightly faster than that of non-infected wounds (i.e., had not been inoculated with *P. aeruginosa*), suggesting that hyperthermia and/or Zn^2+^-release may be slightly beneficial even in the absence of overt infection (Figure S21) ^57–58^. The reason for this slight benefit is unclear; because the mice were not kept in a sterile environment, it is possible that wounds that were not inoculated nevertheless harbored a small population of bacteria, and treatment reduced the burden of these cells. In addition, local heating might increase lymphocyte extravasation^59^, blood flow, and oxygen tension^60^, which may be beneficial to the innate immune response.

Phanorod-Zn treatment with a single irradiation was also assessed (irradiation only on day 0). The single irradiation regimen was also effective, with a wound closure rate comparable to a non-infected wound, but was less effective than irradiation on two consecutive days (Figure S21). The difference is consistent with the fact that a small percentage of bacteria survive a single irradiation treatment (Figure 2d), and that additional Zn^2+^ is released upon the second treatment (Figure 1b). Overall, these results indicated that phanorod-Zn, and phanorods to a slightly lesser extent, were highly effective for reducing bacterial load and accelerating wound healing in the mouse model.

### Comparison of phanorod-Zn treatment to standard antibiotic therapy *in vivo*

To compare the efficacy of phanorod-Zn treatment to standard therapy, multiple antibiotic and antiseptic treatments were assessed in the same wound infection model. Treatments included two topical antiseptic agents (acetic acid (2% solution) and chlorhexidine (4% solution)), a topical antibiotic (polysporin) ointment, and two systemic antibiotics (oral ciprofloxacin and levofloxacin). While all treatments were effective in promoting wound closure faster than no treatment, phanorod-Zn treatment gave significantly faster wound closure, accompanied by ∼10-fold greater reduction in bacterial load (Figure 4, S22). While wound closure was completed with phanorod-Zn treatment by day 8, none of the other treatments yielded wound closure by day 10. These results indicated that phanorod-Zn treatment was more effective than standard antimicrobial therapies in the mouse model.

**Figure 4.**
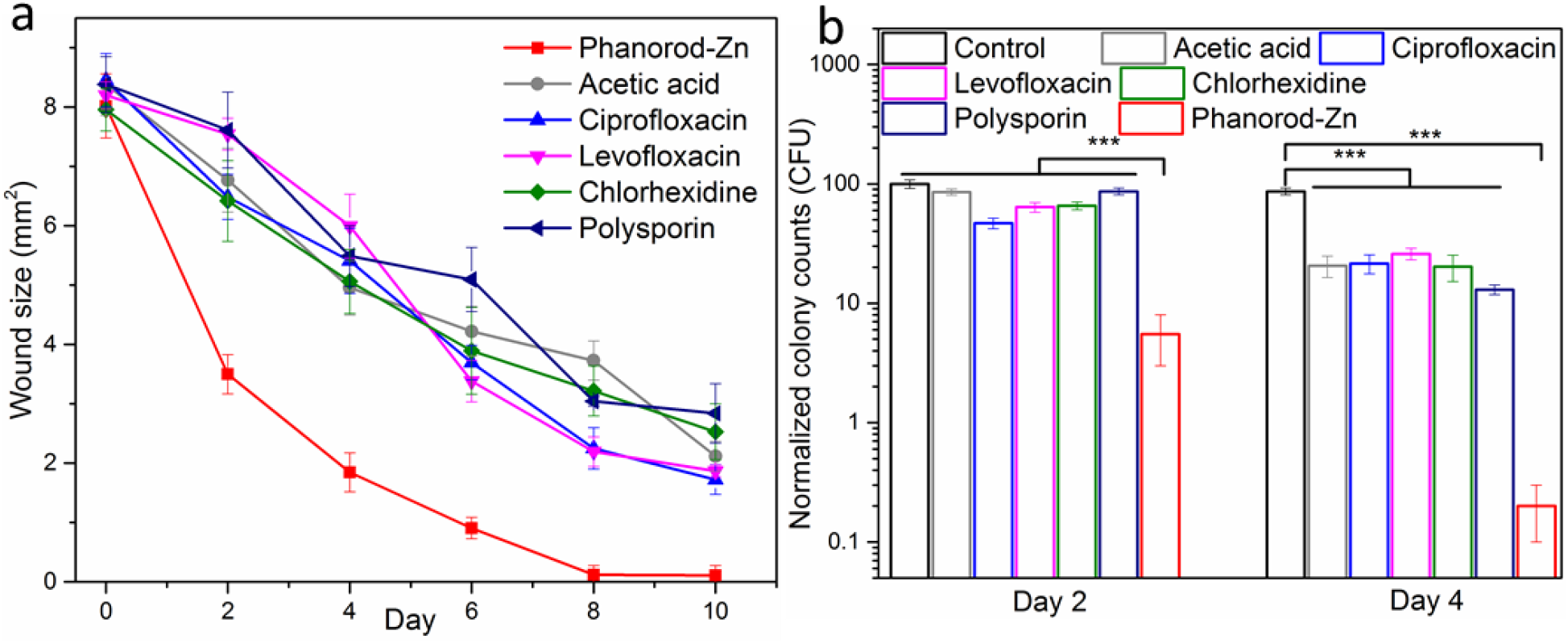
Treatment of infected wounds assessed by wound size (a) over time, comparing antibiotics (ciprofloxacin, levofloxacin, polysporin) and antiseptics (acetic acid, chlorhexidine) with phanorod-Zn. Treatment was initiated on day 0 in all cases. (b) Quantitative assessment of bacterial load (cfu) in wounds during wound treatment. Error bars show standard deviation (n = 5 mice): *** indicates *p* < 0.001 (two-sided *t* test).

### Efficacy of delayed phanorod-Zn treatment *in vivo*

**I**n initial experiments, treatments (including phanorod-Zn and antibiotic therapy) were initiated on day 0, shortly after wound creation and inoculation with *P. aeruginosa*. To determine whether the same treatments were effective late into infection, we tested the effect of treatments initiated once the wounds reached maximum size (day 2). When treatment was delayed, ciprofloxacin treatment was observed to have a minor effect, if any, on the rate of wound closure or bacterial load compared to no treatment (Figure 5, S23), consistent with formation of a biofilm during the two days without treatment. However, with phanorod-Zn treatment beginning on day 2 (with a second round of irradiation performed on day 3), wound size decreased rapidly after day 2, and wound closure was complete on day 10. Unlike delayed ciprofloxacin treatment, bacterial load after delayed phanorod-Zn treatment decreased markedly, consistent with the observed decrease in wound size (Figure 5). These results indicated that, when wound treatment was delayed, phanorod-Zn treatment was still effective, in contrast to ciprofloxacin treatment.

**Figure 5.**
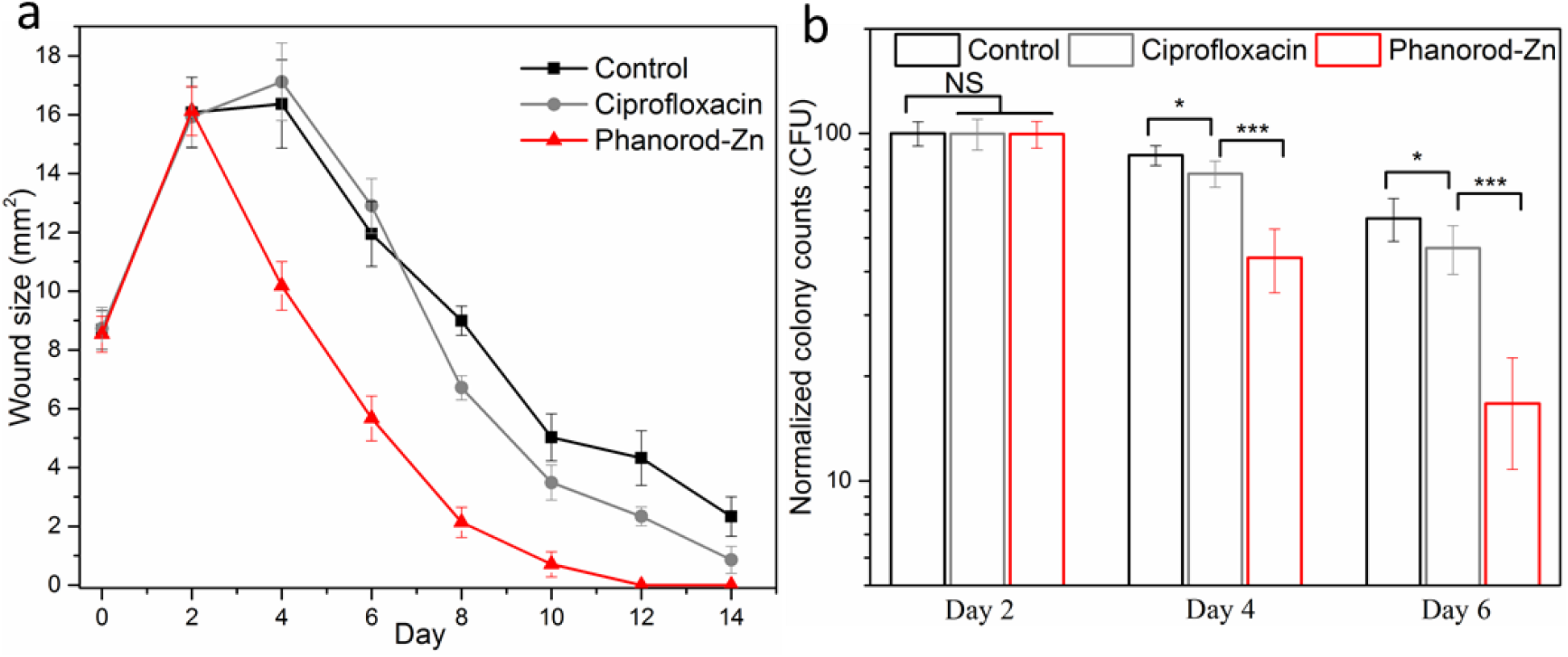
Delayed treatment of infected wounds, with treatment initiated at maximum wound size (day 2). (a) Wound size over time, with either ciprofloxacin or phanorod-Zn treatment begun on day 2. Control = no treatment. (b) Quantitative assessment of bacterial load (cfu) with treatment begun on day 2, assessed on day 2, 4, and 6. Samples on day 2 were taken without treatment. Error bars show standard deviation (n = 5 mice): * indicates *p* < 0.05, *** indicates *p* < 0.001, NS = not significant (*p* > 0.05); two-sided *t* test.

### Efficacy of phanorod-Zn treatment for severe wounds *in vivo*

The wounds described above in this study (initially ∼8 mm^2^, infected by 1 × 10^7^ cfu) were small enough to eventually heal without intervention. While these wounds allowed quantitative comparisons of size and cfu over time, wounds of larger size (∼17 mm^2^) with greater bacterial inoculum (5 × 10^7^ cfu) were also studied to assess treatment of severe cases. For these severe wounds, the control wounds (no treatment: exposure to PBS) showed a mortality rate of 100% (5 out of 5 mice) within three days (Figure 6, S24). However, all mice with severe wounds treated with phanorod-Zn survived (5 out of 5 mice), with wound closure by 10-12 days. Interestingly, even when phanorod-Zn treatment was delayed (initiated on day 2 with second round of irradiation on day 3), all mice survived the severe wounds (5 out of 5 mice), and wound closure was complete by 12-14 days. These results indicate that phanorod-Zn treatment, even if delayed, was effective for rescuing mice from severe wounds that would otherwise result in 100% mortality.

**Figure 6.**
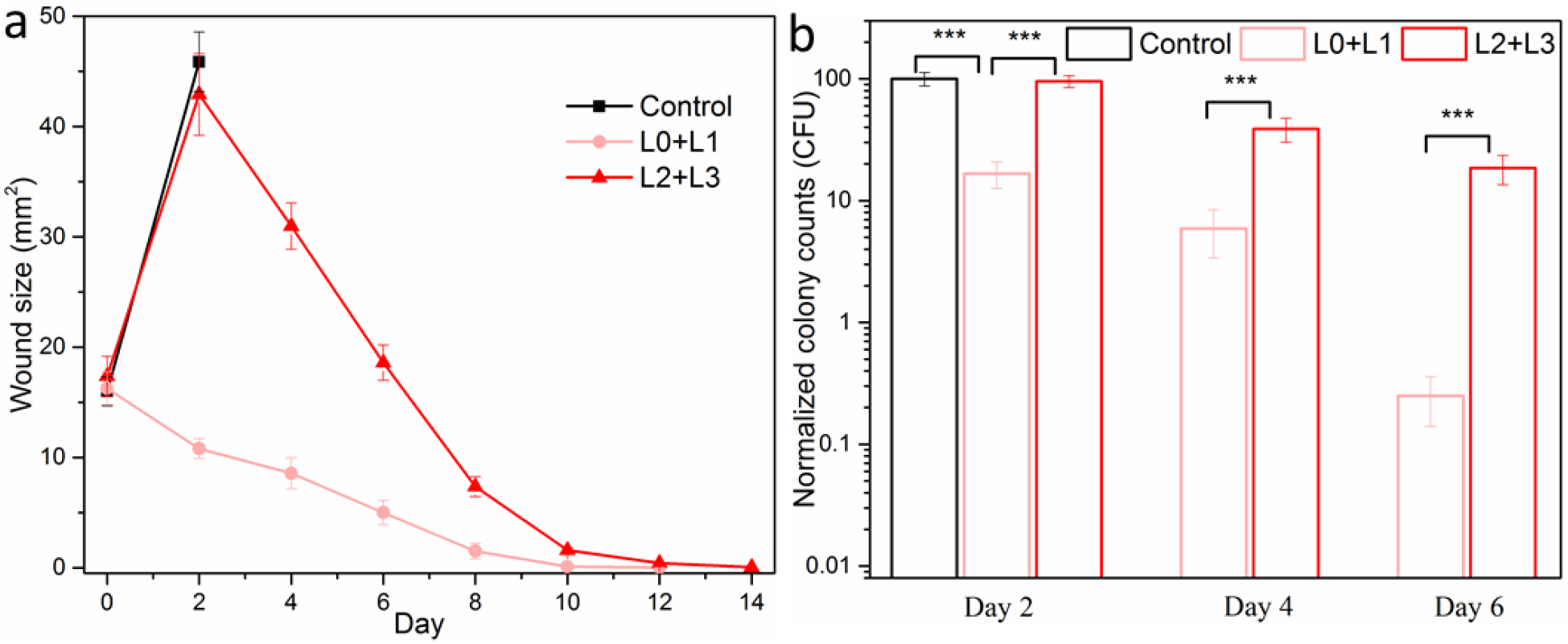
Treatment of severe wound infections by phanorod-Zn, initiated early (L0+L1) or delayed initiation on day 2 (L2+L3). (a) Wound size over time; no data is available for the control group (no treatment) past day 2 due to 100% mortality. (b) Quantitative assessment of bacterial load (cfu) during treatment of severe wound infections, measured on days 2, 4 and 6. For delayed treatment (L2+L3), samples on day 2 were taken without treatment. Error bars show standard deviation (n = 5 mice): *** indicates *p* < 0.001 (two-sided *t* test).

### Treatment of polymyxin-resistant *P. aeruginosa* with phanorod-Zn *in vivo*

The main application of a novel antimicrobial class is anticipated to be treatment of antibiotic-resistant infections. Current last-line therapy for *P. aeruginosa* is polymyxin antibiotics. We tested phanorod-Zn treatment of wounds infected by a polymyxin-resistant strain of *P. aeruginosa*, PAK*pmrB6* ^61–62^. Consistent with this phenotype, polymyxin treatment was ineffective against wounds infected by strain PAK*pmrB6*, while phanorod-Zn treatment was effective, as shown by decreased wound size and decreased bacterial load (Figure 7, S25). Phanorod-Zn treatment delayed to the time of maximum wound size (initiated on day 2) was also effective with this strain (Figure 7). These results confirm that phanorod-Zn treatment is effective against wound infection by an organism resistant to last-line therapy.

**Figure 7.**
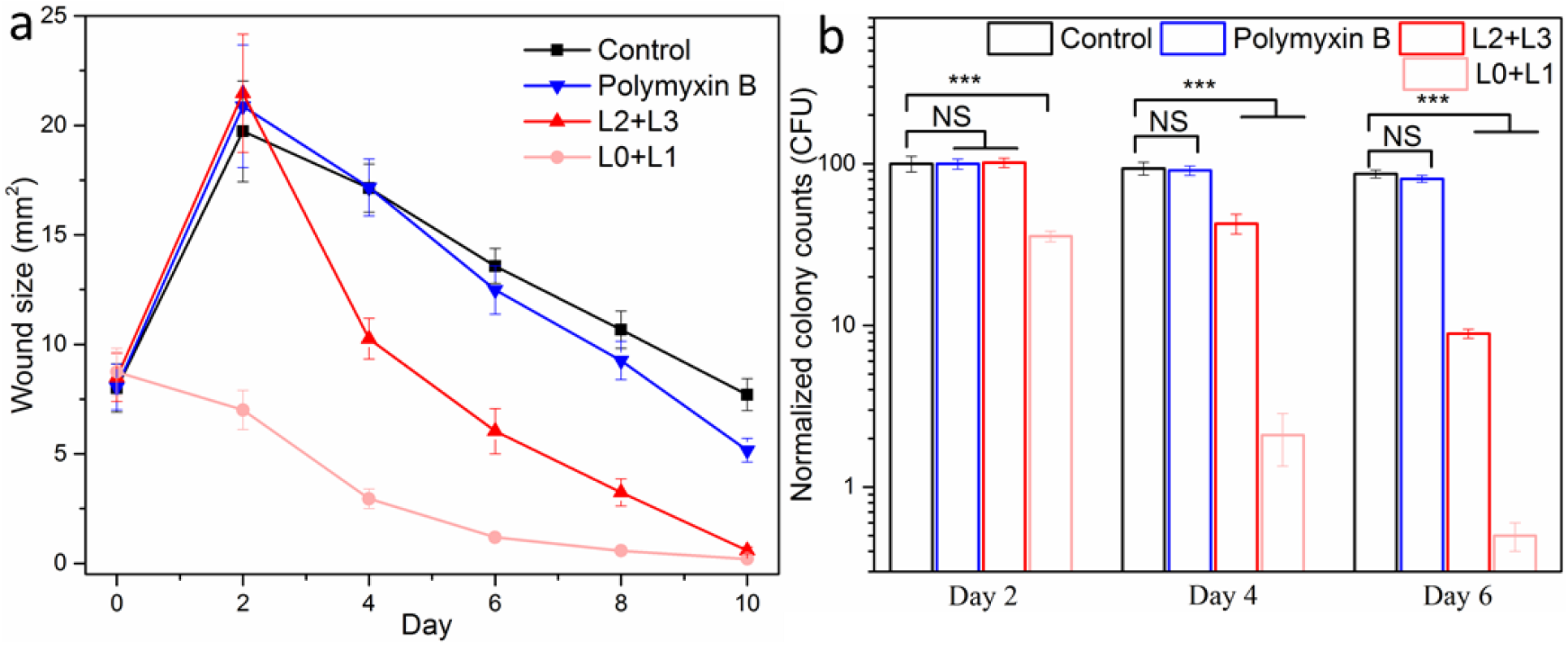
Treatment of wounds infected by polymyxin-resistant *P. aeruginosa*. (a) Wounds assessed by size over time, with polymyxin treatment initiated on day 0, or phanorod-Zn treatment initiated on day 0 (L0+L1) or day 2 (L2+L3). (b) Quantitative assessment of bacterial load (cfu) for polymyxin-resistant *P. aeruginosa* wound infection, measured on days 2, 4 and 6. For delayed treatment (L2+L3), samples on day 2 were taken without treatment. Error bars show standard deviation (n = 5 mice): *** indicates *p* < 0.001; NS = not significant (*p* > 0.05); two-sided *t* test.

### Comparison of phanorod-Zn with antibody-AuNRs *in vivo*

Treatment by gold nanorods conjugated to antibodies (rather than phages) were studied, in order to determine whether phages possessed any advantage over antibodies for targeting bacterial cells. A monoclonal antibody (1010/287) recognizing the *P. aeruginosa* strain used here (PAK, serotype 6) was conjugated to gold nanorods and tested for treatment of *P. aeruginosa*-infected wounds (Figure S20). The antibody-AuNRs were less effective than phanorod-Zn, but gave comparable results to treatment by antibiotics, as shown by the reduction in wound size and bacterial load (Figure S20). Thus, while treatment by antibody-AuNRs was inferior to phanorod-Zn, their moderate efficacy validated the general approach of using an affinity reagent to target gold nanorods toward the bacterial cells for photothermal treatment. Although antibody-AuNRs were less effective than phanorod-Zn, antibody-AuNRs were more effective than Zn^2+^-loaded AuNR-Pol-K, confirming the primary importance of targeting therapeutic agents to the pathogens in antibacterial treatment.

### Histological analysis and collagen deposition during phanorod-Zn treatment *in vivo*

When infected wounds were not treated (exposed to PBS buffer), a high degree of neutrophil infiltration into the wound tissue was observed by hematoxylin and eosin (H&E) staining on day 2 (Figure 8), consistent with acute inflammation and infection. In the control group, neutrophil infiltration is still observed on day 6, indicating prolonged inflammation, consistent with macroscopic signs of inflammation (redness and swelling) seen in the wound photographs for the untreated control group. In comparison, neutrophil infiltration in the phanorod and phanorod-Zn treatment groups was less dense on day 2, and decreased considerably in the following days (Figure 8, S26), consistent with a resolving infection.

**Figure 8.**
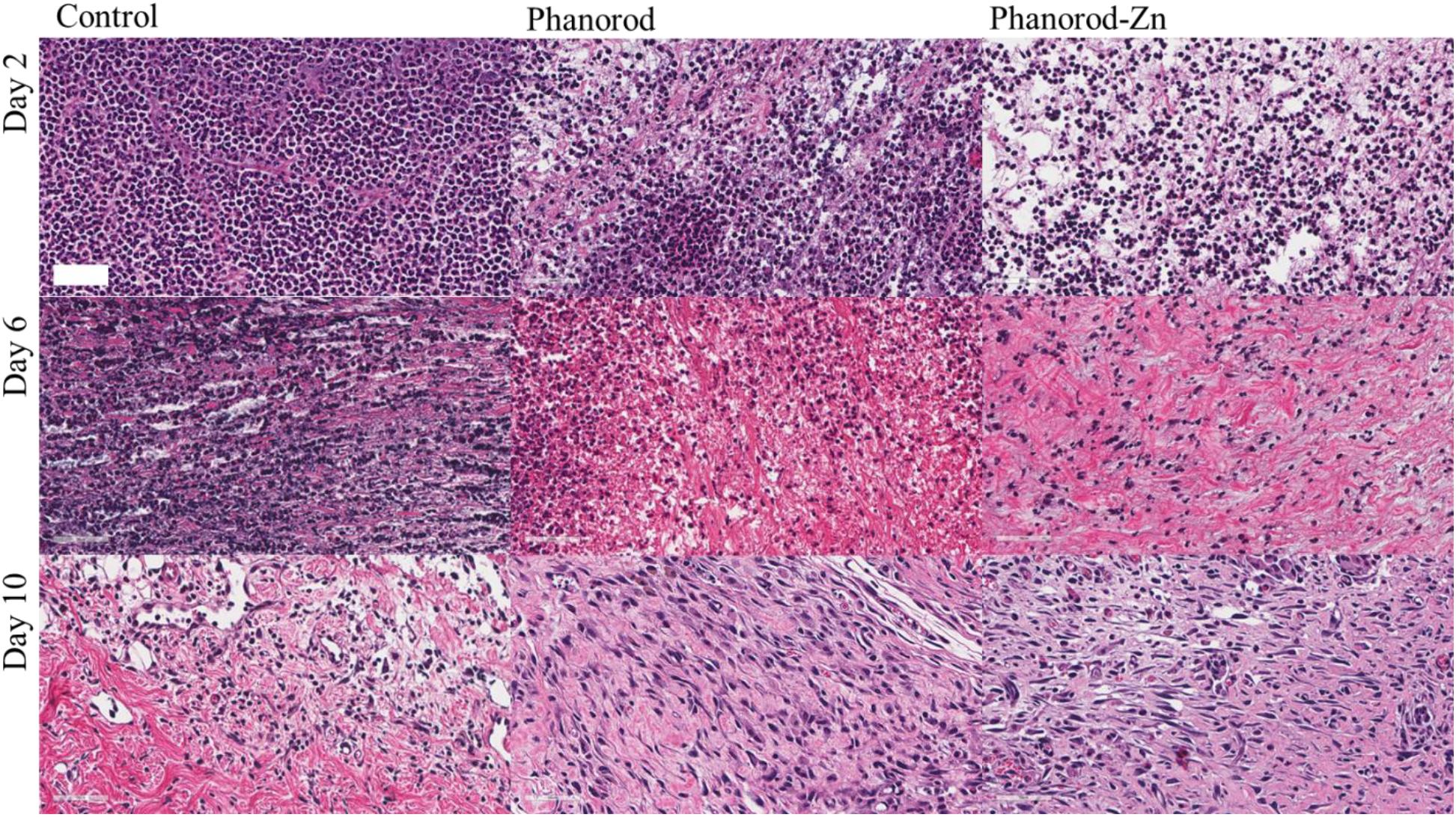
Hematoxylin and eosin staining of wound tissue sections on days 2, 6, and 10. First column: control (no treatment); second column: phanorod treatment; third column: phanorod-Zn treatment. Treatments were initiated on day 0. Scale bar = 50 μm. Neutrophil infiltration is pronounced in the control.

To assess wound repair at a microscopic level, we used Masson’s trichrome staining to monitor collagen deposition in the wounds. Phanorod-Zn treatment resulted in substantially greater collagen deposition than the untreated control throughout the wound healing time frame (Figure S27). Interestingly, phanorod-Zn treatment also gave greater collagen deposition compared to phanorod treatment, indicating that Zn^2+^ release from phanorod-Zn was beneficial for wound healing. Quantitation of collagen area from Masson’s trichrome staining indicated ∼89.4±5.6% collagen coverage of wound tissue sections by day 8 for phanorod-Zn treatment, consistent with wound closure by this day, which was significantly greater than collagen coverage from phanorod treatment (∼54.8±6.5%, *p* < 0.001), which in turn was significantly greater than collagen coverage with no treatment (∼30.9±2.2%, *p* < 0.001) by day 8 (Figure S28). Therefore, phanorod-Zn treatment was associated with reduced inflammation and relatively rapid collagen deposition, consistent with accelerated wound healing.

### Toxicity profile of phanorods and phanorod-Zn *in vivo*

To evaluate possible systemic toxicity of phanorod and phanorod-Zn treatment, major organs (liver, spleen, kidney, heart and lung) were harvested on day 10 and analyzed by H&E staining. No differences were apparent among the untreated, phanorod treatment, and phanorod-Zn treatment groups (Figure S29), showing no histological indication of toxicity. Hematological analysis (hematocrit (HCT), hemoglobin (HGB), mean corpuscular HGB concentration (MCHC), mean corpuscular hemoglobin (MCH), mean corpuscular volume (MCV), platelet count, red blood cell count (RBC), red cell distribution width, and white blood cell count (WBC)) also showed no significant differences among levels in the untreated, phanorod, and phanorod-Zn groups at day 10 (Figure S30a).

To study hepatic and renal toxicity in particular, the biomarkers alkaline phosphatase (ALP), alanine transaminase (ALT), aspartate transaminase (AST), blood urea nitrogen (BUN), creatinine and uric acid were also assessed on day 10. No significant differences were seen for the phanorod and phanorod-Zn treatment groups compared to the untreated group (Figure S30- 32). In contrast, fluoroquinolone and polymyxin treatments resulted in significant elevations, consistent with the known toxicity of these antibiotics.

To test whether phanorod and phanorod-Zn treatment resulted in elevated levels of systemic Au^2+^ or Zn^2+^, the blood concentrations of these ions were measured. Au^2+^ concentrations were undetectable in all groups (limit of detection = 0.002 ng/mL). The serum concentration of Zn^2+^ ions was slightly elevated in the phanorod-Zn group compared to the phanorod and untreated groups, but the difference was not statistically significant (Figure S30b). Overall, phanorod and phanorod-Zn treatments did not show signs of systemic toxicity, as assessed after wound closure.

### Tissue biodistribution of phanorods and phanorod-Zn

To study the biodistribution of phanorods and phanorod-Zn in mice, the concentration of the bioconjugate in the major organs and wound skin was investigated on day 8 following wound infection and treatment, measured by ICP-MS for Au (Supporting Methods). The concentration of Au was >10-fold higher in the wound skin compared to all other organs measured (liver, kidney, spleen, lung, or heart) (Figure S33). Among these organs, the liver and spleen showed higher Au amount compared to other organs. This is consistent with the topical application and washing of the bioconjugates during treatment, and is in accordance with previous studies in biodistributions of nanoparticles for topical wound healing applications^63–64^.

## Discussion

Phage therapy is usually envisioned with lytic phages that self-amplify exponentially to eradicate a bacterial population. However, such a strategy poses multiple concerns about safety and efficacy, particularly if using poorly characterized phages. Yet phages have evolved multiple mechanisms for discovering and attaching to bacterial hosts in natural environments, including receptor-binding proteins whose affinities rival or surpass those of antibodies and subdiffusive ‘search’ mechanisms ^65–69^. By using chimeric M13 phages to carry an activatable, toxic cargo to bacterial pathogens, the phanorod strategy takes advantage of phage attachment while controlling the antimicrobial effect. Unlike phage therapy, the phage component of phanorods does not need to infect or lyse the host cells, limiting the requisite genetic engineering to the receptor-binding interaction, in contrast to strategies that rely on phage infection and gene expression ^70^. Such engineering to bind different host bacterial strains can be achieved by creating chimeric phages, in which the receptor-binding protein of M13 has been replaced by the homolog from a filamentous phage that targets the pathogen; the chimeric phages bind and deliver cargo to the targeted pathogen^20^. At the same time, complete phages may be advantageous over phage proteins (e.g., lysins ^71^) for robustness to exposure to serum and the physiological milieu ^22, 72^. M13 and Pf1 are non-lytic filamentous phages (inoviruses), which bind to specific bacterial pili. In particular, the receptor-binding domain of Pf1 (used in the M13-g3p(Pf1) phanorods) binds to type IV pili, a virulence factor that enables adhesion, motility, and biofilm formation ^73^. In one analysis, 75% of *P. aeruginosa* strains isolated from wound infections expressed type IV pili ^74^. Therefore, filamentous phages may be an appropriate source of receptor-binding domains against pathogens.

The advantage of phage-based targeting of AuNRs was highlighted *in vivo*. An alternative to phage targeting is antibody targeting, which is under study for antibacterial applications ^75–77^. Using an anti-*P. aeruginosa* monoclonal antibody with irradiation intensity tuned to achieve the same wound temperatures, reduction of the wound size to <50% of the initial size required >6 days with antibody-AuNR treatment, but only 2 days with phanorod treatment (Figure S20). This difference could be due to the greater cargo capacity of phanorods, which carry approximately 15 AuNRs per particle ^24^, compared to antibodies, whose size (∼ 12 nm) is unlikely to accommodate more than one AuNR (∼14 × 53 nm) per molecule. Even if a similar number of AuNRs attaches per cell when antibody targeting is used (e.g., with the antibody targeting a highly abundant bacterial receptor compared to pili), activation of multiple AuNRs at the same microscopic location, targeted by phanorods, would create a steeper thermal gradient at that site, compared to activation of the same number of AuNRs spaced apart from one another (Figure S5). TEM images also suggested that phanorods may aggregate together on the bacterial surface, possibly due to multiple interactions, which would further amplify the steepness of the gradient. In addition, phages are more stable than antibodies under harsh chemical conditions and high temperatures ^72, 78^. This latter property may be relevant to the studies here, as early denaturation of the molecular interactions during photothermal heating could lead to dissociation of the bioconjugate from the bacterial cell. Also, while attachment of antibody-AuNRs was verified by TEM in this case, in principle, nonspecific conjugation of AuNRs might be more likely to interfere sterically with the antibody-antigen interaction due to their relative sizes, compared to the phage-receptor interaction, which occurs at one end of the virion (∼9 nm diameter × 1 μm length). Understanding the mechanisms for the advantage of phage- over antibody- targeting would require further study.

While the quantitatively increased efficacy of phanorods over antibody-nanorods is important for individual treatment, it could also be significant at a population level, as under- treatment of microbial infections can lead to development of resistant phenotypes and strains as well as poorer clinical outcomes ^79–82^. Indeed, resistant strains of *P. aeruginosa* emerge in ∼10% of cases during antibiotic treatment ^83^, suggesting that high efficacy is a critical feature for new therapeutic agents. Furthermore, the ability to engineer or evolve phages to overcome bacterial resistance is an important future concern. For the photothermal mechanism described here, resistance mechanisms are presumably limited to disruption of the receptor-binding interaction. A perceived advantage of phage therapy is the potential to conduct directed evolution to overcome bacterial resistance ^84^; given the evolvability of phages against new receptors observed in other studies ^85–88^, such future developments may be feasible.

A well-known problem for antibiotics is penetration through biofilms, as illustrated by the relative lack of efficacy of ciprofloxacin treatment in the mice after the wound infection was left untreated for 2 days. Although we did not measure distribution of the phanorods into the biofilm, the cell-killing effect did not appear to be impeded by a biofilm *in vitro*, to the depth observed (∼9 μm). It should be noted that regardless of whether phanorods themselves penetrate the biofilm, the cytotoxic effect is mediated by the thermal gradient. Since biofilms may be ∼80-95% water ^89–90^, thermal conductivity is likely to be similar to that of water. The excellent efficacy of phanorods and phanorod-Zn *in vivo*, particularly when treatment was delayed for 2 days, also confirmed that the biofilm developed in the wounds of these experiments did not prevent treatment.

An important practical question for phanorod and phanorod-Zn treatment was whether the bacterial population could be effectively eradicated without excessive harm to the animal, either locally or systemically. Greater laser exposure would result in greater bacterial cell-killing, but would also increase the potential for negative side effects from thermal burn at the wound area. Optimization experiments indicated that burns could be minimized by adjusting laser intensity to achieve a certain steady-state temperature (∼55 °C), while maintaining excellent bacteriocidal activity. Another source of potential harm is renal or hepatic toxicity, which limits the use of some antibiotics. For example, current last-line antibiotic treatment for *P. aeruginosa* and other gram-negative infections is polymyxins, a drug class with substantial renal toxicity; in one study, 2/3 of patients treated by colistin developed nephrotoxicity marked by elevated creatinine levels ^91^. In our experiments, indicators of renal and hepatic function indicated compromise after treatment of infected mice by systemic fluoroquinolones and polymyxin antibiotics, as expected. However, no elevations were seen after treatment with phanorods or phanorod-Zn. The lack of systemic toxicity is consistent with the localized application, in which the phanorods or phanorod-Zn were applied and incubated briefly to allow binding to target bacteria before irradiation. The biodistribution assessed here (day 8) showed that relatively little Au was present in organs other than the wound skin itself, consistent with other studies on the biodistribution of nanoparticles in topical wound healing applications^63–64^. However, more work would be needed to assess biodistribution and biosafety over longer time periods. Another concern was whether phanorods or phanorod-Zn might induce a dysfunctional immune response. However, no significant elevation was seen in the white cell count. Other blood cell counts were also statistically indistinguishable from the untreated control, illustrating a lack of measured systemic toxicity.

Phanorod-Zn was designed to be loaded with Zn^2+^ that could be released upon photothermal heating. In these experiments, a temperature change was shown to cause Zn^2+^ release. Similarly, systems such as zinc fingers ^43^ and Zn^2+^-substituted rubredoxins ^42^ are reported to release metal ions like Zn^2+^ upon thermal denaturation. Irradiation also resulted in a decrease in thiol groups. The amount of thiol loss apparently did not reduce the Zn^2+^-loading capacity in this system, possibly because some of the lost thiols were not from Pol-K, and the four thiols of Pol-K may not all be necessary for coordination^92^. However, binding affinity is reduced as the number of coordinating cysteines decreases^92^. Here, both the effects of increased temperature and loss of thiols may contribute to the observed photothermal release of Zn^2+^.

Higher levels of thiol destruction may also affect loading capacity and lead to greater release of Zn^2+^. While Zn^2+^ is an important trace metal for bacterial growth when present at low concentrations, higher concentrations are inhibitory, due to competition for metal-binding sites in proteins as well as direct association with bacterial cell membranes, leading to rupture ^93^. Indeed, phanorod-Zn was more effective at cell-killing compared to phanorods in our experiments, particularly *in vitro*, which appeared to be due to a released solute (presumed Zn^2+^). At the same time, Zn^2+^ is relatively well-tolerated in humans ^94^, and zinc oxide, in particular, has been shown to cause improvement in wound healing in multiple animal and clinical studies ^38^. Zinc is reported to be involved in regulating several phases of the wound healing process, including membrane repair, oxidative stress, coagulation, inflammation and immune defense, tissue re-epithelialization, angiogenesis, and fibrosis formation ^95^. Although still unclear, release of Zn^2+^ is believed to be one of the mechanisms by which zinc oxide affects wound healing. Zn^2+^ is favorable for deposition of collagen, an essential component of tissue repair ^96–98^. Indeed, our studies suggested greater collagen deposition after phanorod-Zn treatment compared to phanorod treatment, although it should be noted that the long-term effect of this deposition is unknown. *In vivo*, phanorod-Zn treatment appeared to be slightly more effective than phanorods in terms of wound reduction and cfu reduction, but these differences were not always statistically significant; more work is needed to clarify the overall effect of zinc release in this system.

The TEM images and prior microscopy with molecular rotors^37^ indicate that photothermal activation of phanorods and phanorod-Zn cause severe damage to bacterial cells and membranes, such that cellular structures were no longer intact. Zn^2+^ is a micronutrient, but at high concentrations Zn^2+^ is toxic to bacteria^99–101^, although the molecular basis of toxicity remains poorly understood. Very high concentrations of Zn^2+^ (336 ppm) significantly alter membrane permeability of bacterial cells and result in cell death^102^. A plausible mechanism for the observed synergistic interaction of phanorods with Zn^2+^, resulting in increased toxicity of phanorod-Zn compared to a mixture of phanorods and free Zn^2+^, may be that localized photothermal heating disrupts the cell membrane, allowing the released Zn^2+^ ions to penetrate the cells. Nevertheless, photothermal ablation appears to be the dominant mechanism of toxicity, given the high efficacy of phanorods alone. Aside from toxicity, another property of phanorod- Zn appears to be improved wound healing compared to phanorods, as suggested by greater collagen deposition and significantly faster decrease in wound size. This difference is also consistent with the release Zn^2+^ ions, which are known to be associated with improved wound healing^38, 103^.

In this work, we validated phanorods and phanorod-Zn as a potential antimicrobial treatment in a mouse model of *P. aeruginosa* wound infection. Although application of phanorods may be less clinically convenient than orally administered antibiotics, this work shows how the practical challenges of phanorod treatment *in vivo*, such monitoring wound temperature, could be surmounted. An important consideration for future application is whether phanorods can be engineered to target particular pathogens. In general, phages, which determine specificity in this system, may be highly specific or have relatively broad host range ^104^. Since phanorods rely only on attachment, expression of the appropriate receptor is expected to be the major, and possibly the sole, requirement for bacterial susceptibility. Some receptors, such as virulence factors, may be expressed by a substantial fraction of pathogens; further work is needed to define the frequency of targetable receptors in pathogenic populations. Given a susceptible organism, phanorod or phanorod-Zn treatment may be advantageous in terms of efficacy and safety compared to standard-of-care antibiotic therapy, particularly for established biofilms, and the high specificity and localized treatment may be advantageous for avoiding undesirable effects on the microbiome ^105–106^. Regardless, the main benefit of a new antimicrobial strategy would be as a last-line therapy for multidrug-resistant organisms.

## Materials and Methods

### Materials

Reagents were obtained from the following sources: gold(III) chloride trihydrate (HAuCl_3_, 99.9%, Sigma), sodium borohydride (NaBH_4_, 98%, Fisher Scientific), trisodium citrate dihydrate (99.9%; Sigma), *P. aeruginosa* (Schroeter) Migula (ATCC 25102; PAK strain), *P. aeruginosa* PAK-pmrB (gift of Prof. Jian Li, Monash University), *V. cholerae* 0395 (gift of Prof. Michael J. Mahan, UCSB), M13KE phage (New England Biolabs), M13-NotI-Kan construct^41^, sodium chloride (NaCl, 99%, Fisher BioReagents), tryptone (99%, Fisher BioReagents), yeast extract (99%, Fisher BioReagents), *E. coli* ER2738 (New England Biolabs), *N*-(3- (dimethylamino)propyl)-*N′*-ethylcarbodiimide hydrochloride (EDC, 99%, Sigma), N-hydroxysuccinimide (NHS, 98%, Sigma), *N*-succinimidyl-*S*-acetylthiopropionate (SATP) (Thermo Fisher Scientific), 5-bromosalicylic acid (5-BAA) (>98.0%; TCI), isopropyl β-D-1- thiogalactopyranoside (IPTG) (99%; Fisher Scientific), thiol-PEG-acid (HOOC-PEG-SH; PEG average M_n_ 5,000; Sigma), poly(ethylene glycol) (PEG-8000, Sigma), dialysis kit (MWCO 3500 Da, Spectrum Laboratories), tetracycline (Sigma), kanamycin sulfate (Sigma), Top 10F′ cyan cells (Thermo Fisher), Mix and Go competent cells (Zymo Research), QIAprep Spin Miniprep Kit (Qiagen), QIAquick Gel Extraction Kit (Qiagen), and KpnI-HF/NotI-HF restriction enzyme and T4 DNA ligase (New England Biolabs), *Pseudomonas aeruginosa* antibody (1010/287) (Novus Biologicals, LLC).

### Pol-K peptide design and synthesis

The CXCX_3_CX_2_C motif was previously identified in ca. 6% of the Zn^2+^-binding sequences of the protein data bank (PDB) ^107^. Unlike other Zn^2+^-binding motifs that can possess ligands to the metal center hundreds of amino acids apart, this specific motif is only 10 amino acids long and is among the shortest known binding sites that completely coordinate a metal ion. The Pol-K peptide GCFCEDACDKCG was based on the Zn^2+^-binding sequence CFCEDHCDKC from RNA Polymerase II (PDB ID: 1i3q, chain C)^108^. Glycines were added to the extremities of the peptide to decrease the likelihood of interference from N- and C-termini on coordination, while also facilitating functionalization by providing spacing to the cysteine ligands. The histidine residue was substituted with an alanine to remove competition with another potential ligand.

Reagents for peptide synthesis were obtained from Sigma-Aldrich and used without further purification. The synthesis of the Pol-K peptide was performed according to standard Fmoc-based solid-phase peptide synthesis procedures ^109^. N,N-dimethyl formamide (DMF) was used as the solvent, and Wang resin was used as the starting polymeric support. Fmoc-protected amino acids (AA) were used. Peptide elongation was by Fmoc-deprotection of the residue anchored to the resin and Fmoc-AA-OH coupling. Fmoc-deprotection was by washing with 20% (v/v) piperidine in DMF. For each coupling, an excess (Fmoc-AA-OH: anchored AA, 3:1) of the Fmoc-α-amino acid derivative was added to the resin. Apart from Fmoc-Cys(Trt)-OH, Fmoc-α- amino acid derivatives were activated with a mixture of hydroxyl-benzotriazole (HOBt), N,N,N′,N′-tetramethyl-O-(benzotriazol-1-yl)uronium tetrafluoborate (TBTU), and N,N- diisopropylethyl amine (DIPEA). Fmoc-Cys(Trt)OH was activated with a N,N’- diisopropylcarbodiimide (DIC)/HOBt mixture. The peptide was cleaved from the resin and deprotected by treatment with a solution of trifluoroacetic acid (TFA):H_2_O:triisopropyl silane (TIS):1,2-ethanedithiol (EDT) (volume ratio 37:1:1:1) for 2 h, and the product was precipitated with a cold solution of diethyl ether followed by washing cycles with diethyl ether. Finally, the peptide was dried under vacuum. The successful synthesis was confirmed by mass spectrometry using an Agilent 6530 LC-QTOF instrument (Agilent Technologies, Inc.) with electrospray ionization (Figure S1e).

Deionized (Milli-Q, Millipore) purified water was deoxygenated by distillation under nitrogen. Solutions of Pol-K peptide were handled under controlled nitrogen atmosphere with a Schlenk line and Schlenk glassware and transferred to an anaerobic glovebox (Iteco Engineering) for metallation and spectral acquisition. UV-Vis absorption spectra of freshly prepared solutions of peptido-metal complexes were recorded with a Thermo Scientific Evolution 60S UV-Visible Spectrophotometer inside of an anaerobic glovebox (Iteco Engineering). Hellma high-performance quartz glass (QS) cuvettes were used for the acquisition.

### Peptide Zn^2+^-binding assay

To overcome the spectroscopic silence of Zn^2+^, competitive binding experiments were performed using a preformed peptido-Co^2+^ complex. To form the saturated peptido-Co^2+^ complex, aliquots from stocks of 333 mM and 33 mM metal salt solution (CoCl_2_·6H_2_O) were added into a cuvette containing 1 mL of 1 mM Pol-K peptide and 20 mM Glycylglycine at pH 8.7. UV-Vis spectra were collected upon each addition. Absorbance values at 750 nm, diagnostic for tetrahedral 4S→Co^2+^ complexes ^44^, were monitored until no further changes were observed. The *K_d_* of the peptido-Co^2+^ complex was calculated by fitting the absorbance data to the equation:

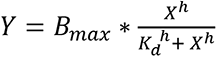

where *Y* is the absorbance, *B_max_* is the absorbance at saturation, *X* is the concentration of Co^2+^, and *h* is the Hill coefficient. GraphPad Prism v. 6.00 (GraphPad Software, La Jolla California USA) for Windows was used to calculate the *K_d_* values.

To assay the competitive binding of Zn^2+^, a decrease in the absorbance at 750 nm indicative of the peptido-Co^2+^ complex was monitored after additions of ZnSO_4_. The concentration of Co^2+^ added to the peptide before the competition assay was equal to the *K_d_* of the Pol-K-Co^2+^ complex, as determined in saturation binding experiments. UV-Vis spectra were collected upon each addition. Titration continued until no change in absorbance at 750 nm was observed. The *K_d_* value of the peptide-Zn^2+^ complex was calculated by fitting to a revised Cheng-Prusoff equation for competitive inhibition, as previously described ^110^ (Figure S1a-c).

### Phage, bacteria, and gold nanorods (AuNRs)

See Supplementary Methods.

### Synthesis of phanorods and phanorod-Zn

To prepare phanorods, phages were chemically thiolated using SATP and conjugated to the AuNRs, and trace CTAB was exchanged for HS-PEG-COOH ^24^. Phanorod-Zn was prepared by coupling the -COOH groups of phanorods with the amino groups of Pol-K through EDC chemistry ^22^. A total of 10^14^ phanorods were reacted with 1 mM EDC, 1 mM NHS, and 1 mM Pol-K (Zn^2+^ binding peptide) in a volume of 2 mL PBS buffer (pH 7.9) with gentle stirring at room temperature. The same amount of EDC was added three more times at time intervals of 30 min. The reaction was run overnight and the phanorod- Zn product purified by dialysis through regenerated cellulose dialysis tubing (molecular weight cutoff of 3500 Da) in PBS buffer. The concentration of the thiol groups from chemically conjugated Pol-K was quantified with Ellman’s assay using a cysteine as a standard (Figure S27a) and phanorods as control ^22, 111^. The number of Pol-K per phage was estimated by dividing the amount of Pol-K (concentration of thiol groups divided by 4 cysteines per Pol-K) by the number of phage particles, as quantified by real-time PCR (Figure S4c) ^22^. The Zn^2+^ ions were loaded by incubating the phanorod-Zn (1 mL of 10^14^/mL) with 1 mM ZnCl_2_ solution overnight by gentle rotating. The product was collected after centrifugation (12,000 rpm x 30 min) and washing 3x with water.

### Zn^2+^ release experiments

The release behavior of Zn^2+^ from phanorod-Zn (10^14^/mL) was investigated in PBS buffer (pH 7.2) with or without laser irradiation for 15 mins. The samples were incubated at 37°C after laser treatment and the released Zn^2+^ ions at different time points were collected by high-speed centrifugation (12000g for 30 min) for ICP-MS analysis.

### Inductively coupled plasma mass spectrometry (ICP-MS) measurement

ICP- MS analysis was performed with the NexION 2000 (PerkinElmer, Inc.). Each sample was transferred to a clean Teflon vessel for acid digestion. Digestion was carried out with a mixture of concentrated HNO_3_ (65-70%, Trace Metal Grade, Fisher Scientific) and HCl (35-38%, Trace Metal Grade, Fisher Scientific) in a ratio of 1:3 with a supplement of H_2_O_2_ (30%, Certified ACS, Fisher Scientific) at 200 °C for 50 min in a microwave digestion system (Titan MPS, PerkinElmer). The sample was cooled to room temperature and subsequently diluted to a final volume of 50 mL by adding filtered DI water before analysis. The calibration curve was established using a standard solution, using a dwell time of 50 ms with thirty sweeps and three replicates with background correction.

### Transmission Electron Microscopy (TEM)

TEM was performed on a Tecnai FEI T12 electron microscope (CNSI, UCLA). The samples were prepared by applying a few drops of solution onto TEM grids coated with a 20 nm-thick carbon film.

### Confocal Microscopy

Fluorescence images were obtained on a Leica SP8 confocal microscope (Leica, Germany) with excitation at 480 nm (CNSI, UCLA). The object-based colocalization was analyzed by JACoP v2.0 (ImageJ), and the number of green objects was counted (*N*_green_) and their centers identified. The number of these centers that colocalize with the center of a red object (dead cells) was counted (*N*_col_). The images were taken and analyzed by LAS X software.

### Ultraviolet−Visible Spectra Measurement

UV−vis absorbance spectra were collected on an HP 8453 UV-Visible Spectrophotometer with a quartz spectrasil UV−vis cuvette using direct detection at a slit width of 2 nm.

### Attenuated Total Reflection Infrared Spectra Measurement

The samples were completely dried with lyophilization before the measurement. ATR-FTIR spectra were measured with a Nicolet iS10 FTIR using a MCT detector and a Harrick Scientific Corporation GATR accessory.

### Photothermal performance measurements

The photothermal stabilities of AuNRs, phanorods and phanorod-Zn were analyzed by irradiating the samples with NIR laser (1.2 W cm^−2^) for 6 min and the samples were allowed to cool. This cycle was repeated for another four times. The photothermal conversion efficiency (*η*) of AuNR, phanorods and phanorod-Zn was calculated by the following equation (more details in Supporting Information) ^112–113^:

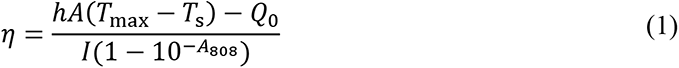

where *h* is the heat transfer coefficient; *A* is the surface area of the container; *T*_max_ is the maximum temperature reached; *T*_S_ is the surrounding temperature; *Q*_0_ is the heat input due to light absorption by the solvent (measured as 12.9 mW independently, using a quartz cuvette cell containing pure water). *I* is the power density of the laser applied (1.2 W cm^−2^), and *A*_808_ is the absorbance of the sample at 808 nm.

Here *hA* can be calculated by the following equation,

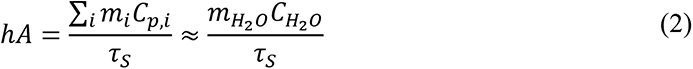

where *τ*_S_ is the system time constant; *m*_H_2_O_ and *C*_H_2_O_ are the mass of the water used as solvent (1.0 g) and specific heat capacity of water (4.2 J/g), respectively. In the natural cooling period, *τ*_S_ can be calculated using the linear regression curve of the equation below:

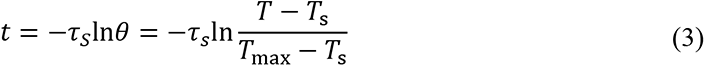

### Evaluation of antibacterial effect *in vitro*

The *in vitro* antibacterial activity of phanorods and phanorod-Zn against *P. aeruginosa*, *V. cholerae* and *E. coli* was quantitatively assessed by determination of colony-forming units (CFU). 0.5 mL of bacterial suspension (1 × 10^8^ CFU/mL) was incubated with PBS (control), phanorod (10^14^ phages/mL) or phanorod-Zn (10^14^ phages/mL) in quartz cuvette cells with or without 808 nm laser irradiation (0.3 W cm^−2^) for 15 min. After the laser treatments, the sample was diluted 1000-fold and 10 μL was inoculated on LB plates and incubated at 37 °C for 24 h. The CFU were counted and the survival rate of bacteria calculated.

Additionally, live/dead staining was performed to assay bacteriocidal activity, according to the manufacturer’s protocol. The samples were stained with a BacLight™ bacterial viability kit (Thermo Fisher Scientific) and imaged on Leica SP8 confocal microscope. The antibody- AuNRs against *P. aeruginosa* were prepared according to previous report.^24^ The *in vitro* antibacterial ability of antibody-AuNRs was evaluated using the same method as described above for phanorods. The concentration of the conjugated AuNRs was adjusted to be the same as phanorod and phanorod-Zn ([Au] = 3.3 μM).

### Evaluation of *in vitro* ablation effect of *P. aeruginosa* biofilm

The *P. aeruginosa* biofilm was prepared using a protocol modified from the literature ^114^. A single colony of *P. aeruginosa* was used to inoculate LB and incubated in a shaker-incubator overnight at 37 °C. The culture was diluted 100-fold into fresh medium and 150 μL of the dilution was added to Lab-Tek plates (culture area, 0.7 cm^2^) for overnight incubation at 37 °C. Then the liquid was removed by turning the plate over and shaking. The remaining biofilm was washed two times by submerging the plate in a small tub of water and shaking out the water. 300 μL of phanorod or phanorod-Zn (10^14^ phages per mL, 3.3 μM Au) or antibody-AuNRs (3.3 μM Au, ∼2×10^15^ AuNRs) were added to the biofilm and incubated for 30 min. Unbound bioconjugates in suspension were removed by pipetting. The biofilm was irradiated with the NIR laser for 15 min as described above. Cell viability was studied by resuspending the bacteria and growing aliquots on LB plates (1 μL of cell suspension was diluted in 1 mL of PBS buffer, and 5 μL of the dilution was plated) for colony counting, and by confocal microscopy with live/dead cell viability staining with SYTO9 and PI (Filmtracer™ LIVE/DEAD™ biofilm viability kit, Thermo Fisher Scientific).

### Mouse model of *P. aeruginosa*-infected wounds

All animal procedures were conducted in accordance with institutional guidelines and approved by the UCLA Institutional Animal Care and Use Committee (Protocol ARC-2020-044). 12-15 weeks-old male and female mice (22-25g, strain 027-C57BL/6 from Charles River Laboratories, MA, USA) were randomly assigned to different groups. Wound exposure to PBS buffer was used as the negative control group. Phanorod and phanorod-Zn treatments were used in the experimental groups, and antibody- AuNRs were used as a comparison group. The following standard clinical treatment methods were applied as positive control groups: two systemic antibiotics (ciprofloxacin and levofloxacin, orally fed), one topical antibiotic (polysporin), and two topical antiseptics (acetic acid and chlorhexidine).

On day 0, the mice were anaesthetized by isoflurane vaporized in O_2_ (2.5%) prior to surgery and placed upon a sterile towel on a warm pad with circulating water. The dorsal side was shaved, depilated and then disinfected with ethanol pads and povidone iodine swabs. An artificial wound ∼6 mm long (or ∼12 mm long for the large wound group) was made on the dorsum of each mouse with a scissor, resulting in an initial wound area of ∼ 8.5 mm^2^ (or ∼17.6 mm^2^ for the large wound group). Wounds were inoculated with 20 or 100 μL of *P. aeruginosa* (5×10^8^ cfu/mL) and incubated for 1 h.

For wound treatment by PBS (control), phanorod (10^14^/mL in PBS), phanorod-Zn (10^14^/mL in PBS), or antibody-AuNR (3.3 uM in PBS), a particle suspension (∼50-100 μL) was applied and incubated for 30 min, followed by 808 nm laser irradiation (0.3 W cm^−2^, unless otherwise specified) for 15 min. Unless otherwise specified, phanorod and phanorod-Zn treatment (reagent application followed by irradiation) was performed on day 0, with a second round of irradiation also applied on day 1. Alternative treatment regimens included treatment on day 2 with a second round of irradiation on day 3, or single treatment on day 0. For systemic antibiotics, treatment was applied by feeding the antibiotics orally (250 mg kg^−1^ day^−1^). The topical antibiotic and antiseptic treatments were applied by covering the wound area with polysporin ointment (Johnson & Johnson Consumer Inc.), 4% chlorhexidine (McKesson Corporation) or 2% acetic acid (Akorn Pharmaceuticals). The antibiotics and antiseptics were applied every 24 hours unless stated otherwise.

To evaluate the therapeutic effect, wounds were photographed (by iPhone 8) at regular time intervals and wound size was measured by a ruler. For analysis of CFUs, mice were sacrificed and the wound tissue harvested, weighed, chopped and homogenized in PBS buffer. The homogenized samples were centrifuged at 3000 g for 6 min to pellet debris, and 100 μL of supernatant was plated onto LB agar for CFU counting. For histomorphological analysis, wound tissue was fixed with 4% paraformaldehyde solution and washed with 75% ethanol. The tissue samples were analyzed by hematoxylin and eosin staining ^115^ or Masson’s trichrome staining ^116^ (performed by the Translational Pathology Core Laboratory (TPCL) at UCLA). The relative collagen area was determined using ImageJ. The area of collagen was measured by manual adjustment of a threshold value in the blue channel to match visual inspection. Similarly, the area of the tissue was measured by manual adjustment of the threshold in all color channels to match visual inspection. To assess toxicity, the major organs (heart, liver, spleen, lung and kidney) of the mice were stained by hematoxylin and eosin on day 10, and blood and serum were collected for biochemical analysis (performed by IDEXX Laboratories) on day 10. The concentrations of Zn^2+^ and Au^3+^ ions were measured by ICP-MS as described above.

To evaluate delayed treatment by phanorod-Zn, an experimental group was treated with phanorod-Zn on day 2 and 3 (phanorod-Zn was applied only on day 2 and irradiated with laser on day 2 and 3), and compared to a control group with ciprofloxacin treatment starting from day 2.

No unexpected or unusually high safety hazards were encountered.

### Statistical analysis

All the quantitative data in each experiment are presented as mean ± standard deviation of at least three independent experiments. Student’s *t* test (two-sided) was utilized to evaluate the statistical significance. Values of *p* < 0.05 (*), *p* < 0.01 (**) and *p* < 0.001 (***) were considered statistically significant.

### Data availability

All data generated or analyzed during this study are included in this manuscript and its supplementary information file.

## Acknowledgements

The authors thank J. Li for provision of the *P. aeruginosa* PAK*pmrB6* strain. The authors acknowledge the use of ICP-MS facility within the UC Center for Environmental Implications of Nanotechnology in CNSI at UCLA. The Leica SP8 confocal laser scanning microscopy was performed at the Advanced Light Microscopy/Spectroscopy Laboratory and the Leica Microsystems Center of Excellence at the California NanoSystems Institute at UCLA with funding support from NIH Shared Instrumentation Grant S10OD025017 and NSF Major Research Instrumentation grant CHE-0722519. The authors acknowledge the use of instruments at the Electron Imaging Center for NanoMachines supported by NIH (1S10RR23057 to ZHZ) and CNSI at UCLA. Funding is acknowledged from the National Institute of General Medical Sciences (Grant DP2GM123457 to IAC), Camille Dreyfus Teacher-Scholar Program (IAC), Simons Foundation (290358FY18 to SSM) and the Natural Sciences and Engineering Research Council of Canada (NSERC) [RGPIN-2020-04375 to SSM].

## Author contributions

HP and IAC conceived of the project and designed experiments. HP conducted experiments and analyzed data. IAC supervised the project direction. DR and SSM developed the Pol-K peptide. HP and MCJ conducted animal experiments. KPR supervised animal experiments. HP and IAC wrote the manuscript with significant input from all authors.

## Supporting Information available

Figures S1-S33, Supporting Methods, Supporting References.

## TOC graphic

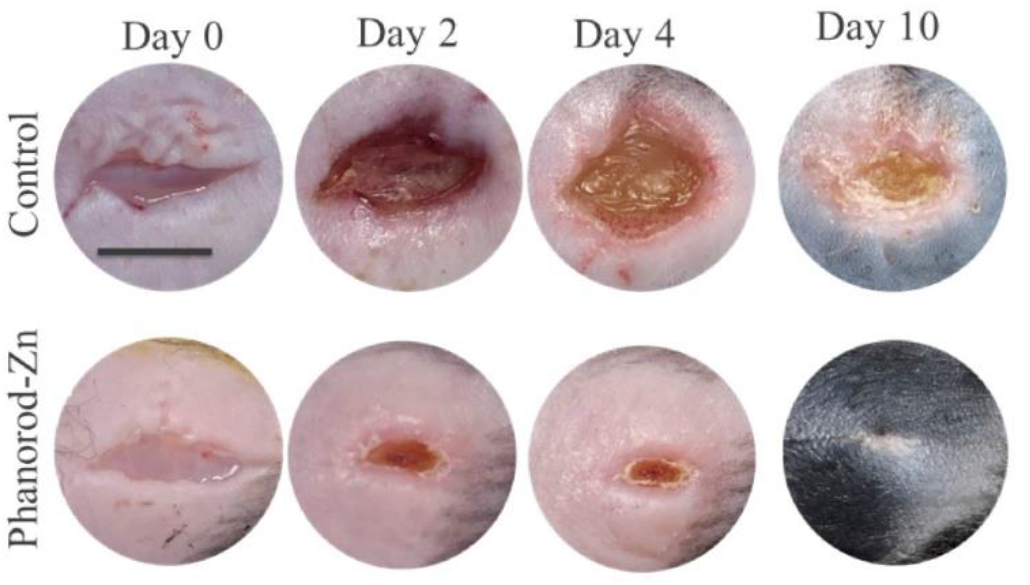

**Synopsis**: Antibiotic-resistant bacteria are a major threat to modern medicine. Zinc-loaded phages conjugated to gold nanorods are an effective, non-toxic treatment for bacterial infections in mice.

