## Supporting Information for "Treatment of wound infections in a mouse model using Zn^2+^-releasing phage bound to gold nanorods"

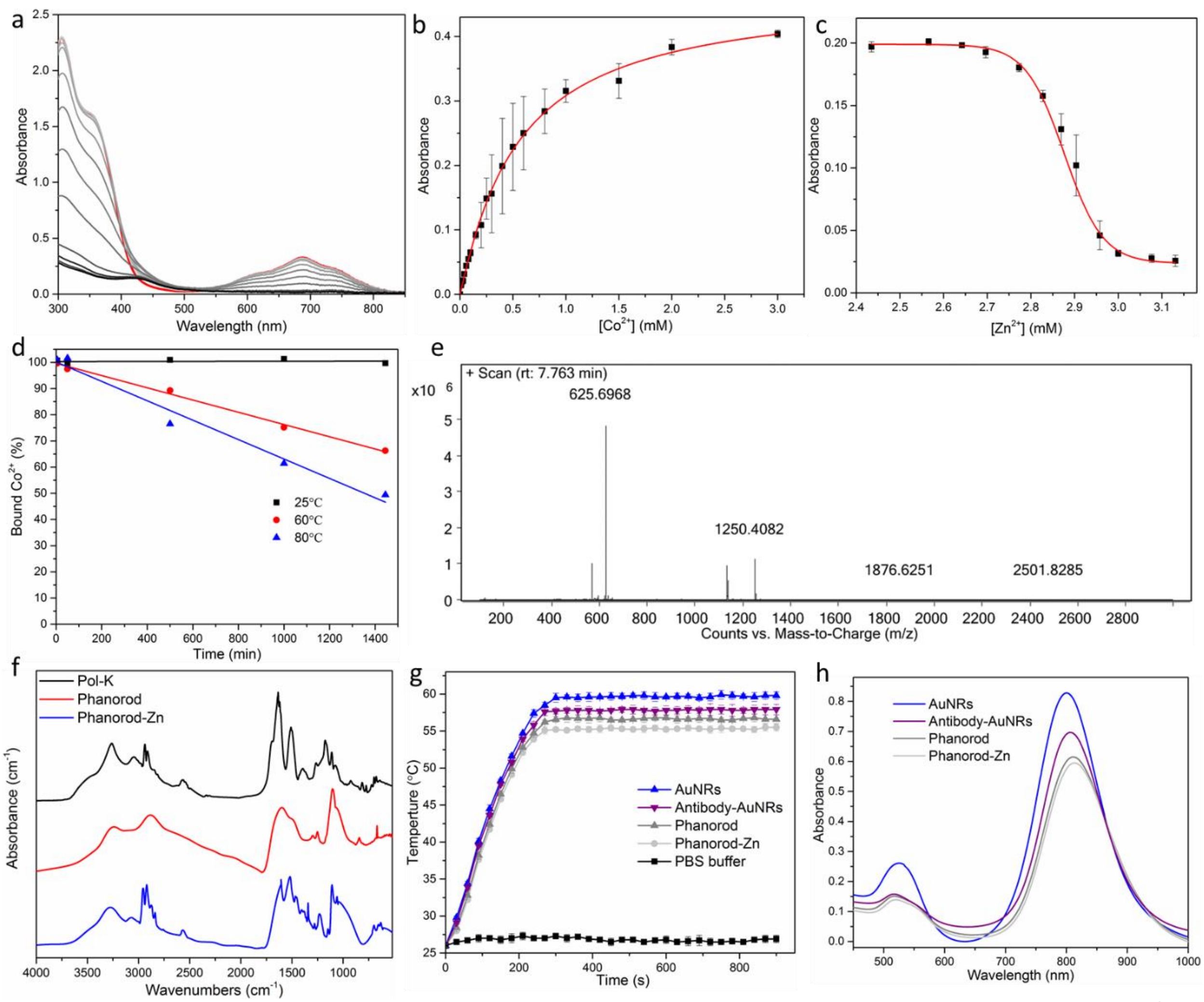

**Figure S1.** Characterization of Pol-K and phanorod-Zn. (a) Competitive binding of Pol-K-$Co^{2+}$ complex (red), with displacement by addition of aliquots of $Zn^{2+}$ (black). $Zn^{2+}$ competes with $Co^{2+}$ for binding to the deprotonated Pol-K thiols, causing the progressive disappearance of the typical S→$Co^{2+}$ electronic transitions, particularly the absorbance peak near 750 nm. (b) Saturation binding curve of $Co^{2+}$ titrated into 1 mM Pol-K peptide solution. The Hill equation fit gives a $K_d$ of 0.53 mM. (c) Competition binding curve of $Zn^{2+}$ titrated into a preformed Pol-K-$Co^{2+}$ complex (as seen in part a). The calculated $K_d$ for $Zn^{2+}$ was 43 μM. Solutions were buffered with 20 mM glycylglycine, pH 8.7. For (b) and (c), absorbance change was monitored at 750 nm, and values shown are the average of three independent experiments (plotted as mean ± standard deviation). (d) $Co^{2+}$ release from Pol-K measured at different bulk temperatures. While the absolute rate of ligand release of $Co^{2+}$ is generally slower than that of $Zn^{2+}$, the trend over varying temperatures is expected to be analogous. (e) High-resolution mass spectrum of the synthesized Pol-K peptide (GCFCEDACDKCG) in aqueous solution. The peak found at 1250.4082 m/z is consistent with the value of the isotopic mass of the protonated molecule [M+1] (chemical formula: $C_{47}H_{71}N_{13}O_{19}S_4$, exact mass: 1249.39). The peak at m/z 625.6968 is the peptide with 2 charges on it. (f) FTIR spectra of the $Zn^{2+}$-binding peptide Pol-K, phanorods and phanorod-Zn. The signals at 1510 and 3036 $cm^{-1}$ (from the C-C and C-H stretching, respectively, of the phenyl group of the Pol-K) and at 2550 $cm^{-1}$ (from the S-H stretching of the thiol groups) appeared after conjugation to synthesize phanorod-Zn, indicating successful conjugation of Pol-K to phanorods. (g) Photothermal heating curves of AuNRs, phanorods, phanorod-Zn, and antibody-AuNR solutions ([Au] = 3.3 μM) and the control (PBS buffer) under 808 nm laser irradiation (0.3 W $cm^{-2}$). (h) UV-Vis spectra of AuNRs, phanorods, phanorod-Zn, and antibody-AuNRs.

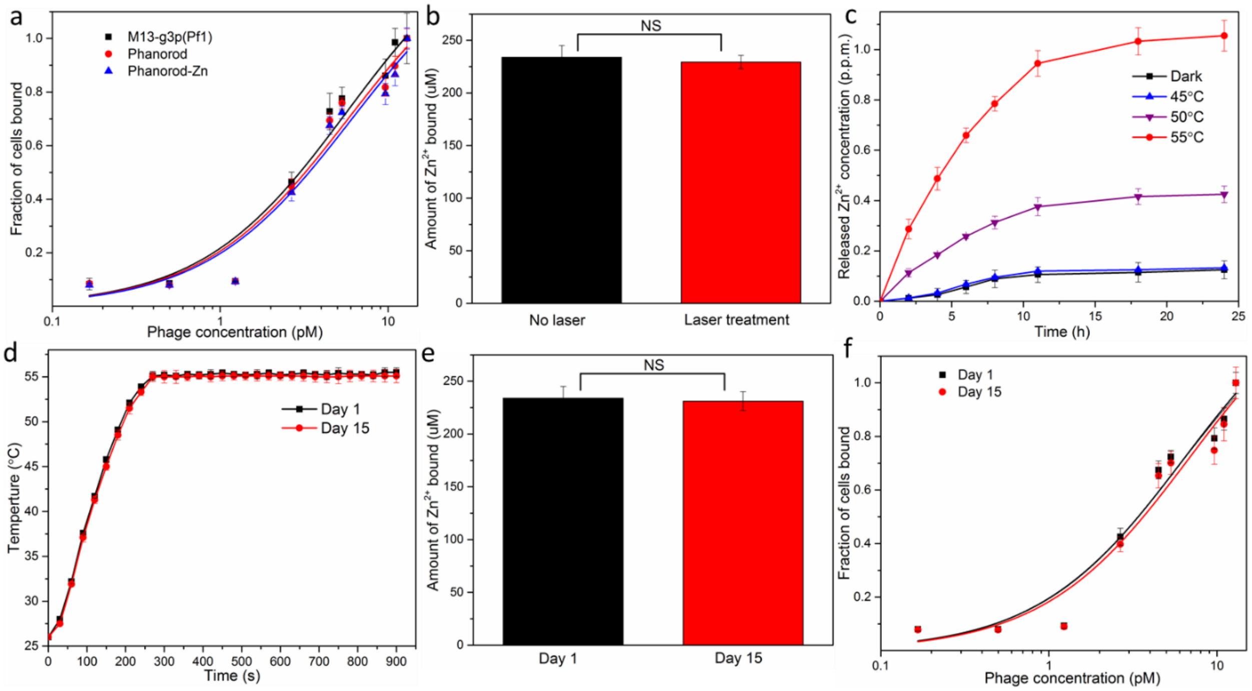


**Figure S2.** (a) Binding curves for M13-g3p(Pf1) (black, $K_d$ ≈5.4 pM), phanorod (red, $K_d$ ≈ 5.8 pM) and phanorod-Zn (blue, $K_d$ ≈ 6.1 pM), demonstrating similar affinity to *P. aeruginosa* (ATCC25102 (Schroeter) Migula). (b) Loading of phanorod-Zn ($10^{14}$ particles/mL with [Pol-K] = 294 μM) with $Zn^{2+}$ ions with (red) and without (black) a single laser irradiation for 15 min (bulk temperature: 55°C). (c) Release of $Zn^{2+}$ ions over time by phanorod-Zn, in PBS buffer after 15 min laser irradiation, at varying laser flux resulting in different bulk temperatures as given. (d-f) Stability test of phanorod-Zn after storage, showing little change in photothermal stability (d), $Zn^{2+}$-loading capacity (e), and host cell binding affinity ($K_d$ ≈ 6.8 pM on day 15) (f). Error bars are standard deviation (n = 5 independent samples); NS = not significant ($p > 0.05$, two-sided *t* test).

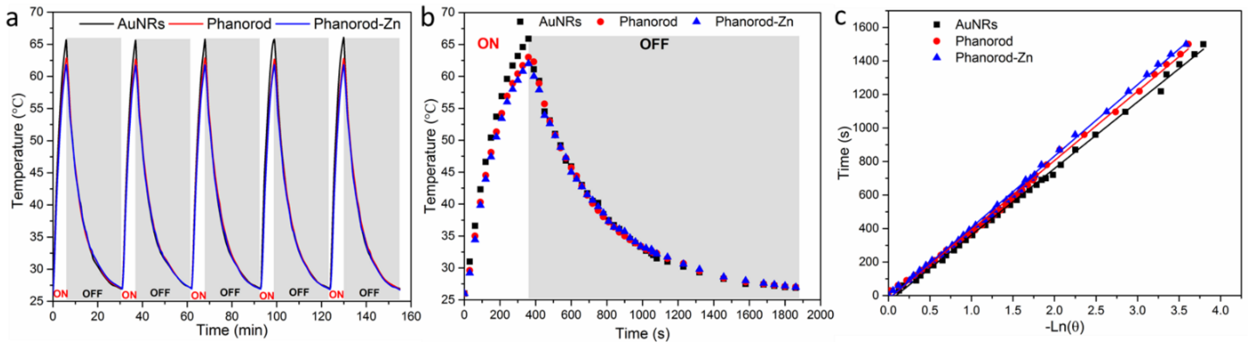


**Figure S3.** Photothermal reversibility and stability. (a) Cyclic heating curves of AuNRs, phanorods, and phanorod-Zn solutions ([Au] = 3.3 μM) under 808 nm laser irradiation (1.2 W $cm^{-2}$) for five on/off cycles (6 min on, 25 min off), as indicated. (b,c) Calculation of the photothermal conversion efficiencies (η) of phanorod-Zn, phanorods and AuNRs at 808 nm (see Methods for calculation details).

The time constants for heat transfer from the systems were determined to be $\tau_s$ = 395, 415, and 426 s for AuNRs, phanorods and phanorod-Zn, respectively, from fitting the linear time data from the cooling period versus negative natural logarithm of driving force temperature (see Equation (3) in Methods) from Figure S3c. The $hA$ is calculated by Equation (2) where $m_{H2O}$ and $C_{H2O}$ are the mass of the water used as solvent (1.0 g) and specific heat capacity of water (4.2 J/g). In Equation (1), the $T_{max}$ is the maximum temperature reached (for AuNRs, phanorod and phanorod-Zn, $T_{max}$ = 65.9, 63 and 62 °C, respectively); $T_S$ is the surrounding temperature (26°C); $Q_0$ is the heat input due to light absorption by the solvent (measured as 12.9 mW independently, using a quartz cuvette cell containing pure water). $I$ is the power density of the laser applied (1.2 W $cm^{-2}$), and $A_{808}$ is the absorbance of the sample at 808 nm (for AuNRs, phanorod and phanorod-Zn, $A_{808}$ = 0.71, 0.59 and 0.57, respectively; see Figure S1g). With the above data, the photothermal conversion efficiencies (η) of AuNRs, phanorods, and phanorod-Zn, at 808 nm can be calculated as 42.6%, 40.6% and 38.9% according Equation (1). The calculated photothermal conversion efficiencies (η) are comparable with previously reported values[1-3]. The slightly diminished photothermal conversion efficiencies (η) of phanorods and phanorod-Zn compared with AuNRs is probably due to the enhanced scattered light by the phage and peptide[4]. However, this influence does not represent a major effect on the capacity of AuNRs to transform the electromagnetic radiation into heat. A similar phenomenon was observed in previous literature[5].

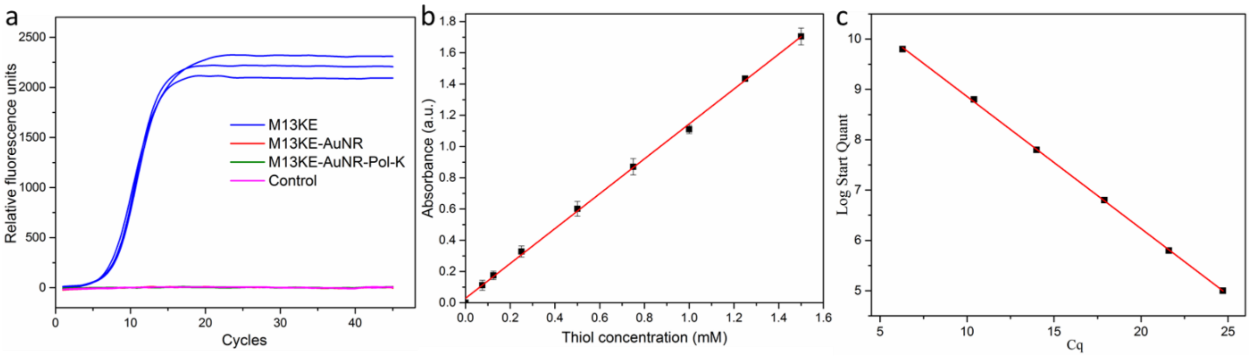


**Figure S4.** (a) Quantitative PCR amplification of samples using M13KE without NIR irradiation (blue), or using M13KE-AuNR (red) or M13KE-AuNR-Pol-K (green) bioconjugates with 15 min NIR irradiation followed by phage propagation. The control sample (magenta) included water instead of template DNA. The results indicate that no detectable phage resulted from attempted propagation of the irradiated M13KE-AuNR or M13-AuNR-Pol-K sample. (b) Standard curve of Ellman's assay for quantifying thiol group using absorbance measurement at 412 nm. Fitting equation: $y = 1.11684x+0.0278$, $R^2 = 0.9989$. Error bars are standard deviations ($n = 3$ independent samples). (c) Standard curve of real-time PCR of phanorod-Zn particles (amount given as log(# phanorod-Zn particles)). Fitting equation: $y=-0.26249x+11.48504$, $R^2=0.9997$.

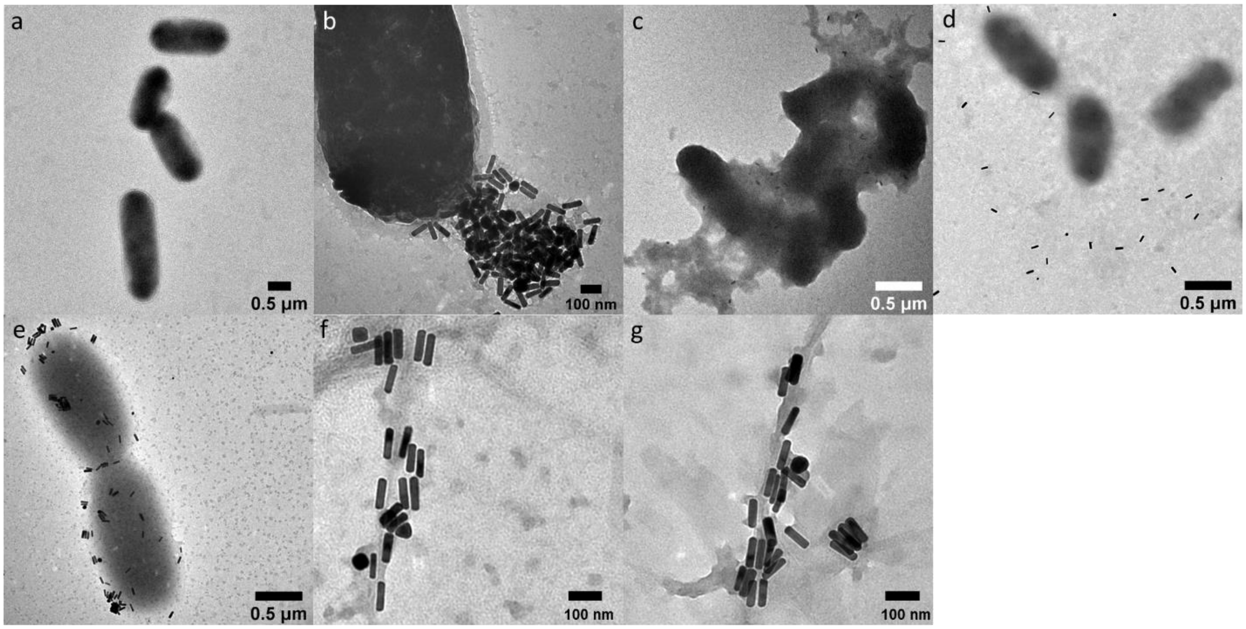


**Figure S5.** TEM morphology of *P. aeruginosa* with no treatment (a) and treated by 808 nm NIR laser irradiation (0.3 W cm$^{-2}$, 15 min) with phanorod-Zn (b, before irradiation; c, after irradiation) shows attachment of phanorod-Zn and grossly altered morphology and loss of cytoplasmic contents after photothermal treatment.

The anti-*P. aeruginosa* antibody also directs AuNRs to the cells, as shown by TEM images of *P. aeruginosa* mixed with unconjugated AuNRs (coated with PEG-COOH, d) and antibody-AuNRs (e). The TEM morphology of phanorods (f) and phanorod-Zn in dry state (g), shows the association of the conjugated AuNRs with the filamentous phages.

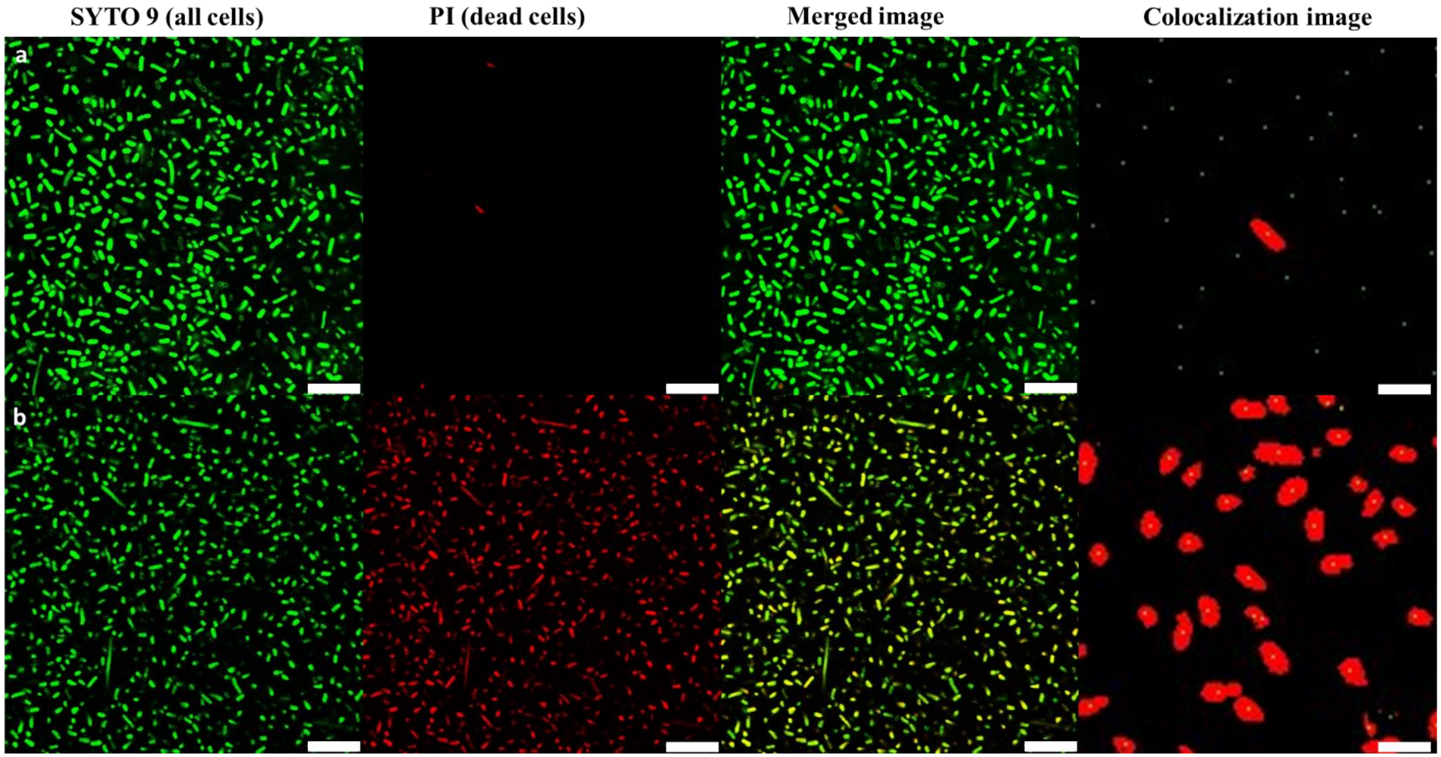


| | $N_{green}$ | $N_{col}$ | $f_{col}$ |
|---|---|---|---|
| a | 586 | 2 | 0.00341 |
| b | 627 | 620 | 0.988 |

**Figure S6.** Live/dead staining of *P. aeruginosa* cells in suspension before (row a) and after (row b) 15 min NIR irradiation (808 nm, 0.3 W cm$^{-2}$) with phanorod-Zn. From left to right, panels are green field (SYTO9: staining both live and dead cells), red field (PI: staining dead cells), merged image of the two fields, and representative colocalization image. Scale bar in the colocalization images = 2.5 μm. The other scale bars are 10 μm.

In object-based colocalization analysis by JACoP v2.0 (ImageJ), the number of green objects (all cells) was counted ($N_{green}$) and their centers identified. The number of these centers that colocalize with the center of a red object (dead cells) was counted ($N_{col}$). $f_{col}$ is the fraction of cells identified as dead by this colocalization analysis ($N_{col}/N_{green}$). The colocalization image shows the green centers (appearing as dots) as well as the red objects (thresholded).

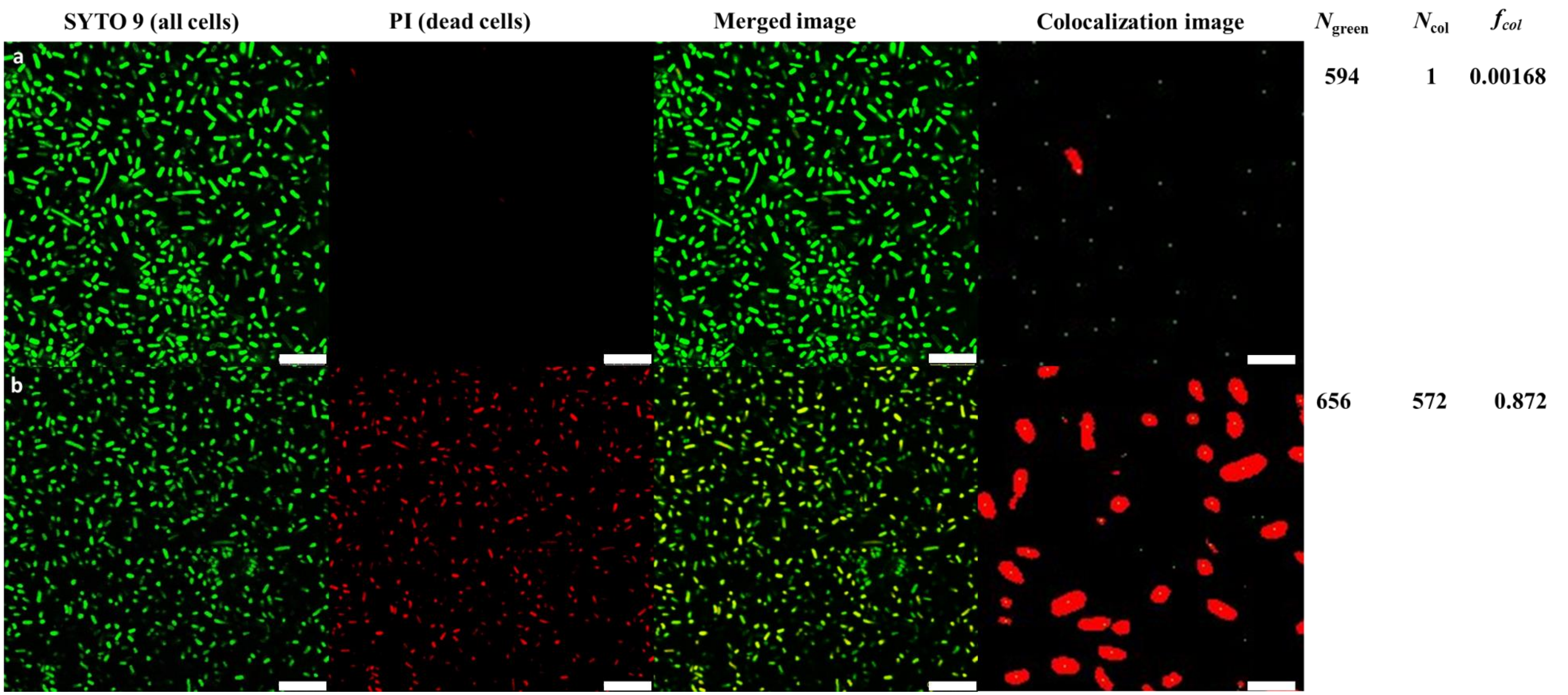


**Figure S7.** Live/dead staining of *P. aeruginosa* cells in suspension before (row a) and after (row b) 15 min NIR irradiation (808 nm, 0.3 W $cm^{-2}$) with phanorods. From left to right panels are green field (SYTO9: staining both live and dead cells), red field (PI: staining dead cells), merged image of the two fields, and representative colocalization image. Scale bar in the colocalization image = 2.5 μm. The other scale bars are 10 μm. See caption of Figure S6 for explanation of colocalization analysis.

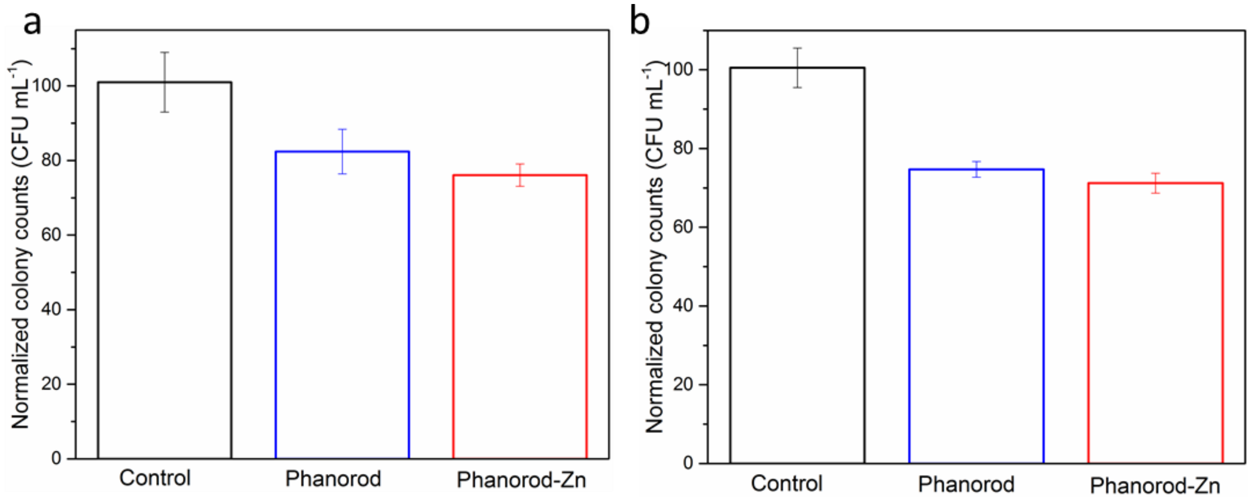


**Figure S8.** Normalized colony counts of (a) *E. coli* and (b) *V. cholerae* in suspension, with NIR laser irradiation with phanorods or phanorod-Zn. Error bars are standard deviation (n = 5 independent samples).

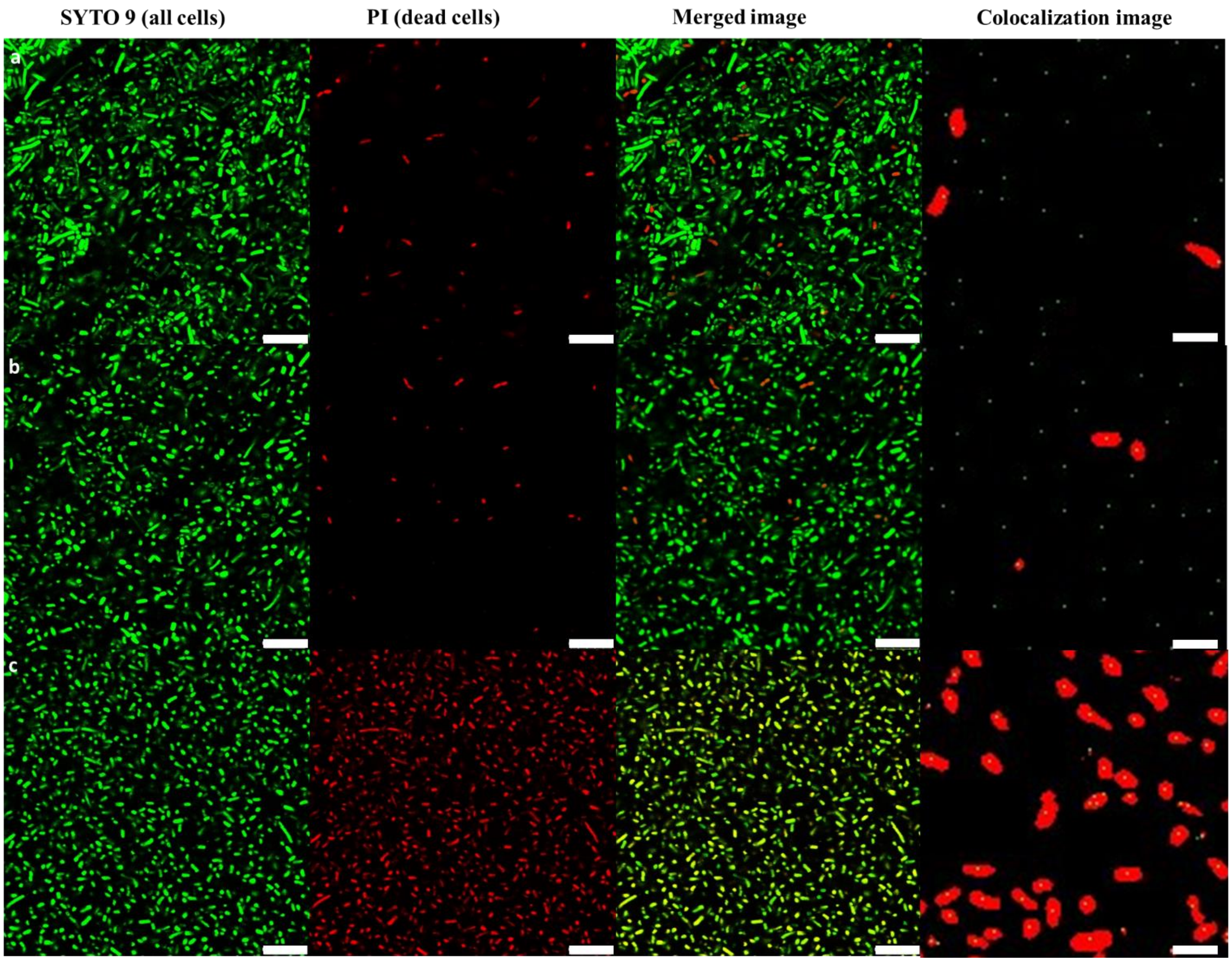


| | $N_{green}$ | $N_{col}$ | $f_{col}$ |
|---|---|---|---|
| a | 851 | 34 | 0.0399 |
| b | 702 | 26 | 0.0370 |
| c | 761 | 743 | 0.976 |

**Figure S9.** Live/dead staining of *P. aeruginosa* biofilm in PBS (row a), or before (row b) or after (row c) 15 min NIR irradiation (808 nm, 0.3 W cm$^{-2}$) with phanorod-Zn. From left to right, panels are green field (SYTO9: staining both live and dead cells), red field (PI: staining dead cells), merged image of the two fields, and representative colocalization image. Scale bar in the colocalization images = 2.5 μm. The other scale bars are 10 μm. See caption of Figure S6 for explanation of colocalization analysis.

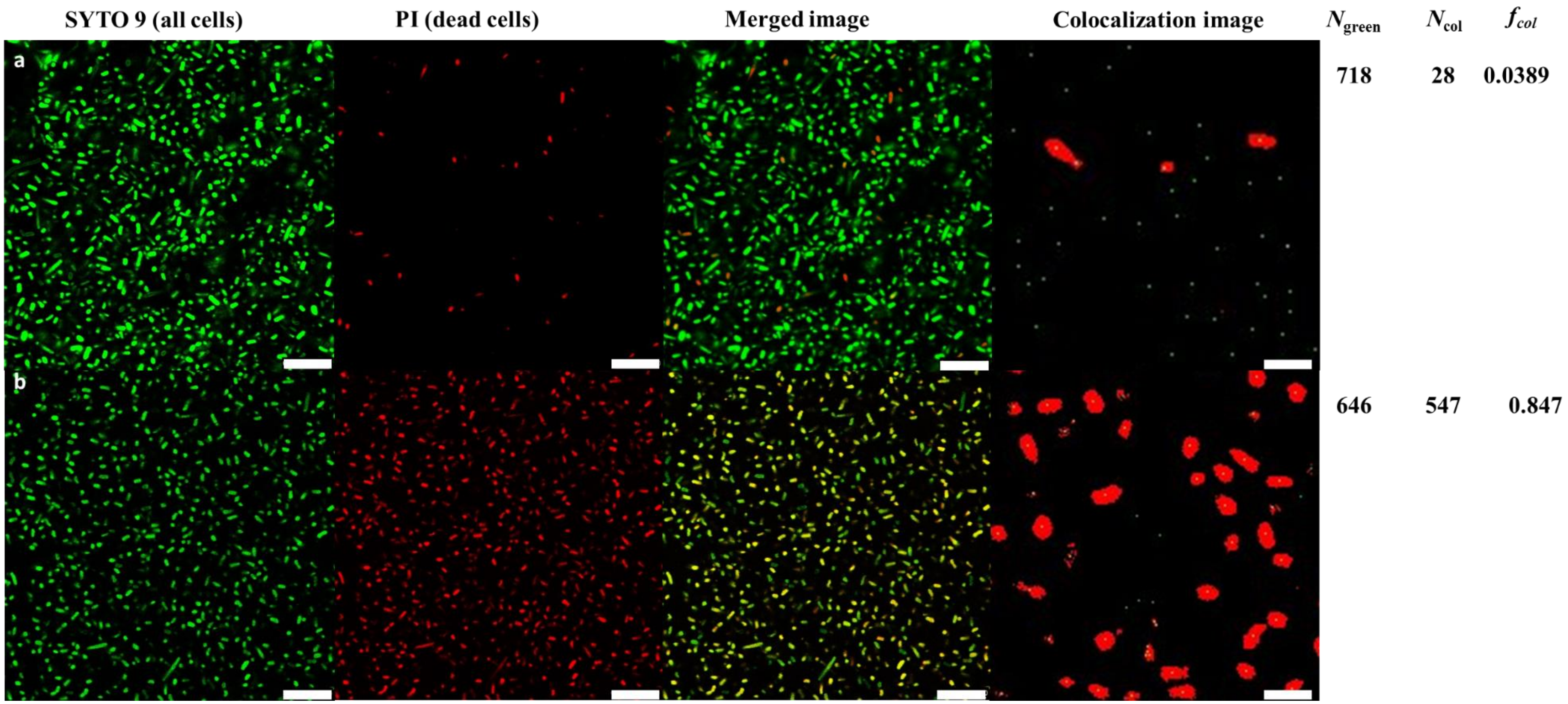


**Figure S10.** Live/dead staining of *P. aeruginosa* biofilm before (row a) and after (row b) 15 min NIR irradiation (808 nm, 0.3 W cm$^{-2}$) with phanorods. From left to right, panels are green field (SYTO9: staining both live and dead cells), red field (PI: staining dead cells), merged image of the two fields, and representative colocalization image. Scale bar in the colocalization images = 2.5 μm. The other scale bars are 10 μm. See caption of Figure S6 for explanation of colocalization analysis.

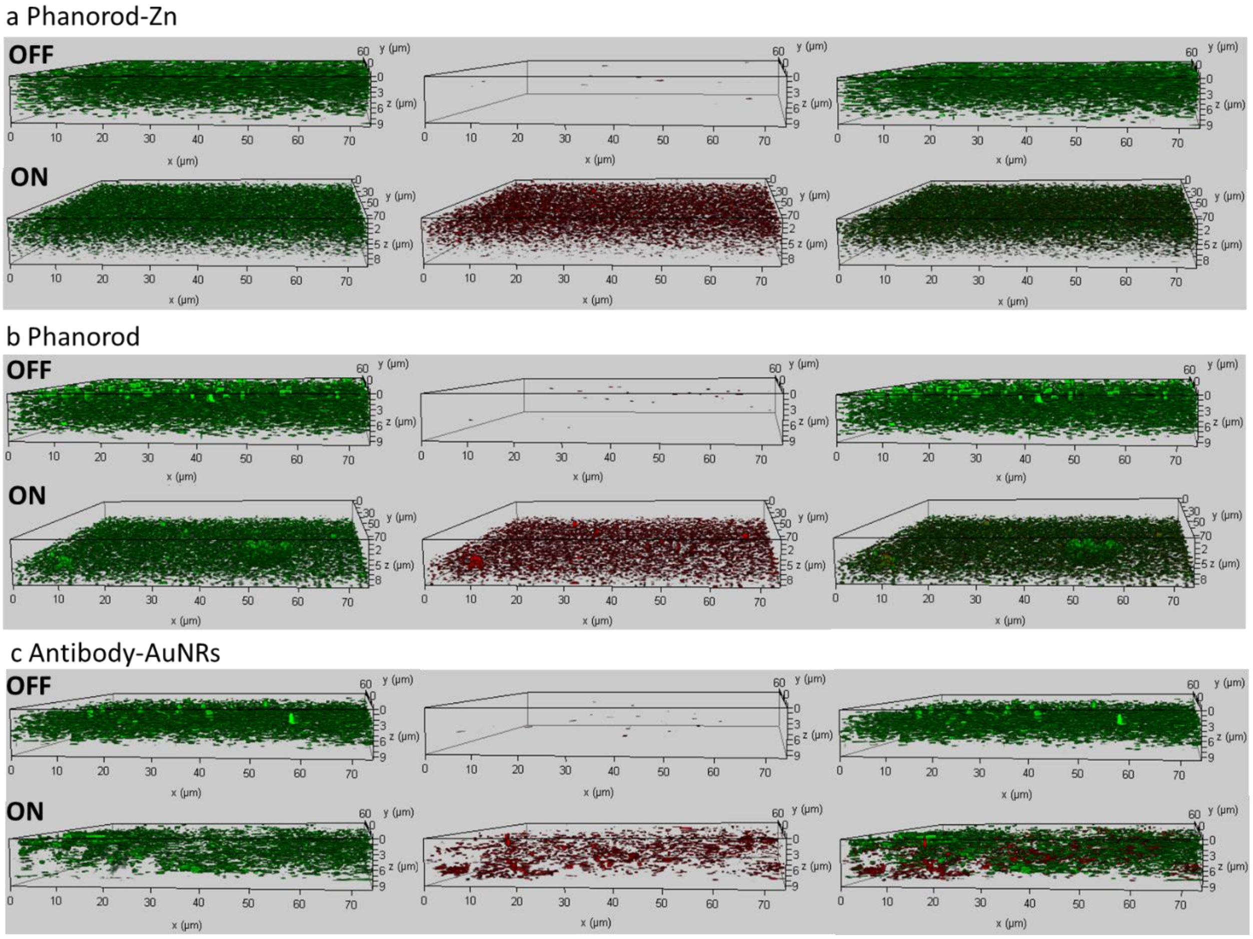


**Figure S11.** Three-dimensional reconstructed image of *P. aeruginosa* biofilm with live/dead staining, before (laser off) and after (laser on) 808 nm NIR irradiation (0.3 W cm$^{-2}$) for 15 min, with phanorod-Zn (a), phanorods (b), or antibody-AuNRs (c). From left to right, the first column shows staining by SYTO9 (green: all cells), the second column shows staining by PI (red: dead cells), and the third column shows the merged image.

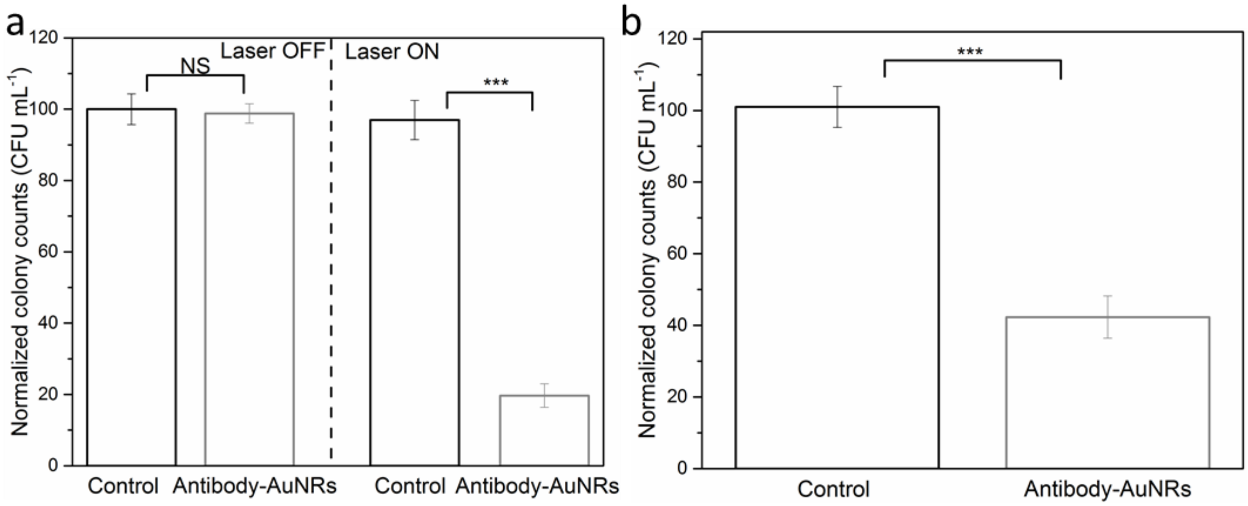


**Figure S12.** Antibacterial effect of antibody-AuNRs *in vitro*, determined by colony-forming units. (a) Normalized colony counts of *P. aeruginosa* in suspension, without or with 808 nm NIR laser irradiation (0.3 W cm$^{-2}$, 15 min) with antibody-AuNRs, as marked. The control (no antibody-AuNRs applied) indicates that antibody-AuNRs themselves did not cause cell death. (b) Normalized colony counts of *P. aeruginosa* biofilm exposed to 808 nm NIR laser irradiation (0.3 W cm$^{-2}$, 15 min) with antibody-AuNRs. Error bars are standard deviation (n = 5 independent samples); *** indicates $p < 0.001$; NS = not significant ($p > 0.05$); two-sided $t$ test.

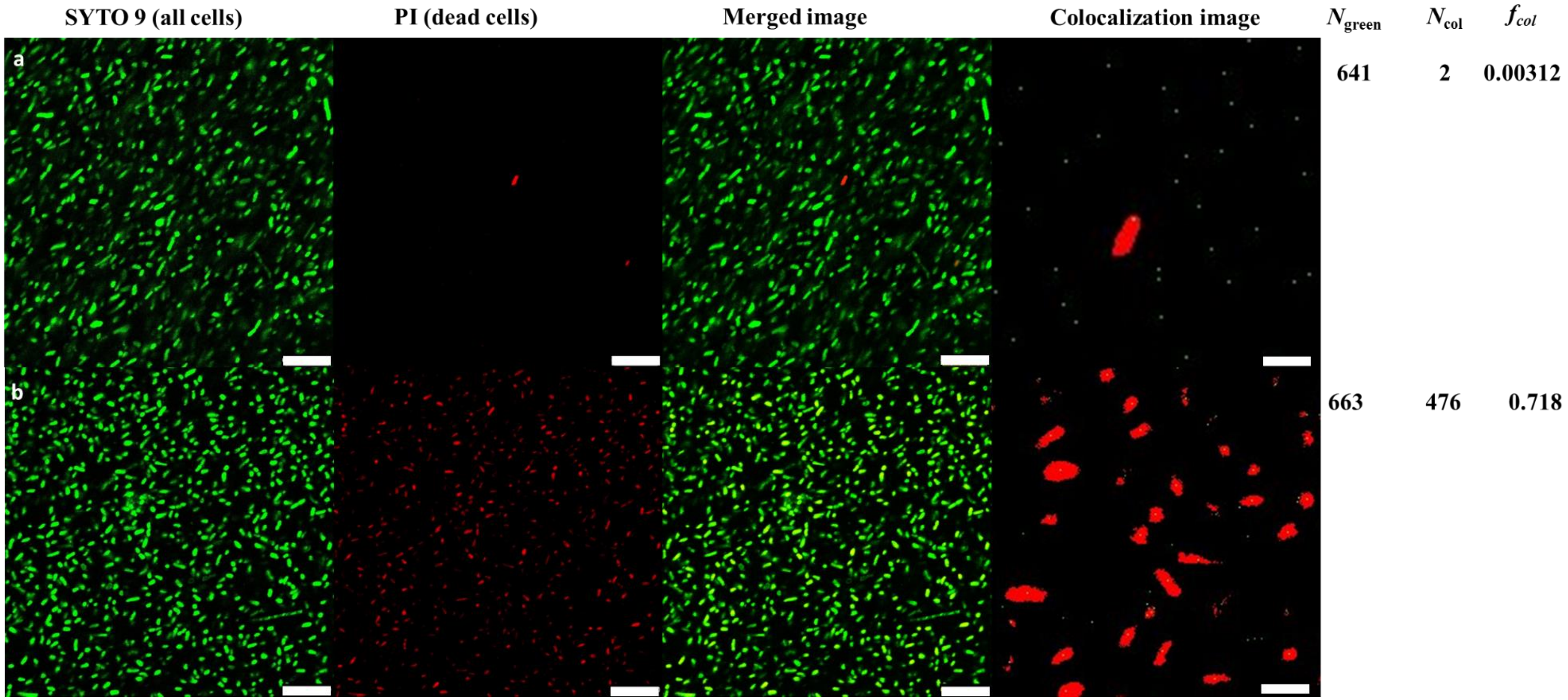


**Figure S13.** Live/dead staining of *P. aeruginosa* cells in suspension before (row a) and after (row b) 15 min NIR irradiation (808 nm, 0.3 W cm$^{-2}$) with antibody-AuNRs. From left to right: panels are green field (SYTO9: staining both live and dead cells), red field (PI: staining dead cells), merged image of the two fields, and representative colocalization image. Scale bar in the colocalization image = 2.5 μm. The other scale bars are 10 μm. See caption of Figure S6 for explanation of colocalization analysis.

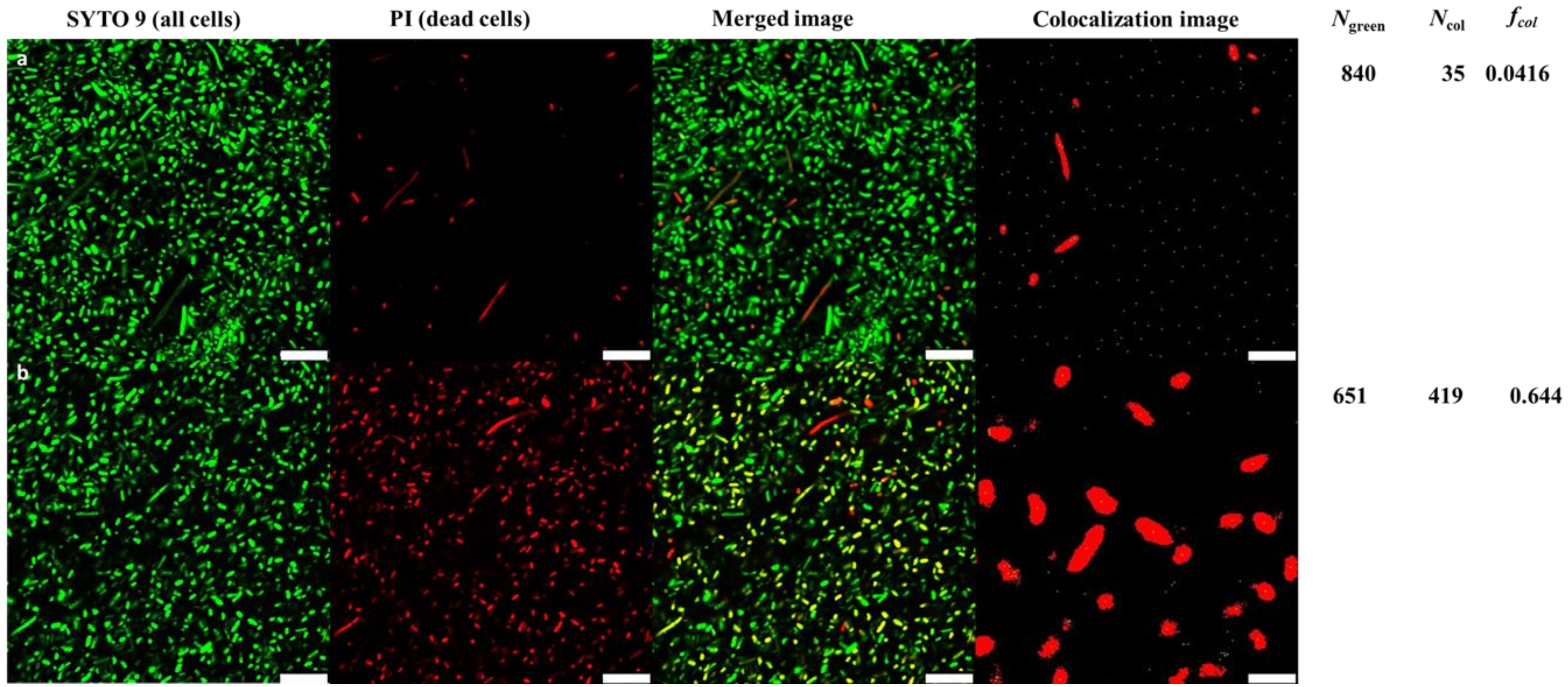


**Figure S14.** Live/dead staining of *P. aeruginosa* biofilm before (row a) and after (row b) 15 min NIR irradiation (808 nm, 0.3 W cm$^{-2}$) with antibody-AuNRs. From left to right, panels are green field (SYTO9: staining both live and dead cells), red field (PI: staining dead cells), merged image of the two fields, and representative colocalization image. Scale bar in the colocalization image = 2.5 µm. The other scale bars are 10 µm. See caption of Figure S6 for explanation of colocalization analysis.

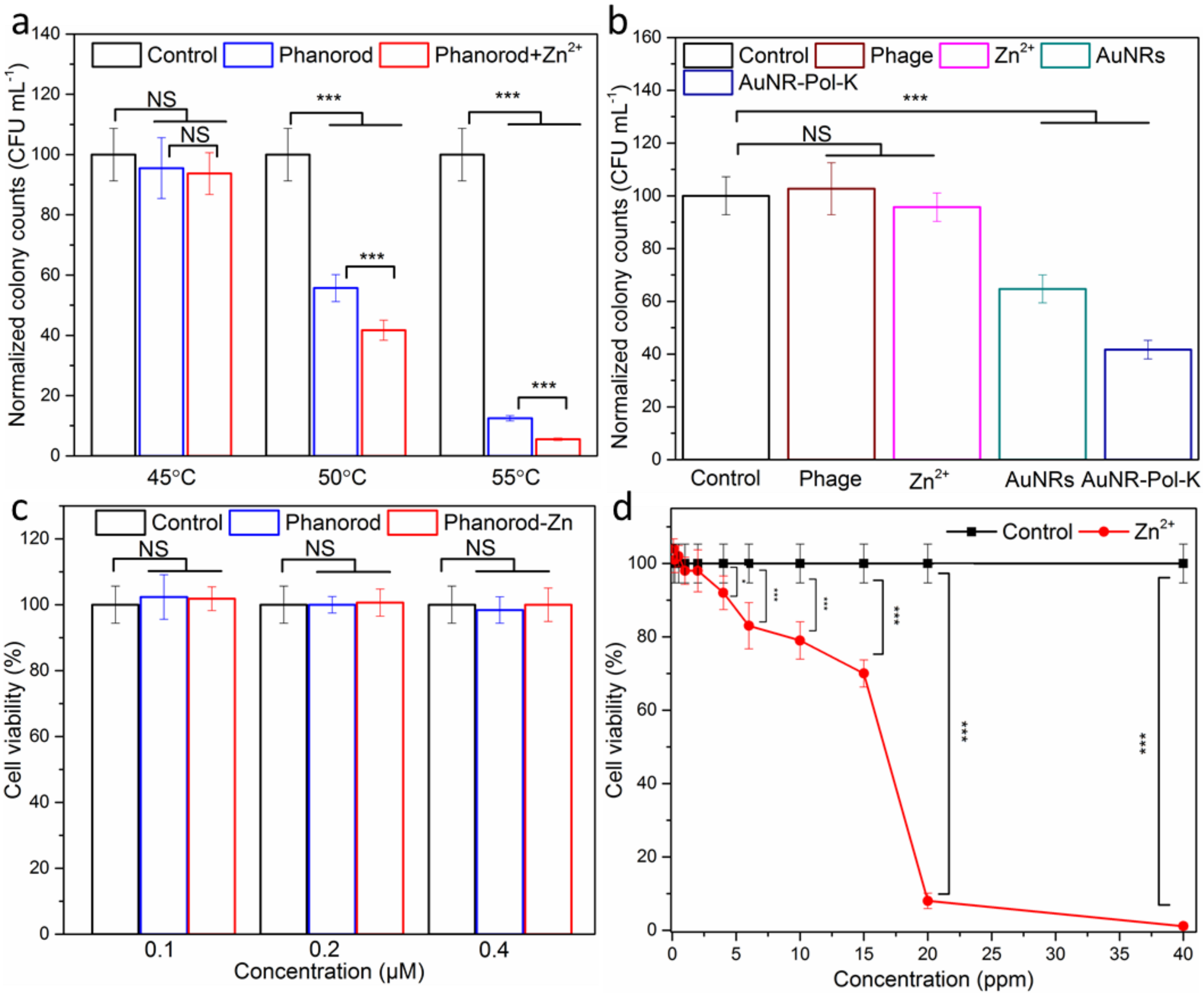


**Figure S15.** (a) Antibacterial effects of phanorod and mixture of phanorod with $Zn^{2+}$ ($Zn^{2+}$ concentration = 1.06 ppm) after laser irradiation for 15 mins, tested against *P. aeruginosa* in suspension. The bulk temperatures shown here were controlled by adjusting the laser flux. (b) Antibacterial effects of phage ($10^{14}$ particles/mL), $Zn^{2+}$ (1.06 ppm), AuNRs and AuNR-Pol-K, tested against *P. aeruginosa* in suspension. The AuNRs and AuNR-Pol-K were treated with laser irradiation for 15 mins to achieve bulk temperature of 55 °C. (c, d) The *in vitro* cytocompatibility study of (c) phanorod/phanorod-Zn and (d) $Zn^{2+}$ with HEK293T cells at different concentrations. In part (c), the *x*-axis concentration is given for Au of phanorods or phanorod-Zn. Error bars are standard deviation (n = 5 independent samples). * indicates $p < 0.05$, *** indicates $p < 0.001$, NS = not significant ($p > 0.05$), two-sided *t* test.

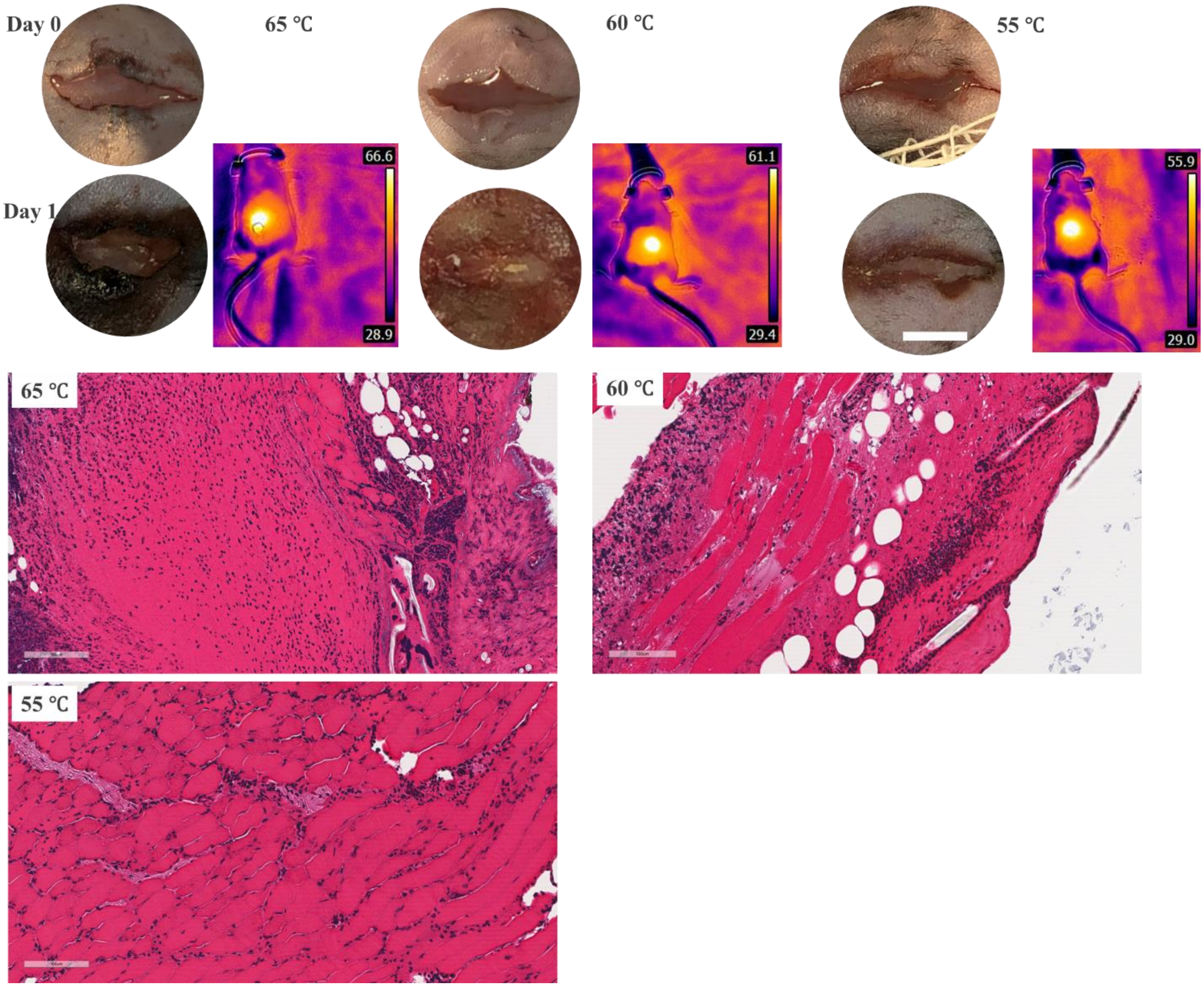


**Figure S16.** Wounds treated with phanorod-Zn with NIR laser (808 nm, 0.3-0.4 W cm$^{-2}$) for 15 min, with corresponding thermal imaging (taken during irradiation), as well as photography and H&E staining (scale bars = 100 μm) of wound tissue (taken on day 1). Wound creation, inoculation, and irradiation took place on day 0.

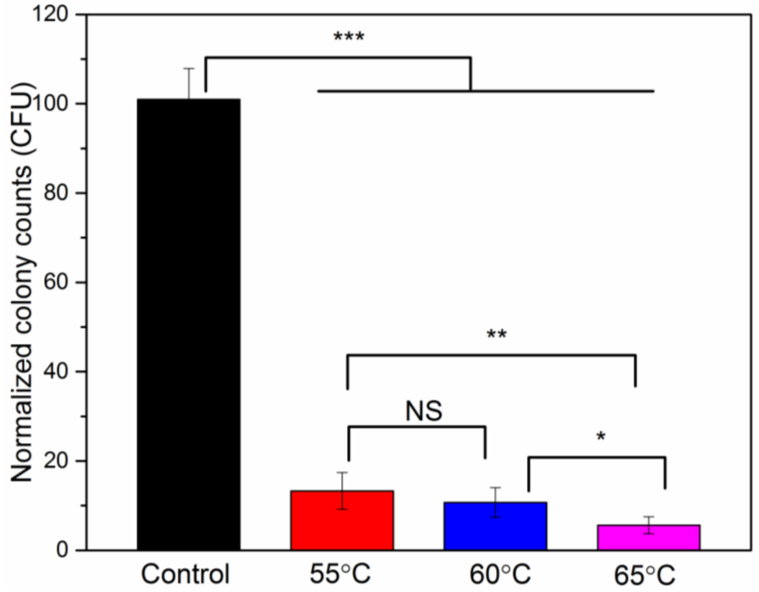


**Figure S17.** Survival of *P. aeruginosa* determined for wound tissue under exposures shown in Figure S15. The control is wound exposure to PBS buffer. Error bars are standard deviation (n = 5 mice: * indicates $p < 0.05$, ** indicates $p < 0.01$, *** indicates $p < 0.001$, NS = not significant ($p > 0.05$); two-sided *t* test.

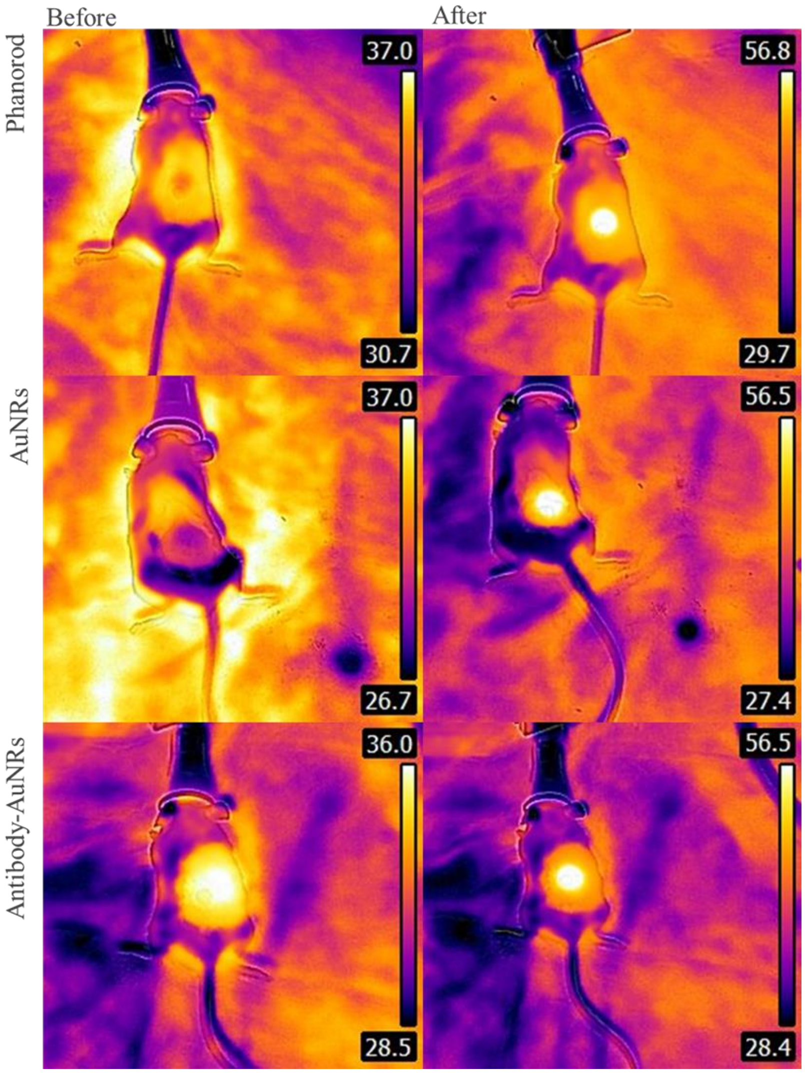


**Figure S18.** Thermal images of phanorod, AuNR (PEG-COOH coated) and antibody-AuNR treatment of infected wounds before and after 808 nm NIR laser irradiation. The laser power density applied to the phanorod group is 0.3 W cm$^{-2}$. For the AuNR and antibody-AuNR treatment groups, the laser flux was adjusted to obtain a wound temperature of 55-56°C.

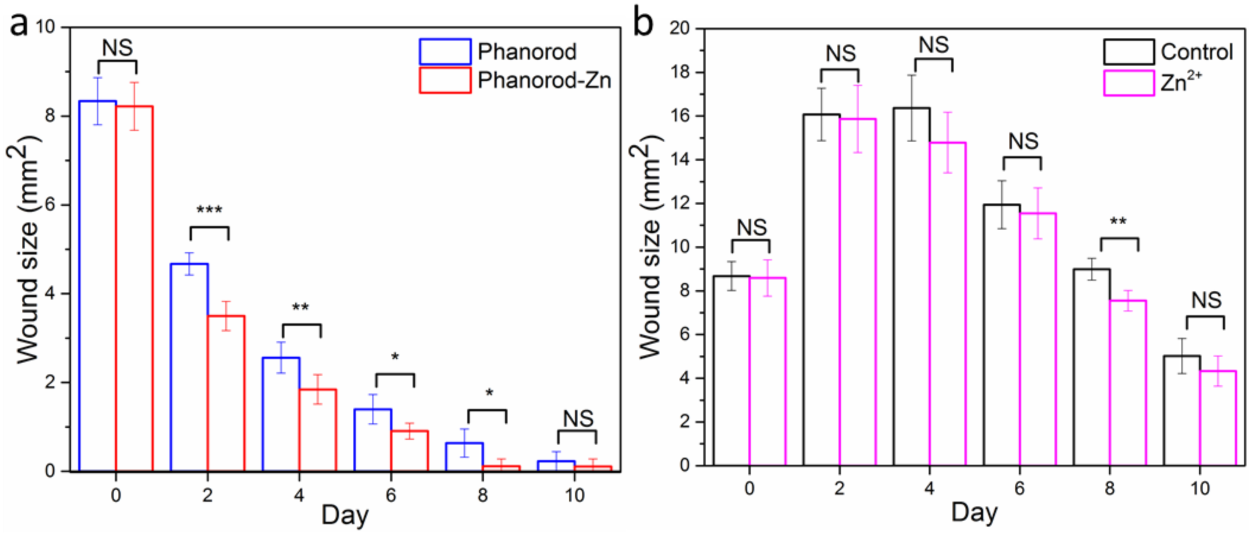


**Figure S19.** The wound area over time for the phanorod and phanorod-Zn treatment, as in Figure 3c in the main text, for phanorod vs. phanorod-Zn (a) and control vs. $Zn^{2+}$ (b). Error bars show standard deviation (n = 5 mice). * indicates $p < 0.05$, ** indicates $p < 0.01$, *** indicates $p < 0.001$, NS = not significant ($p > 0.05$); two-sided *t* test.

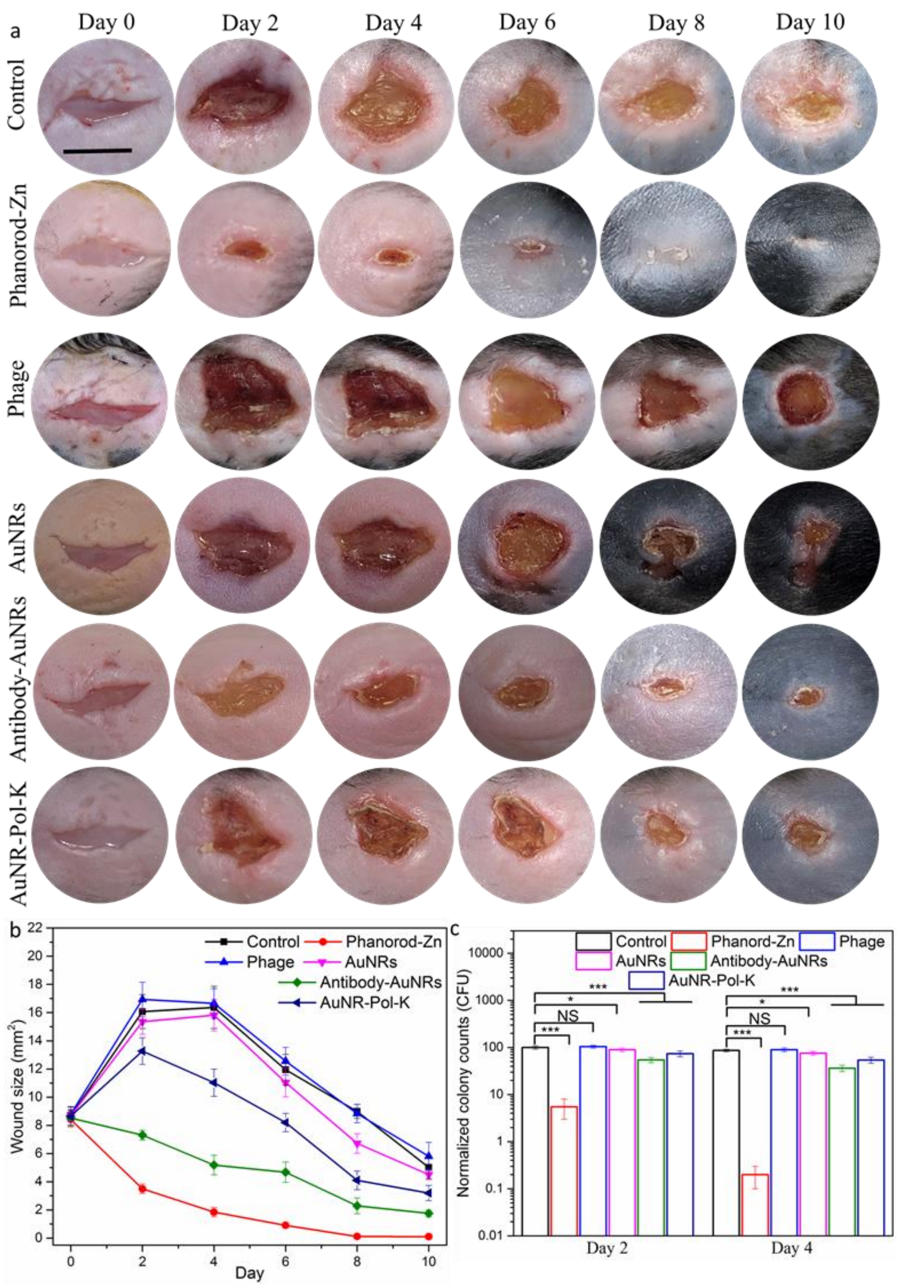


**Figure S20.** (a, b) Representative photographs of the wound healing process and corresponding wound size of *P. aeruginosa*-infected wounds over time (scale bar = 5 mm). The control is treatment with PBS buffer. (c) Quantitative assessment of bacterial load (cfu) in severe wound treatment by colony counting of *P. aeruginosa* on days 2 and 4. Error bars are standard deviation (n = 5 mice). * indicates $p < 0.05$, *** indicates $p < 0.001$, NS = not significant ($p > 0.05$); two-sided $t$ test). Differences between the control and phage groups (b,c) are not significant.

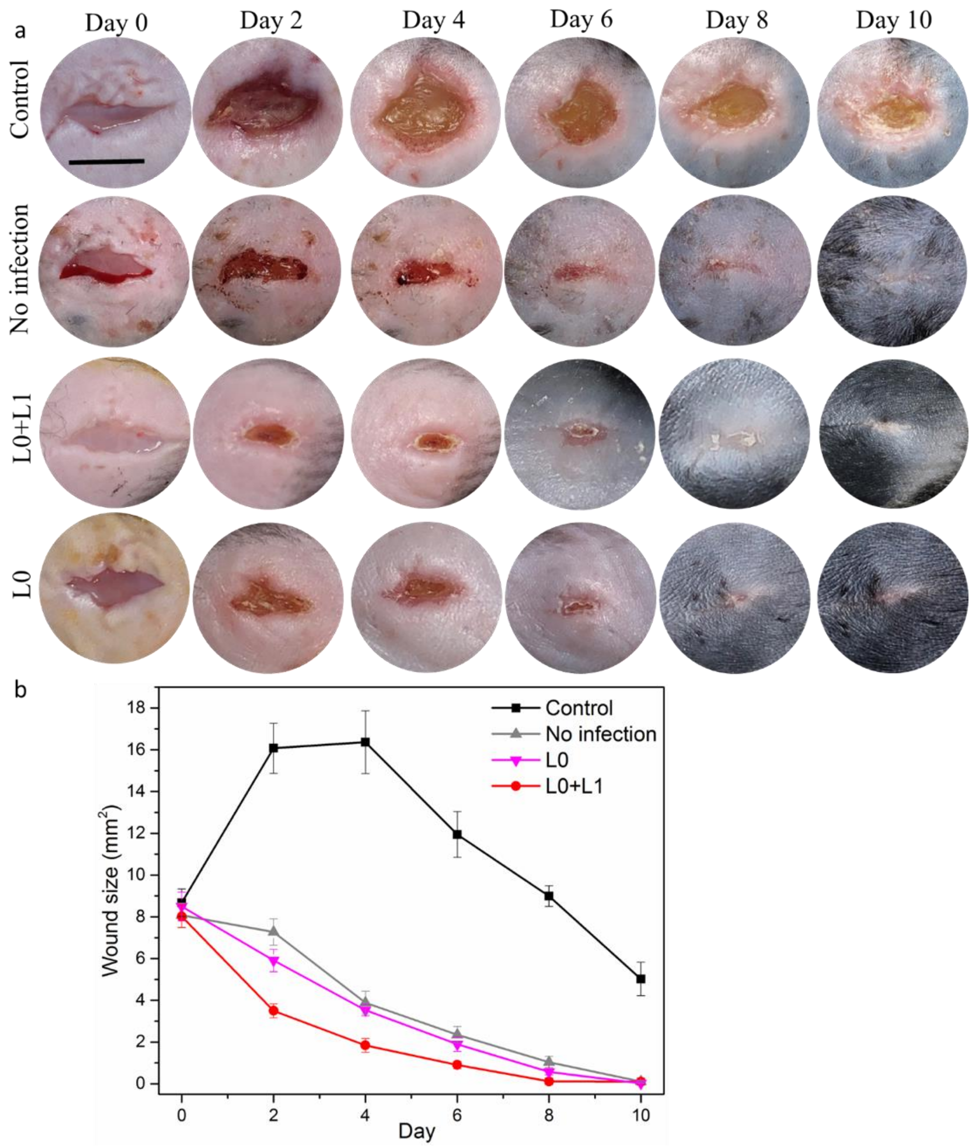


**Figure S21.** Comparison of wound healing for non-infected wounds and shortened phanorod-Zn-treatment for infected wounds. Wound photographs (a) and wound size (b) are shown. Scale bar = 5 mm. The control is treatment with PBS buffer. Phanorod-Zn treatment was performed with laser irradiation on day 0 only (L0; shortened regimen), or day 0 and 1 (L0+L1). Error bars are standard deviation (n = 5 mice).

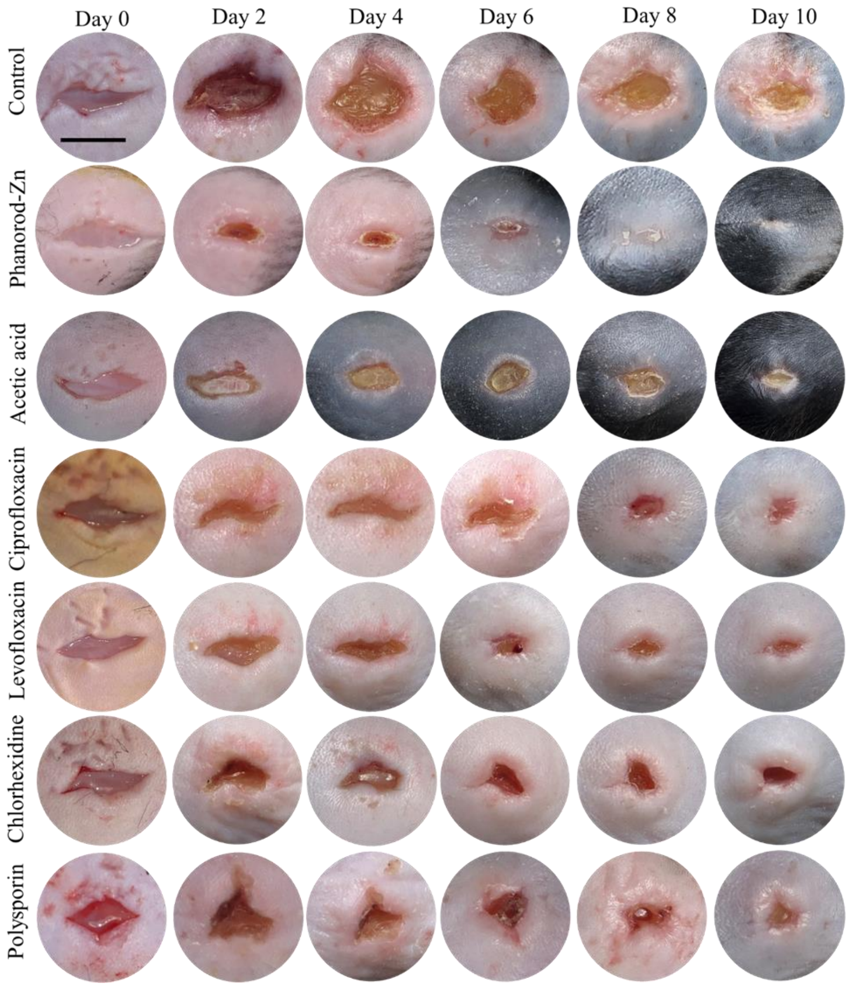


**Figure S22.** Representative photographs of the wounds treated with phanorod-Zn and standard therapies (antibiotics and antiseptics, as listed; scale bar = 5 mm).

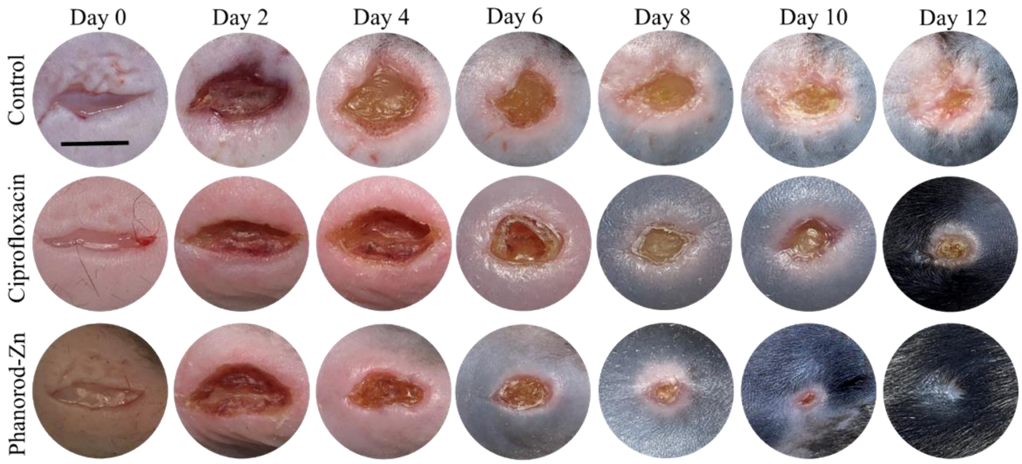


**Figure S23.** Representative photographs of delayed wound treatment by phanorod-Zn or ciprofloxacin. Scale bar = 5 mm. Treatment was initiated on day 2. The control is treatment with PBS buffer.

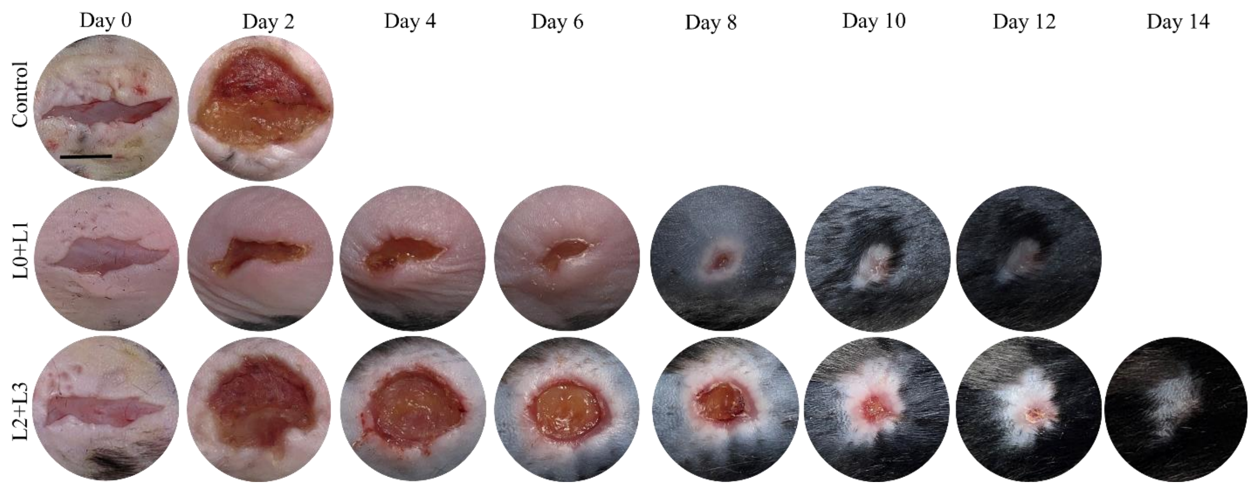


**Figure S24.** Representative photographs of phanorod-Zn treatment of severe wounds over time (scale bar = 5 mm). The control is exposed to PBS buffer. Mice in the control group did not survive after day 2. Treatment was initiated on day 0 (L0+L1) or day 2 (L2 +L3).

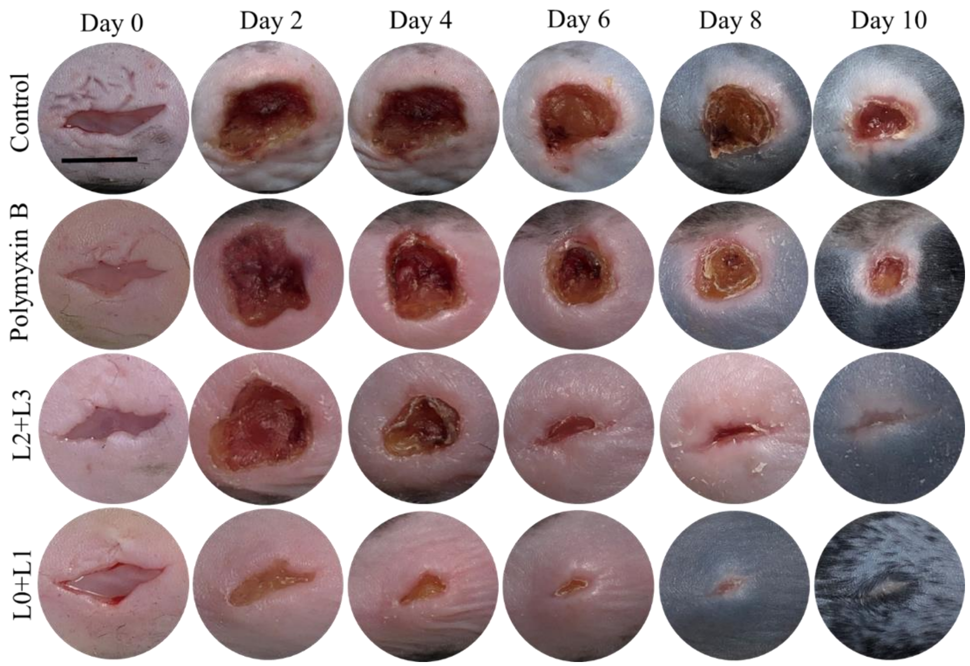


**Figure S25.** Representative photographs of phanorod-Zn treatment or polymyxin treatment of wounds infected by *P. aeruginosa* strain PAK*pmrB6*. Scale bar = 5 mm. The control is exposed to PBS buffer.

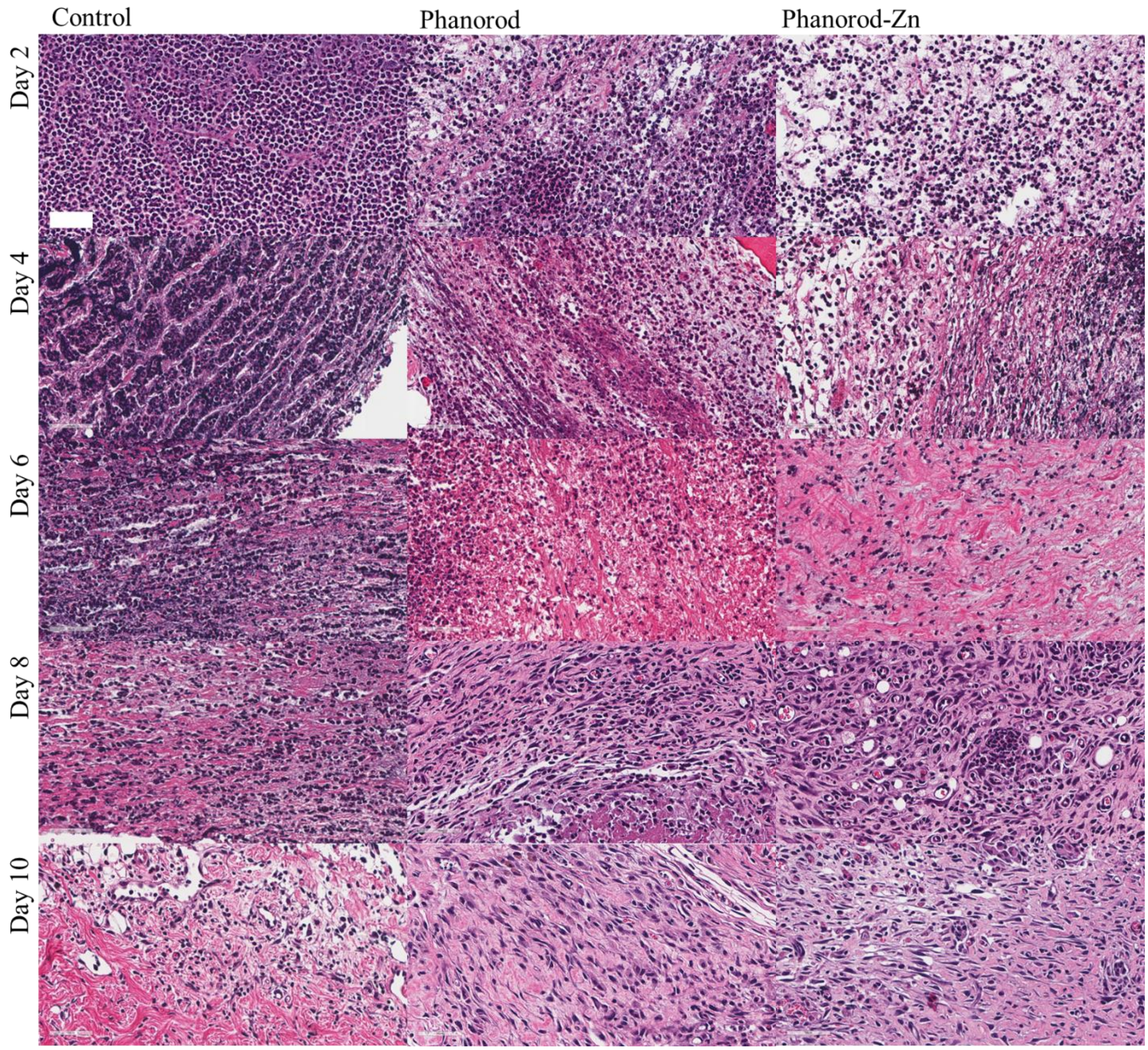


**Figure S26.** Histological analyses of wound tissue through the wound healing process of the control, phanorod and phanorod-Zn groups by H&E staining (scale bar = 50 μm).

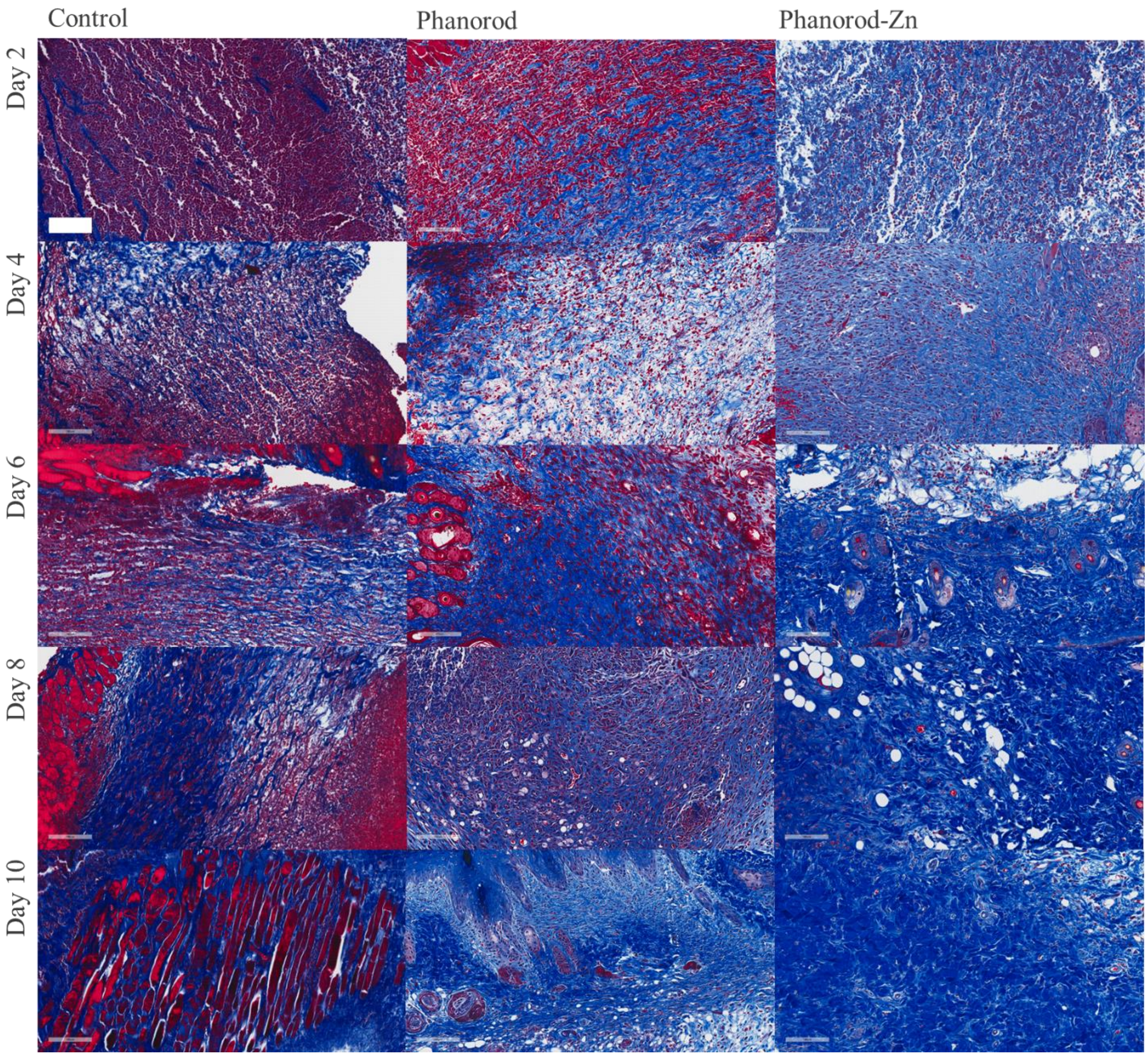


**Figure S27.** Masson's trichrome staining of histological sections of wound tissue on day 2, 4, 6, 8, and 10. First column: control (no treatment); second column: phanorod treatment; third column: phanorod-Zn treatment. Treatments were initiated on day 0. In this stain, collagen appears blue, and cells and other components appear red. Extensive collagen deposition can be observed in the phanorod-Zn treatment. Scale bar = 100 μm.

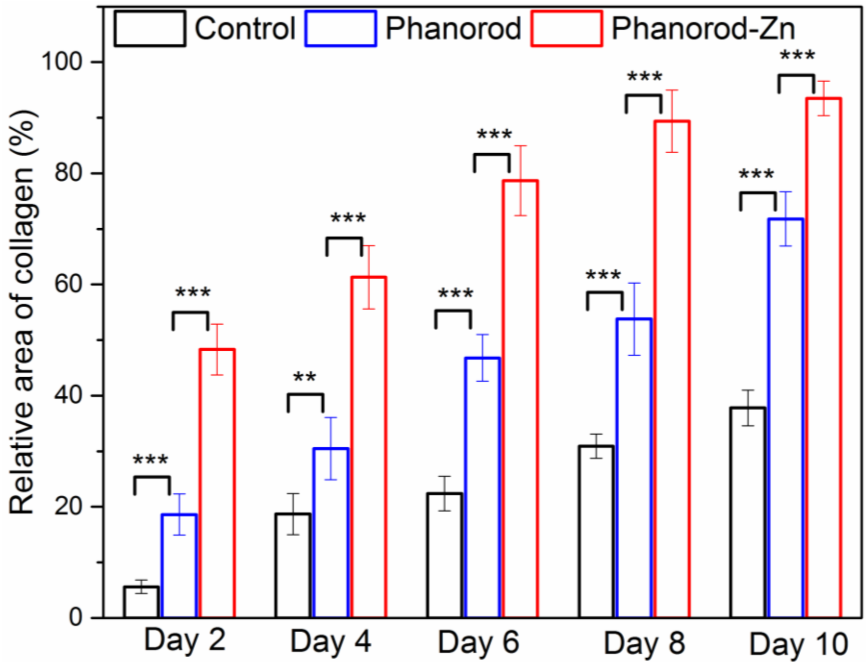


**Figure S28.** Relative area of the deposited collagen in the control, phanorod and phanorod-Zn groups quantified Masson's trichrome staining in Figure S26. Error bars are standard deviation (n = 3 sections). ** indicates $p < 0.01$ and *** indicates $p < 0.001$ (two-sided $t$ test).

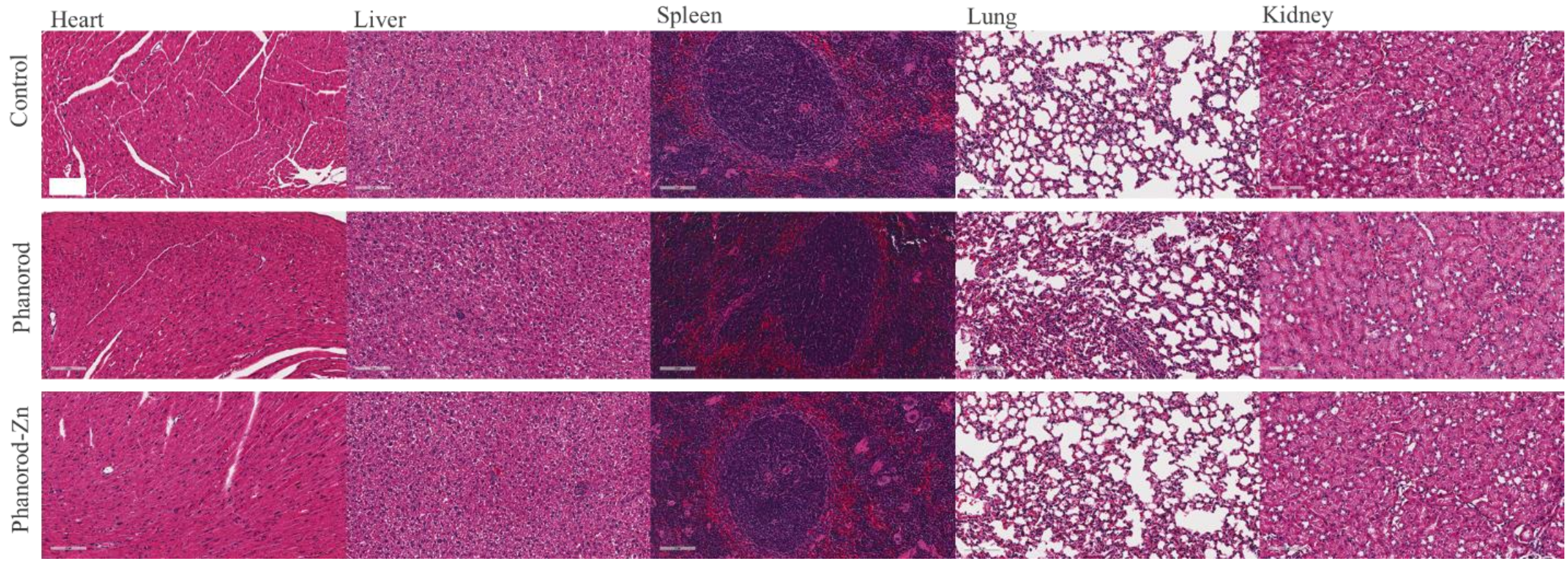


**Figure S29.** Histological analyses of major organs (liver, spleen, kidney, heart, and lung) on day 10 through H&E staining (scale bar = 100 μm).

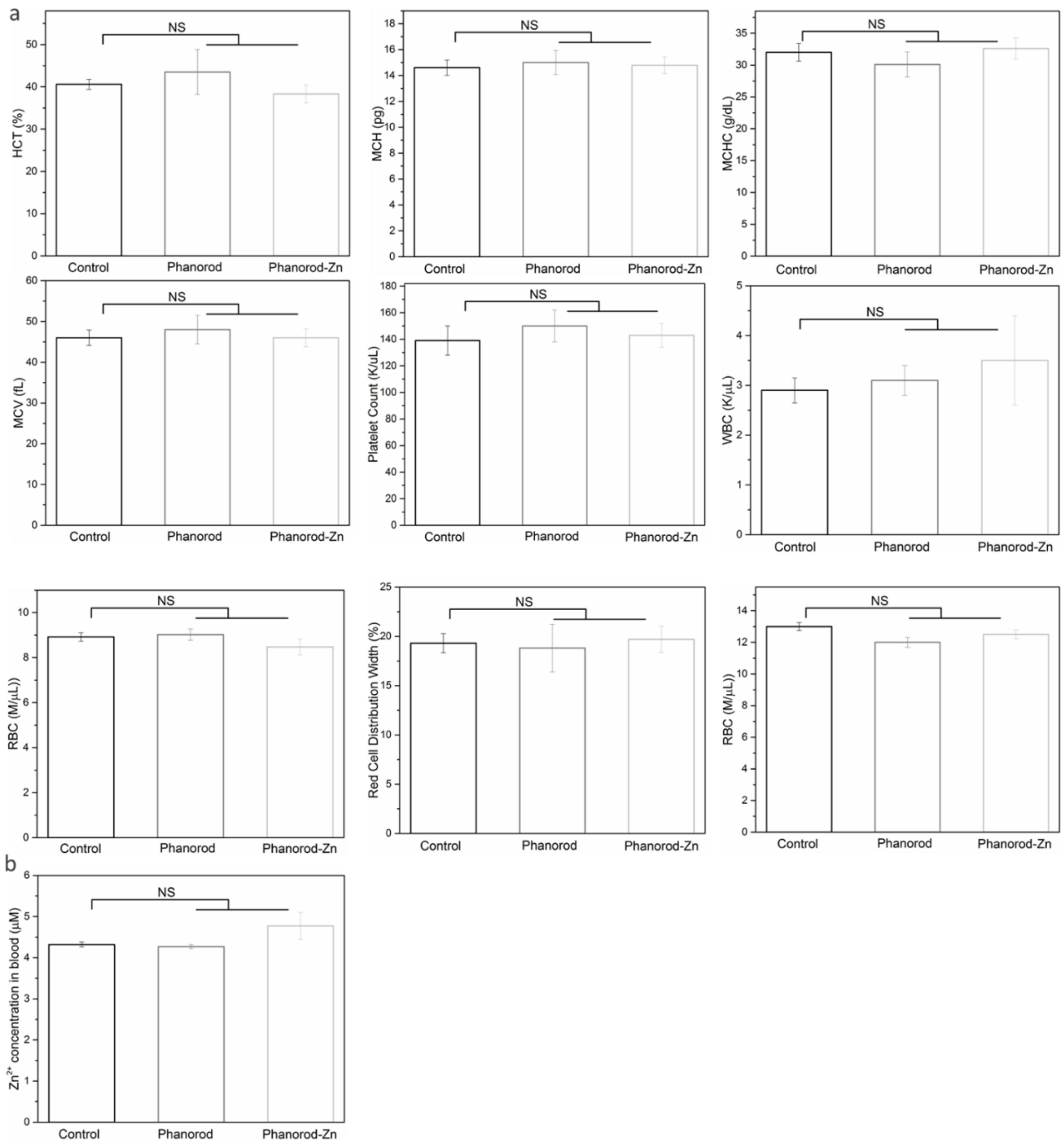


**Figure S30.** (a) Hematological analysis on day 10 of control, phanorod, and phanorod-Zn treatments, showing hematocrit (HCT), mean corpuscular hemoglobin (MCH), mean corpuscular hemoglobin concentration (MCHC), mean corpuscular volume (MCV), platelet count, white blood cell count (WBC), red cell distribution width (RDW) and red blood cell count (RBC). (b) Concentration of $Zn^{2+}$ in blood analysis of control, phanorod and phanorod-Zn groups on day 10. Error bars indicate means ± standard deviations (n = 3). NS = not significant ($p > 0.05$; two-sided Student's $t$ test).

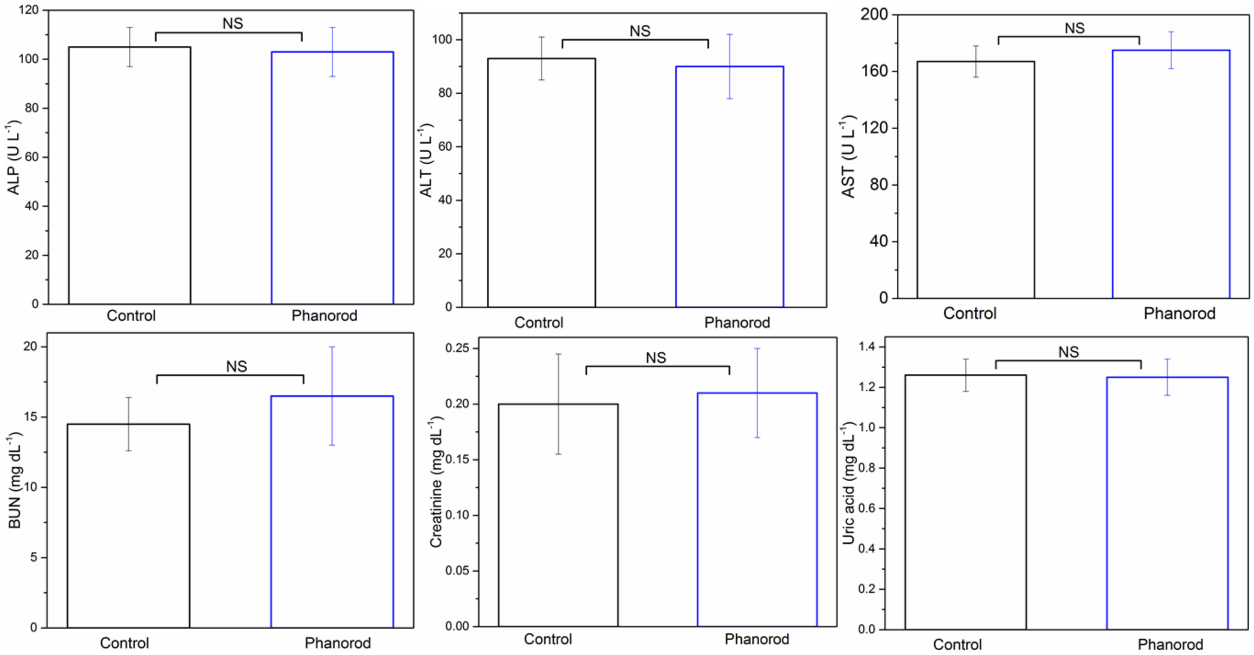


**Figure S31.** Serum biomarkers of hepatic function (ALP, ALT, and AST) and renal function (BUN, CR, and UA) for the control (no treatment), or treatment by phanorods, measured on day 10. Error bars indicate mean ± standard deviation (n = 3). NS = not significant ($p > 0.05$; two-sided $t$ test).

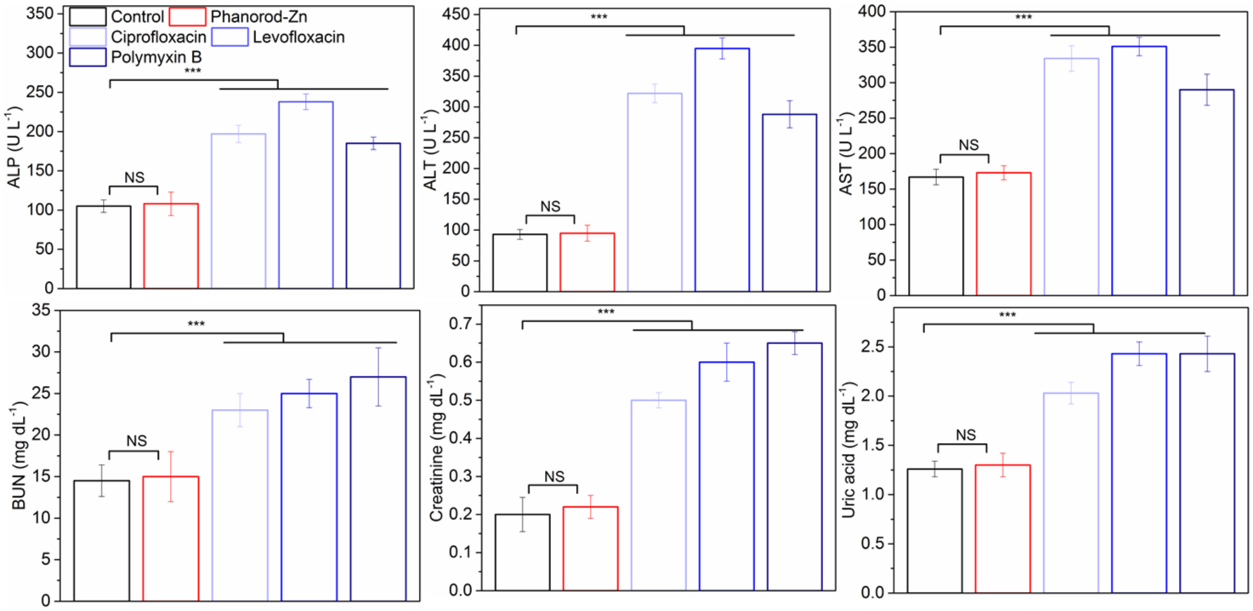


**Figure S32.** Serum biomarkers of hepatic function (ALP, ALT, and AST) and renal function (BUN, CR, and UA) for the control (no treatment), or treatment by phanorod-Zn or antibiotics (ciprofloxacin, levofloxacin, or polymyxin), measured on day 10. Error bars indicate means ± standard deviations (n = 3 mice): *** indicates $p < 0.001$, NS = not significant (two-sided $t$ test).

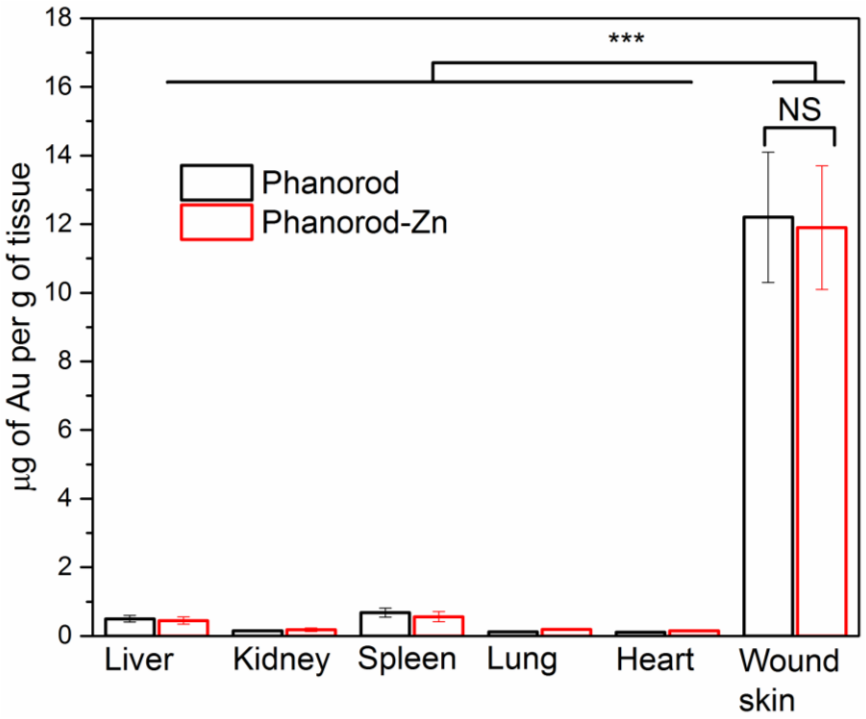


**Figure S33.** Biodistributions of phanorod and phanorod-Zn in different tissues on day 8 measured by ICP-MS for Au. The amount of Au in the control (no phanorods, no phanorod-Zn) was undetectable (<0.002 ng/mL). Error bars indicate mean ± standard deviation ($n$ = 3): *** indicates $p < 0.001$, NS = not significant (two-sided $t$ test).

### Supporting Methods

**Phage preparation.** The construction of the chimeric phage used here, M13-g3p(Pf1)), was previously reported.[6] In brief, the N-terminal domain of g3p of M13KE was replaced by the homolog from the filamentous phage Pf1 (host: *P. aeruginosa*), using a NotI restriction site introduced between the N- and C-terminal domains of g3p. The M13 derivative also encodes a kanamycin resistance gene, allowing maintenance and production of the chimeric phages in *E. coli* under selection. Phages were propagated and quantified by real-time PCR, as previously described[6].

**Propagation of bacteria.** To propagate *E. coli* (ER2738), *V. cholerae* and *P. aeruginosa*, a single colony was inoculated into 5 mL LB medium with tetracycline (10 µg/mL) for *E. coli*, or without antibiotics (for *V. cholerae* and *P. aeruginosa*) in a 50 mL Falcon tube and shaken at 37 °C overnight or for 24 h. For the polymyxin-resistant PAK*pmrB6* strain of *P. aeruginosa*, 4 µg /mL of polymyxin B was added to the growth media. The bacterial cell density was determined by measuring the optical density at 600 nm (OD600). The conversion factor was determined by a colony formation tittering assay as described previously[6]. For *E. coli* ER2738, cell density (CFU/mL) = $4\times 10^8\times$(OD600) + $2.18 \times 10^7$; *V. cholerae*, cell density (CFU/mL) = $8 \times 10^8\times$(OD600); for *P. aeruginosa*, cell density (CFU/mL) = $2 \times 10^8\times$ (OD600) + $4 \times 10^6$.

**Synthesis of gold nanorods (AuNRs).** The gold nanorods were synthesized through a modified method by Murray et al[7]. The seed solution for AuNRs was prepared as follows. Typically, 5 mL $HAuCl_4$ (0.5 mM) was mixed with 5 mL CTAB (0.2 M) solution. 1 mL freshly prepared $NaBH_4$

(0.01 M) was injected into the above solution under vigorous stirring, resulting in a color change from yellow to brownish-yellow. The seed solution was stirred for another 2 min and then aged at room temperature for 35 min before use. To prepare the growth solution, 9.0 g CTAB and 1.1 g 5-BAA were completely dissolved in 250 mL warm water in a 500 mL Erlenmeyer flask by vigorous stirring. The solution was cooled down to 30 °C before 15 mL $AgNO_3$ solution (4 mM) was added. 250 mL $HAuCl_4$ solution (1 mM) was added to the mixture at 30 °C after 15 min. The solution was stirred for another 15 min at 800 rpm, and then 2 mL ascorbic acid (0.064 M) was added. The solution was vigorously stirred for another 2 min resulting in a colorless solution. Finally, 0.4 mL seed solution was injected into the growth solution and stirred for 2 min. The solution was left undisturbed at 30 ° C for 24 h for gold nanorod growth. The AuNRs were isolated, purified by repeated centrifugation and resuspension (in water), and finally redispersed in 15 mL of water.

**Synthesis of AuNR-Pol-K conjugates.** Conjugation of Pol-K peptide to gold nanorods was performed according to previous report[8]. The residual CTAB on the surface of AuNRs were exchanged with HOOC-PEG-SH as described previously[8]. In a typical reaction, 2 mL of PBS buffer containing AuNRs (3.3 μM Au), EDC (1 mM), NHS (1 mM) was incubated overnight at 4 °C. The particles were purified by repeated centrifugation/decantation cycles (8000 rpm for 30 min) and resuspended in 1 mL water. The $Zn^{2+}$ ions were loaded by incubating the AuNR-Pol-K (1 mL of 3.3 μM Au) with 1 mM $ZnCl_2$ solution overnight by gentle rotating. The product was collected after centrifugation (8000 rpm for 30 min) and washing 3 times with water. For *in vitro* bacterial ablation and *in vivo* wound treatment experiments, the applied gold concentration was adjusted to 3.8 μM, with bulk temperature adjusted to 55°C by tuning the laser flux to result in the amount of released $Zn^{2+}$ being similar to the phanorod-Zn group.

**Determination of the binding affinities of M13-g3p(Pf1), phanorod and phanorod-Zn to host cells.** The binding affinity was determined according to previous method[6]. Several tubes of 1 mL of $10^8$ CFU/mL *P. aeruginosa* (Schroeter) Migula (ATCC 25102) cells were incubated with phage or bioconjugate samples of different concentrations, for 10 min at room temperature. The bound cells were separated by centrifugation at 2000g for 10 min and the supernatant was discarded. The pellets were washed extensively with water for three times. The DNA of the bound phages or conjugates was extracted with the PureLink viral RNA/DNA mini kit and quantified by real-time PCR (forward primer: 5'-AAACTGGCAGATGCACGGTT-3'; reverse primer: 5'-AACCCGTCGGATTCTCCG-3'; PCR conditions: 95 °C for 10 min and then 45 cycles of 95 °C for 15 s, 60 °C for 60 s) with SsoAdvanced™ Universal SYBR® Green Supermix and Biorad C1000 PCR machine).

**Determination of infectivity of M13KE-AuNR and M13KE-AuNR-Pol-K**. After 15 mins laser treatment, 5 µL of $10^{14}$/mL M13-AuNRs (or M13KE-AuNR-Pol-K) were incubated with 200 µL overnight culture of *E.coli* ER2738, in 20 mL LB medium in a shaker (250 rpm) at 37°C for 8 hours. The cells were then removed by centrifugation (4500g, 10 min) and the supernatant was transferred to a fresh tube for repeat centrifugation. Avoiding the pellet, 16 mL of the supernatant was transferred to a new tube and mixed well with 4 mL of 2.5 M NaCl/20 % PEG-8000 (w/v). The solution was incubated in 4°C overnight and the putative phage was separated by centrifugation (12000 g, 15 min). The putative viral DNA was extracted with the PureLink viral RNA/DNA mini kit and quantified by real-time PCR (forward primer: 5'-AAACTGGCAGATGCACGGTT-3'; reverse primer: 5'-AACCCGTCGGATTCTCCG-3'; PCR conditions: 95 °C for 10 min and then 45 cycles of 95 °C for 15 s, 60 °C for 60 s) with SsoAdvanced™ Universal SYBR® Green Supermix and Biorad C1000 PCR machine). Wild-type M13KE phage was used as a positive control in the assay.

**Determination of the $Zn^{2+}$-loading capacity of phanorod-Zn.** Briefly, 0.5 mL of phanorod-Zn samples (2x$10^{14}$ phage particles/mL) were incubated with 0.5 mL of 2 mM $ZnCl_2$ solutions overnight at room temperature. The phanorod-Zn were separated by centrifugation at 12000g for 30 min and the supernatant was discarded. The pellets were washed extensively with water for three times. The pellets were digested with aqua regia and 5% $HNO_3$ for a week before ICP-MS measurement to measure the concentration of the bound $Zn^{2+}$. The same experiment was also performed using phanorods to determine the amount of $Zn^{2+}$ bound by phanorods.

***In vitro* cytotoxicity of $Zn^{2+}$, phanorod and phanorod-Zn.** The *in vitro* cytotoxicity was evaluated by the PrestoBlue cell viability assay according to the manufacturer's protocol (Invitrogen). HEK293T cells in suspension were seeded at 5.0 × $10^4$ cells per well in a 96-well microtiter plate and cultured in Dulbecco's modified Eagle's medium (DMEM) (Thermo Fisher; 11885076) supplemented with 10% FBS (Thermo Fisher; 10437036) and 1% penicillin–streptomycin (P/S) (Thermo Fisher; 15140122) in an incubator at 37 °C with 5% $CO_2$. After 48 h, the culture medium was removed, and the cells were washed twice with PBS buffer to remove traces of P/S. Then 180 µL of fresh DMEM (containing 10% FBS) was added with 20 µL of samples of various concentrations of $ZnCl_2$ or the bioconjugates in PBS buffer. The cells were incubated for another 48 h, and the medium was replaced by 90 µL of fresh medium, and mixed with 10 µL of PrestoBlue. The cells were incubated at 37 °C for 1 hour, and fluorescence signals were measured on a TECAN infinite 200 Pro plate reader (TECAN). The excitation wavelength was 560 nm (bandwidth, 9 nm), and the emission wavelength was 600 nm (bandwidth, 20 nm). The cell viability was expressed as a percentage relative to the control cells (HEK293T cells incubated with PBS buffer with no $ZnCl_2$ or bioconjugates, under the same conditions).

**Biodistribution of phanorods and phanorod-Zn.** On day 8, the mice in the phanorod and phanorod-Zn groups were sacrificed. The hearts, lungs, livers, spleens, kidneys and wound skin were collected, weighed, ground, and digested with aqua regia. Then the insoluble precipitates were pelleted by centrifugation (8000 g for 15 mins) and the gold element concentrations in the supernatant were quantified by ICP-MS measurement.
